# Loss-of-function phenomics, ncORFs, and ambiguity of mutant phenotypes in *Medicago truncatula*

**DOI:** 10.64898/2026.03.07.710271

**Authors:** Umut Çakır, Noujoud Gabed, Selen Kaya, Vagner A. Benedito, Marie Brunet, Xavier Roucou, Igor S. Kryvoruchko

## Abstract

Non-canonical open reading frames (ncORFs) are an emerging area of research that is quickly gaining momentum. Many peptides and proteins missed in initial annotation efforts (ncProts) were subsequently shown to be crucial for a wide range of biological processes. The discovery of ncORFs continues to improve the accuracy of loss-of-function studies because they often occupy the same genomic spaces as annotated ORFs. While databases of mutant phenotypes linked to genomic loci exist in a few species, none of these databases integrate the information on ncORFs present in already characterized loci. In this study, we introduce a nearly comprehensive loss-of-function phenomics dataset of *Medicago truncatula* (673 loci characterized over the past 30 years), which was integrated as a new track into the genome browser of this organism. This dataset helped critically analyze the potential contribution of ncORFs to published phenotypes. We detected mass spectrometry (MS)-validated ncORFs in 10 characterized genes, including major regulators of development and symbiotic relationships. We also found conserved ncORFs in 113 characterized genes, including four genes with highly conserved ncORFs. In some studies, the contribution of these ncORFs can be ruled out, while in others it cannot. Using real examples, we systematized ambiguities associated with ncORFs. Furthermore, we highlighted little-known trans effects of insertional mutagenesis on splicing as contributors to that ambiguity. Finally, our meta-analysis of published phenotypes revealed that different protein classes have significantly different (unique) proportions of unconditional, conditional, and neutral phenotypes, potentially reflecting their relative functional importance.

**Significance statement:** This study is the first to merge a nearly comprehensive inventory of loss-of-function studies in a eukaryotic organism with the information on novel MS-validated and conserved ncORFs.

## INTRODUCTION

For a long time, functional genomics and genome annotation efforts have primarily been focused on reference proteins (refProts) translated from canonical (reference) open reading frames (refORFs), often overlooking additional coding potential within the same transcripts (Kozak, 1999). Mass spectrometry (MS) proteomics challenged this paradigm through the discovery of the non-canonical proteome (Vanderperre et al., 2013; Mouilleron et al., 2016; Orr et al., 2020; Cardon et al., 2021; Wright et al., 2022). It was deduced from the abundance of high-quality mass spectra that cannot be assigned to refProts (Ning et al., 2010; Pathan et al., 2017). Such a hidden proteome exists even in the smallest culturable bacterium (*Mycoplasma pneumoniae*) with only 729 ORFs (Lluch-Senar et al., 2016). A considerable part of the non-annotated proteome is thought to originate from translation of non-canonical ORFs (ncORFs) to non-canonical peptides and proteins (ncProts). The recognition of the biological relevance of these ncProts is relatively recent (Mouilleron et al., 2016; Orr et al., 2020; Mudge et al., 2022) and has been accelerating in the past few years (Ardern, 2023; Álvarez-Urdiola and Riechmann, 2025; Çakır et al., 2025b; Cardon et al., 2025; Pavesi, 2025; Chang et al., 2025).

The ultimate goal of functional genomics studies is to establish accurate relationships between phenotypes and genotypes for the benefit of biomedical research, agriculture, and biotechnology. We can refer to the multitude of such phenotype-genotype units as a phenome. Deducing a true phenome from functional data can be very complex due to the polycistronic nature of eukaryotic transcripts (Mouilleron et al., 2016). Specifically, it introduces a new level of complexity to the interpretation of all functional genomics studies. A ncORF with a biological function can be present on the same RNA molecule together with the refORF. For example, it can be located in a sequence annotated as an untranslated region (UTR) based on the refORF, even though this region may contain a functional coding sequence in any reading frame. For such transcripts, loss-of-function approaches based on post-transcriptional gene silencing, such as RNA interference (RNAi) (Sharp & Zamore, 2000; Baulcombe, 2004; Chaudhary et al., 2024), yield phenotypes that cannot be unequivocally attributed to refProts alone because an RNAi construct simultaneously affects both refProts and ncProts. Insertion-deletion mutagenesis systems such as CRISPR/Cas9 (for Clustered Regularly Interspaced Short Palindromic Repeats [CRISPR] and CRISPR-associated protein 9 [Cas9]; Doudna & Charpentier, 2014; Guo et al., 2023) and retrotransposons (e.g., *Tnt1*, for Transposon of *Nicotiana tabacum*; Tadege et al., 2008; Lee et al., 2018; Kaur et al., 2021) may seem to be more proof against such ambiguity because they can be more selective regarding the target region of a transcript. However, these systems, too, can affect both a refProt and a ncProt, especially if their ORFs overlap and share the same genetic space. In this study, we described a scenario in which insertional mutagenesis affects both ORFs, even if the insertion is located only in a refORF, acting on a ncORF in trans through the effect on splicing (“in trans” because a sequence beyond the immediate insertion site is affected; see Supporting Discussion). Likewise, gain-of-function studies introduce similar ambiguity because an overexpression of a refORF simultaneously overexpresses an embedded ncORF. These considerations call for an objective re-evaluation of published phenotypes in light of the potential contribution of ncORFs.

Although very comprehensive phenomics databases have been established for mouse (Wang et al., 2015; Baldarelli et al., 2024), *Arabidopsis thaliana* (Lloyd and Meinke, 2012), and other eukaryotes (Courtier-Orgogozo et al., 2020), none of them considered the contribution of ncORFs so far. Our analysis was conducted using *Medicago truncatula* because it stands out as a model organism. In contrast to *Arabidopsis*, it can establish symbiotic nitrogen fixation (SNF) in association with soil bacteria (rhizobia) and arbuscular mycorrhizal (AM) symbiosis with fungi (Barker et al., 1990; Kang et al., 2016; Bandyopadhyay et al., 2019; Nandety et al., 2022). *M. truncatula* has been used as a model for functional genomics since 1995 (Sagan et al., 1995). Despite 30 years in this status, the model has no comprehensive inventory of loss-of-function studies so far. Likewise, a large-scale detection of ncORFs on annotated transcripts had not been conducted in *M. truncatula* until very recently (Çakır et al., 2025b). Even though much attention was paid to the inventory of bioactive peptides in *M. truncatula* (Mergaert et al., 2003; Zhang et al., 2006; Volkening et al., 2012; de Bang et al., 2017; Kereszt et al., 2018; Boschiero et al., 2020), most of them should only conditionally be categorized as ncProts because they originate from transcripts with no annotated long refORFs. Their short ORFs act as refORFs of corresponding transcripts (see supporting data, for example, Data S12).

In this study, we filled the knowledge gap about ncORFs, primarily mRNA-ncORFs, in conjunction with published loss-of-function phenotypes attributed to refORFs. We collected information on nearly all such phenotypes linked to specific loci or groups of genes and merged it with data on MS-validated and conserved ncORFs. For the MS validation, we analyzed three large datasets available on ProteomeXchange (Deutsch et al., 2023). They correspond to 16 samples of 10 different organs (PXD002692, Marx et al., 2016; PXD013606, Shin et al., 2021; and PXD022278, Castañeda et al., 2021). To identify conserved ncORFs, we conducted a three-frame global amino acid sequence similarity search of the whole transcriptome of *M. truncatula* using UniProt v. 2020_02 as a subject database (UniProt Consortium, 2019). We deployed a simple, “naïve” approach based on the difference in the number of hits between ncORFs (conservation at the nucleotide level) and corresponding ncProts (conservation at the amino acid level). This way, we detected highly conserved ncORFs under purifying selection in four well-characterized genes, including the master regulator of phosphate homeostasis *MtPHO2-A* (Curtin et al., 2017; Čermák et al., 2017; Huertas et al., 2023). We exemplified the applicability of this “naïve” approach for the large-scale detection of ncORFs that undergo purifying selection. Our analysis permitted discrimination between studies that can rule out the contribution of ncORFs to reported phenotypes and studies in which such ambiguity calls for additional experiments, such as differential mutagenesis of ncORFs and refORFs. By straightforward sequence comparisons, we explained the likely origin of highly conserved ncORFs in five well-characterized genes and showed that strong conservation may not necessarily imply translation of a given ncORF. Instead, some conserved ncORFs are better understood in the context of alternative splicing or gene model structure. Thus, our approach is not only informative for identifying potentially functional ncORFs, but also valuable for refining current gene annotations. Due to space limitations, we placed much of the valuable information about MS-validated and conserved ncORFs into files called Supporting Results and Supporting Discussion. We refer readers to these files for the deep coverage of ncORFs in transcripts characterized by loss-of-function approaches.

Our phenomics resource will be useful to the *M. truncatula* community regardless of the ncORF dimension, because it serves the following basic purposes: (1) avoiding the duplication of efforts in studying genes that have already been targeted by other groups; (2) consistency of naming new loci and avoiding redundancy; (3) correction of gene models. It should be noted that despite the very impressive accuracy of the current genome annotation (v. 5.1.9, Pecrix et al., 2018), we found surprisingly large number of cases in which annotation accuracy can be further improved.

Lastly, our inferential meta-analysis of loss-of-function phenotypes indicates that diverse protein classes have unique proportions of altered (non-conditional), conditional, and neutral phenotypes, which are significantly different from the overall and specific proportions of functional protein groups in the dataset. These distinct frequencies of phenotypes may reflect the degree of biological importance or functional redundancy of specific groups of proteins. This information can help navigate searches to genes with strong or weak but clear phenotypes by focusing on specific protein classes with high proportions of non-conditional phenotypes. Likewise, it may help prevent misallocation of time and resources in functional genomics or breeding efforts when the likelihood of a given gene to produce a notable phenotype is low. For this reason, our resource is also relevant for applied research and industry, because it can be mined for trait-associated genes, including those involved in symbiosis. This type of information is not available even in the most comprehensive plant phenomics database developed for *A. thaliana* (Lloyd and Meinke, 2012).

## RESULTS

### Structure, scope, and general features of the loss-of-function phenomics dataset

Year 2025 marked the 30^th^ anniversary of *M. truncatula* as a model for loss-of-function genomics (Sagan et al., 1995). We conducted a very detailed human curation of 565 loss-of-function studies published between 1995 and 2025. We adopted the style and structure of the first large list of loss-of-function phenotypes of *M. truncatula*, which was compiled by Roy et al. (2020). The list contained 115 loci with SNF phenotypes. We extended the list to 380 loci with SNF phenotypes and 293 loci with non-SNF phenotypes, 673 loci in total (Figure 1). Additional reputable resources contributed to the compilation of our non-SNF list. Especially useful were the review by Kang et al. (2016) and multiple chapters in two books: Cañas and Beltrán (2018) and Sinharoy et al. (2022). Here, it is important to mention a recent review article by Ye et al. (2024). Although we have not used it as a navigator to the original loss-of-function studies, it provides a very comprehensive coverage of functional genomics in *M. truncatula*, which is complementary to our discussion.

**Figure 1.**
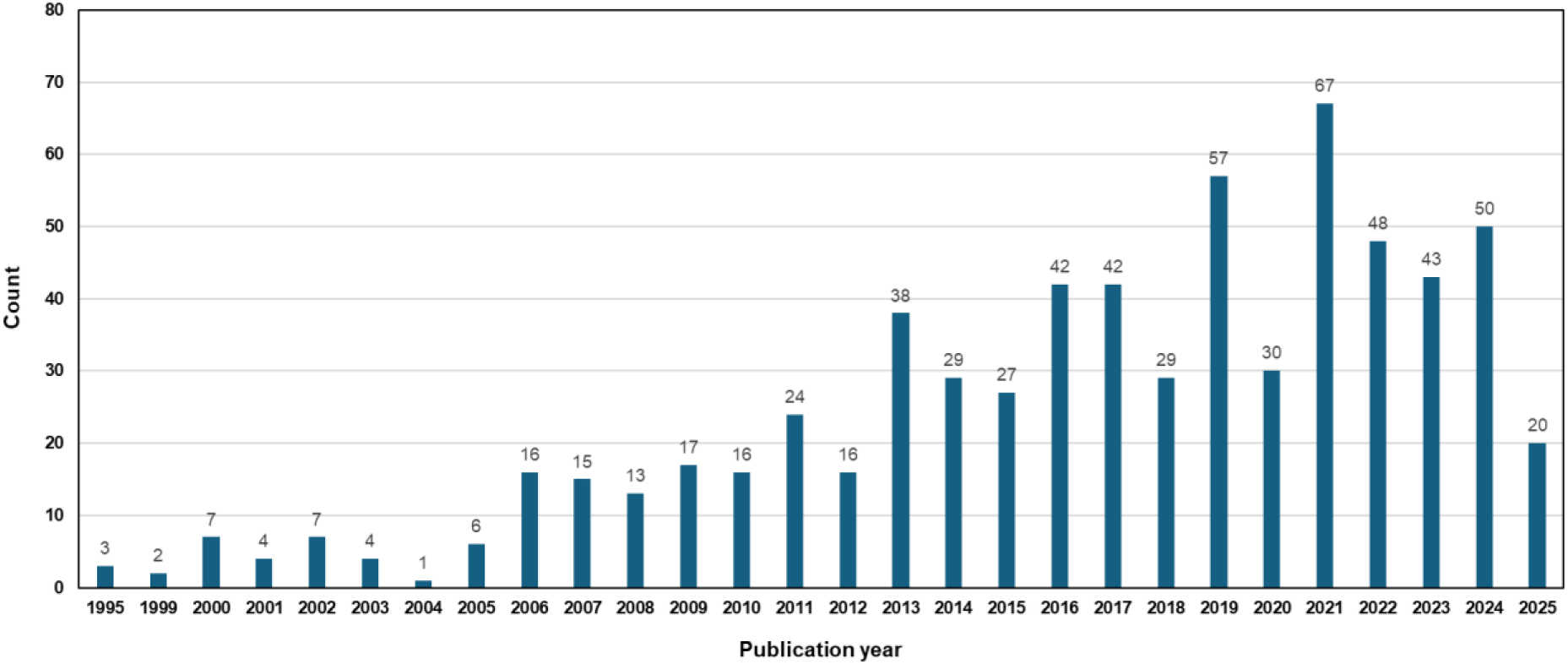
Timeline of loss-of-function studies in Medicago truncatula. The graph is based on the first publication of each genetic locus, 673 loci in total. 219 of these genes were studied in the context of both symbiotic nitrogen fixation (SNF) and other processes. 380 loci were tested for SNF phenotypes and 512 for non-SNF phenotypes. On average, ca. 47 new genes have been characterized each year since 2020 (2025 excluded). The complete dataset consists of 565 published works and bioRxiv preprints as of 16 June 2025.

Since *M. truncatula* is primarily a model for studies on SNF, we kept SNF-based lists separately and contrasted them with non-SNF lists in some datasets. While importing the information about the 115 loci from Supplemental Dataset 1 of Roy et al. (2020), we rechecked each phenotype using an approach centered on *M. truncatula*, which resulted in some minor differences between our dataset and the dataset of Roy et al. (2020). We added a column with the current locus identifiers (genome annotation v. 5.1.9). Deducing this information was often difficult because not all studies provided an easy means of identifying the targeted genes. We had to conduct careful analysis of primer sequences, RNAi constructs, *Tnt1* flanking sequence tags (FSTs), and other data to find the exact correspondence of studied loci with gene models of the current genome release. Despite these efforts, it was not possible to ascertain the locus identifiers of seven characterized genes: *MtCIR2*, *MtCRA1*, *MtGLYI*, *MtLAL*, *MtLHCB1.4*, *MtLYK2bis*, and *MtLYK5bis*. Three of them are likely to exist only in the ecotype R108: *MtGLYI*, *MtLYK2bis*, and *MtLYK5bis*. The sequence of *MtCIR2* was kindly provided by the authors (Zhao et al., 2023). However, it has no corresponding gene model in the latest release of the genome annotation. Locus *MtuORF1p* is a special case. It is an upstream ORF (uORF) located in the 5’-UTR of a characterized transcription factor *MtNF-YA1/MtHAP2-1* (Combier et al., 2006, 2008). These eight genetic units, specifically, *MtCIR2*, *MtCRA1*, *MtGLYI*, *MtLAL*, *MtLHCB1.4*, *MtLYK2bis*, *MtLYK5bis*, and the uORF *MtuORF1p* found in the transcript of *MtNF-YA1/MtHAP2-1*, were excluded from the count of unique locus identifiers (673 unique loci listed but 681 cistrons described; uORF MtuORF1p and MtNF-YA1/MtHAP2-1 are two different cistrons of a single locus). Consequently, these eight loci are also excluded from all figures and most datasets. Details of corresponding phenotypes of these and other genes can be found in Data S1-S6.

It should be noted that studies describing mutants with no linked genetic loci were beyond the scope of our analysis. Thus, the few mutants mentioned above (for example, mutants of *MtCIR2* and *MtCRA1*) are an exception because their corresponding loci were thought to be known. Data S1 (non-SNF refORFs) and Data S2 (non-SNF ncORFs) contain information on refORFs and ncORFs associated with non-SNF phenotypes, respectively. Data S3 (SNF refORFs) and Data S4 (SNF ncORFs) provide the corresponding information for SNF phenotypes. Data S5 lists genes studied both for SNF and non-SNF phenotypes. Data S6 combines the information on all 681 genes studied by loss-of-function approaches. Data S7 contains references for loss-of-function studies. Data S8 is reserved for references describing gain-of-function phenotypes of ncORFs. Our core dataset consists of six Excel files described in Table 1. Although our study focused on loss-of-function phenotypes, an exception was made for characterized ncORFs. To provide a deeper functional description of ncORFs, we also included gain-of-function phenotypes in Data S2 and S4. However, all other datasets in our study are based exclusively on loss-of-function phenotypes. The complete list of datasets pertinent to this study is shown in Table 2 (Data S7-S19). Data S10 and S11 are of special importance because they can be used as “cheat-sheets” by functional genomics specialists and researchers in the industry. Specifically, the lists can predict the likelihood of a gene giving a scorable phenotype. Data S12 is a master table listing and analyzing characterized loci that have MS-validated and/or conserved ncORFs.

**Table 1.** Description of the core datasets.

| Dataset | Locus type | Phenotypes | Type of functional characterization | Number of loci<sup>†</sup>, LOF | Number of loci<sup>†</sup>, GOF only |
| --- | --- | --- | --- | --- | --- |
| Data S1 | RefORFs | Non-SNF | LOF | 508 (501) | - |
| Data S2 | NcORFs | Non-SNF | LOF, GOF | 11 (11) | 16 (15) |
| Data S3 | RefORFs | SNF | LOF | 354 (351) | - |
| Data S4 | NcORFs | SNF | LOF, GOF | 30 (30) | 18 (17) |
| Data S5<sup>‡</sup> | RefORFs and NcORFs | Non-SNF and SNF | LOF | 222 (219) | - |
| Data S6<sup>§</sup> | RefORFs and NcORFs | Non-SNF and SNF | LOF | 681 (673) | - |

<sup>†</sup>Numbers in brackets correspond to unique loci for which the identifiers in v5.1.9 are known.
<sup>‡</sup>Data S5 lists only those genes that were tested for both non-SNF and SNF phenotypes.
<sup>§</sup>Data S6 lists all genes.
Abbreviations:
RefORFs – reference open reading frames
NcORFs – non-canonical open reading frames
LOF – loss-of-function phenotypes
GOF – gain-of-function phenotypes
SNF – symbiotic nitrogen fixation phenotypes

**Table 2.** Description of the supporting datasets.

| Dataset | Description |
| --- | --- |
| Data S7 | References for LOF studies only (refORFs and ncORFs) |
| Data S8 | References for GOF studies only (ncORFs only) |
| Data S9 | The complete list of 97 biological processes examined in LOF studies |
| Data S10 | The complete list of protein classes, corresponding phenotype counts, and statistical analysis of proportions |
| Data S11 | The complete list of combined protein groups, corresponding phenotype counts, and statistical analysis of proportions |
| Data S12 | A master table that summarizes various parameters of 148 ncORFs found in 123 genes targeted by functional analyses |
| Data S13 | Visual demonstration of conservation at the amino acid level for eight selected ncORFs |
| Data S14 | Evidence for purifying selection of the eight selected ncORFs |
| Data S15 | 24 nucleotide alignments that demonstrate evidence for purifying selection of the eight selected ncORFs |
| Data S16 | Sequence analysis demonstrating the origin of highly conserved ncORFs in five selected genes |
| Data S17 | Nucleotide alignments used in Data S16 |
| Data S18 | Gene models and mapped ncORFs of 123 loci targeted by functional analyses in *M. truncatula* (Data S12) |
| Data S19 | NcProts grouped with refProts during the MS analysis |

**Table 3.** Loci incorrectly named in the genome browser v. 5.1.9.

| Name<sup>†</sup> | Actual locus ID (publication) | Publication |
| --- | --- | --- |
| *MtCbf4* | MtrunA17_Chr1g0203861 | Zhang et al., 2016b |
| *MtChOMT1*<sup>‡</sup> | MtrunA17_Chr1g0200491 | Wu et al., 2024b |
| *MtGA20ox1* | MtrunA17_Chr1g0204181 | Li et al., 2021b |
| *MtHDT3* | MtrunA17_Chr7g0265761 | Li et al., 2021a |
| *MtMYC2* | MtrunA17_Chr8g0366751 | Guo et al., 2024 |
| *MtPRP1*<sup>‡</sup> | MtrunA17_Chr4g0013391 | Erickson et al., 2020 |
| *MtPRR7* | MtrunA17_Chr1g0181811 | Wang et al., 2023 |
| *MtRac1* | MtrunA17_Chr6g0486041 | Wang et al., 2021 |
| *MtROP9* | MtrunA17_Chr5g0405791 | Kiirika et al., 2012 |
| *MtSCR* | MtrunA17_Chr7g0245601 | Dong et al., 2021 |
| *MtSLB1* | MtrunA17_Chr5g0447381 | Yin et al., 2020; Zhou et al., 2021 |

<sup>†</sup>The table lists correct names and corresponding locus identifiers. These names were incorrectly assigned to other loci in the genome browser v. 5.1.9.
<sup>‡</sup>Names *MtChOMT1* and *MtPRP1* were misassigned to two genes each in the genome browser v. 5.1.9.
Table S1 provides an extended version of this list.

### The critical need for a comprehensive loss-of-function phenomics dataset in *M. truncatula*

The progress of functional genomics in *M. truncatula* has been greatly accelerated by the insertional mutagenesis using tobacco retrotransposon *Tnt1* (Tadege et al., 2008; Lee et al., 2018; Kaur et al., 2021), although the toolkit is not limited to this system (Grandbastien et al., 1989; Mazier et al., 2007; Cui et al., 2013; Vives et al., 2016). Over the past five years, on average, nearly 47 new genes have been characterized by loss-of-function approaches each year (Figure 1). With the advent of CRISPR/Cas9 (Doudna & Charpentier, 2014; Guo et al., 2023), this number will hopefully rise further each year. The total number of genes with information on mutant phenotypes reached 673 by 16 June 2025, which is only a fraction of the corresponding number of genes characterized in *Arabidopsis* (Lloyd and Meinke, 2012). Nevertheless, 673 is the number large enough to require systematic recording, curation, and meta-analysis.

The lack of a standardized protocol for collecting and displaying information on newly characterized genes compromised the status of *M. truncatula* as a functionally annotated model. When a researcher identifies a gene that looks interesting for the functional analysis, one of the first questions is whether it has already been studied. Unfortunately, this information is not readily available for 60% of genes out of 673 characterized so far (Figure 2a; Data S6). In this context, it is important to emphasize the central role of the *M. truncatula* genome browser v. 5 developed in 2018 (Pecrix et al., 2018). This indispensable resource is the hub linking many types of data and displaying them in a well-organized, user-friendly format for the maximal benefit of the *Medicago* community. However, integrating old and new functional genomics data into the genome browser is a major technical challenge. This is because the truly comprehensive and error-free process of collecting such data cannot be easily automated. For example, it is very unlikely to be efficiently outsourced to artificial intelligence. The team curating the browser made an enormous effort and accurately referenced loss-of-function studies for 269 genes, which is 40% of the total (Figure 2a; Data S6, “Yes” in Columns V and W). Unfortunately, 173 genes (26%) completely evaded the functional annotation (Figure 2b; Data S6, “No” in Columns V and W), while 218 genes (32%) were provided with references that do not inform the users about the corresponding loss-of-function analyses (Figure 2a, categories “Yes/YN” and “No/YN”; Data S6, “NY” in Column W). Naturally, newer studies tend to be missed at a higher rate compared to older ones. For example, more than half of genes functionally characterized since 2023 are not labeled as such by the end of July 2025 (Data S6 Column W). However, the population of 173 “missed” genes (“No/No” category in Figure 2a) extends in time back to 2001, with 81 genes characterized more than five years ago (Figure 2b; Data S6). This lack of integration into the browser, even years after the original publication, reflects the difficulty of manual curation of such data. We found no bias toward any method, phenotype, or protein class among those “missed” genes. They were overlooked simply due to the lack of a centralized system integrating all functional data into the browser.

**Figure 2.**
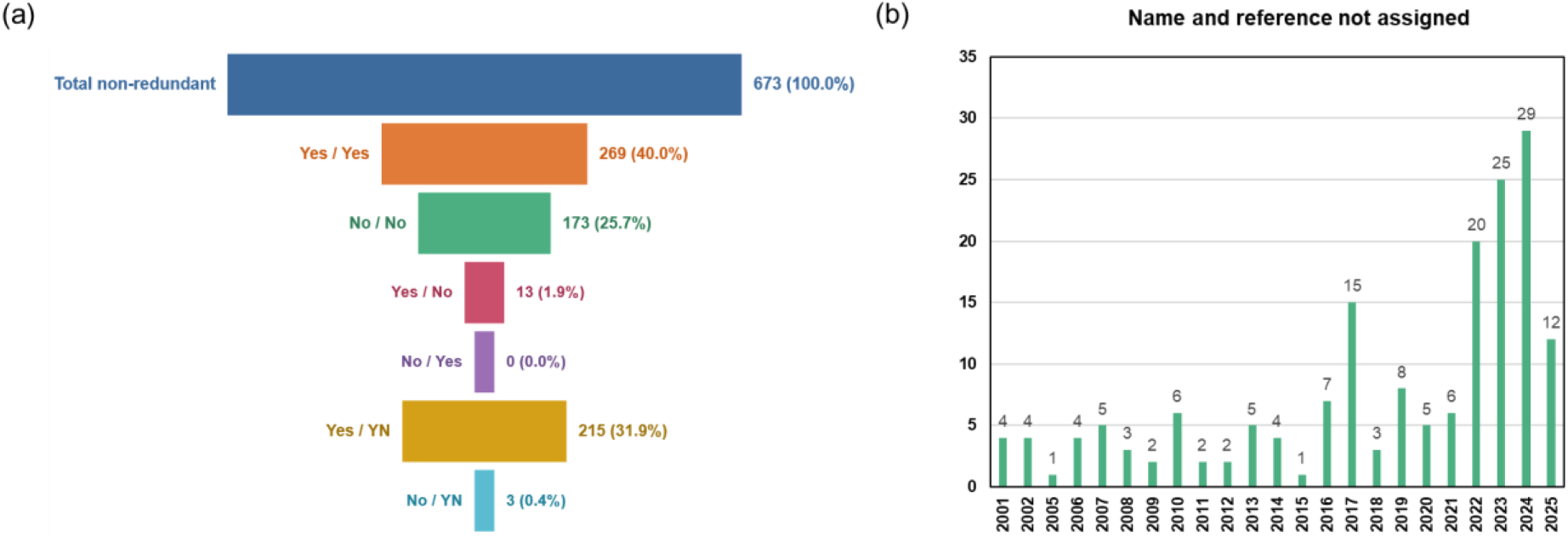
Functional annotation status of 673 genes studied by loss-of-function approaches in *M. truncatula*, based on the genome browser v. 5.1.9. (a) Six categories of loci according to their annotation status. The first element in each category’s code corresponds to a gene name (“Yes” - name assigned, “No” - name not assigned). More than one quarter (26%) of characterized genes were not named in the browser (Data S6 Column V). The second element in each category’s code corresponds to a reference (“Yes” - reference assigned, “No” - reference not assigned, “YN” for “Yes and No” - an irrelevant reference assigned). More than one quarter (27%) of characterized genes had no reference (Data S6 Column W). Among genes with a reference, 32% had only irrelevant references or references to transcriptomic studies/reviews instead of correct references to loss-of-function studies. (b) The distribution of 173 “missed” genes over their publication years (category “No/No”, no name and no reference in the genome browser v. 5.1.9). Only the year of the first loss-of-function publication is counted for each gene.

These observations strongly suggest the need for such a system. Possibly, it could be developed with a Wikipedia-like format, with authors encouraged or obligatorily requested to enter all relevant information about genes at the time it is submitted for publication. One reason for such an open format is the difficulty of allocating this task to a specific researcher/curating team member. Because dozens of new genes are characterized each year, keeping functional data up to date requires significant and continuous effort. For example, for our team, it took nearly three years to collect and systematize information about 673 genes. This problem can be efficiently addressed by agreeing upon a set of rules and implementing these rules for all researchers who publish functional studies on *M. truncatula* genes. Our dataset can be the basis for such a system, which can be further developed in real time as more genes are characterized each year.

In the absence of such a system, the corresponding author will continue the collection, analysis, and integration of loss-of-function data into the separate track opened in the genome browser of *M. truncatula* on 10 April 2026 (see Data Availability).

### Specific examples where a unified loss-of-function database could help avoid redundancy and confusion

An important question is whether this lack of functional annotation had any negative impact on studies in *M. truncatula*. Can we find examples of such impact through the meta-analysis of our dataset? Unfortunately, yes. Column X (Notes) in Data S6 describes a few such instances. They are categorized below.

### Multiple names for the same gene: potential duplication of efforts

The first loss-of-function study on locus MtrunA17_Chr5g0422541 (*MtAOC1*) was published in 2005 (Isayenkov et al., 2005). Recently, a different team characterized this gene very completely, too, and named it *MtAOC2* (Yang et al., 2023). Both studies were published in Plant Physiology and covered different aspects. The second team, and the editor of the journal, were unaware of the 18-year-old study on this gene. It is evident that no researcher or editor would ever wish to miss such important information by the time of the intended publication, or ideally much earlier. Had it been available through a centralized resource, it would greatly facilitate planning, experimental design, and manuscript writing for the second team and would have avoided confusion for anyone wishing to study this gene in the future. The fact that this gene was targeted by two high-quality studies was not reflected in the genome browser, which exemplifies the difficulty of manual curation of such data. To further complicate the situation, other loci were called *MtAOC2* (MtrunA17_Chr7g0220911) by Isayenkov et al. (2005) and *MtAOC1* (MtrunA17_Chr4g0034431) by Yang et al. (2023), neither of which were targeted by loss-of-function studies so far. Developing a consistent and non-redundant set of gene names is another important and challenging task that can be addressed efficiently by a unified resource.

Another locus with a similar fate is MtrunA17_Chr8g0343011. It was called *MtLOX9* in the first loss-of-function study (Li et al., 2022), *MtLOX24* in the second, unrelated study (Xu et al., 2024), and *MtLOX4* in the genome browser v. 5.1.9. Like in the first example, the authors of the second study were unaware of the previous work on this gene, which was not referenced in the genome browser. It may be argued that the problem is minor because the groups studied different processes. However, under certain circumstances, funding, time, and material resources could be wasted by duplicating earlier efforts. This possibility should serve as a motivation for the research community to establish a system that integrates all loss-of-function data into corresponding genome browsers, not only for *M. truncatula*.

### One name for multiple genes: ambiguity

A case of the opposite type can be exemplified by studies that gave the same name to different and sometimes unrelated loci. *MtCHS* was a name given to three different genes: MtrunA17_Chr7g0252331 (Zhang et al., 2009), MtrunA17_Chr7g0220211 (Samac et al., 2011), and MtrunA17_Chr1g0201391 (Cook et al., 2025). None of these loci were named or referenced in the genome browser v. 5.1.9. Other examples are the loci called *MtMYB1* and *MtbZIP60* in original articles. *MtMYB1* is MtrunA17_Chr7g0242521 in Floss et al. (2017) but MtrunA17_Chr3g0080861 in Li et al. (2022). *MtbZIP60* is MtrunA17_Chr1g0171291 in Ribeiro et al. (2022) but MtrunA17_Chr4g0035414 in Xing et al. (2024). Only one of them was correctly referenced in the genome browser v. 5.1.9 (*MtMYB1*, Floss et al., 2017). To avoid confusion, we added lowercase letters to the names of these seven genes, in the chronological order (*MtCHSa*, *MtCHSb*, *MtCHSc*; *MtMYB1a*, *MtMYB1b*; *MtbZIP60a*, *MtbZIP60b*, respectively). *MtSymSCL1* was a name given to locus MtrunA17_Chr1g0183641 by Kim & Nam (2013) and locus MtrunA17_Chr4g0057891 by Rey et al. (2017). In the current dataset, we use alternative names of these genes: *MtSCL2* and *MtRAD1*, respectively, which are correctly named and referenced in the genome browser v. 5.1.9.

### Converging acronyms: even more ambiguity

The loci *MtCRA1* and *MtCRA2* fall into a special sub-category. In contrast to the genes mentioned above, these names were given to unrelated loci. A locus with still unknown identity (Data S6) was called *MtCRA1* for COMPACT ROOT ARCHITECTURE 1 (Scholte et al. 2002; Laffont et al., 2010). Locus MtrunA17_Chr3g0140861 was named *MtCRA2* (the same acronym) in Huault et al. (2014) and several subsequent studies. However, two other genes were named *MtCRA1* and *MtCRA2* for CHITINASE-RELATED AGGLUTININ 1 and 2, respectively (Data S6), although they have not been targeted by loss-of-function studies so far (Tian et al., 2013; Zhang et al., 2016a). Thus, their names are not revised and were not included in the current dataset.

### Incorrect locus naming in the genome browser v. 5.1.9

We found numerous examples of gene names that conflict with naming from original loss-of-function studies. We summarized these examples in Tables 3 and S1. At least 11 names were incorrectly assigned to other loci in the genome browser v. 5.1.9. These names were already used for genes characterized by loss-of-function approaches. Two names, *MtChOMT1* and *MtPRP1*, were misassigned to two genes each. For example, MtrunA17_Chr8g0366751 is the locus ID of a gene called *MtMYC2* by Guo et al. (2024). However, in the genome browser, a different member of this family was named *MtMYC2*: MtrunA17_Chr5g0411341. This gene was called *MtBHLH2* by Deng et al. (2019) (see Data S1, Data S3, and Table S1). These examples further emphasize the need for a unified resource that would implement and regulate gene naming based on original publications or some other systematic principle.

### Splitting one characterized gene model into two

Genome annotation is a dynamic process. Naturally, some older gene models characterized some time ago do not correspond to a single locus in the current genome annotation (v. 5.1.9). Such models are now split into two loci each. This is relevant to the annotation of these loci because it introduces functional ambiguity. Deciding which of two resulting loci was affected in an original study is not a trivial task. In some cases, both loci might have been affected. In a unified system proposed here, such a decision may require an independent expert who re-examines the original data. Alternatively, it can be based on the manual input of a team that produced the data. In this article, we simply draw attention to such models, with a few examples.

Functional analysis of v4 locus Medtr7g113890 (*MtGSHSa*) was conducted more than 20 years ago (Frendo et al., 2001, 2005). Now this gene model is split into two loci: MtrunA17_Chr7g0273151 and MtrunA17_Chr7g0273161. Which of these two transcripts was affected by two antisense constructs in the original study? Probably both, even though they share no sequence similarity. Although the article provides no sequence data of their constructs, the mRNA of locus MtrunA17_Chr7g0273151 was likely to be co-affected with a related locus MtrunA17_Chr7g0273131 by the shorter of the two antisense constructs (1.1 kb) because this pair has 76% identity over the length of 859 nt. The mRNA of the second locus, MtrunA17_Chr7g0273161, was probably co-affected with a related locus MtrunA17_Chr7g0273141 by the longer of antisense constructs (1.8 kb) because this pair has 75% identity over the length of 815 nt. Thus, the reported phenotype can be attributed to all four loci unless a dedicated study shows otherwise. Unfortunately, this information was not reflected in the genome browser v. 5.1.9, nor was the fact that the loci were targeted by a loss-of-function approach. In this and a few other cases, we attempted to fill this gap by assigning unique but related names to these loci (*MtGSHSa* and *MthGSHSa*; *MtGSHSb and MthGSHSb*, respectively). Specifically for these homologs, we modified the original names introduced by Frendo et al. (2001, 2005), where “h” stands for “homo”. They are fully explained in Supporting Results. This group of genes presents special interest to our study and will be discussed in detail in the context of ncORFs (Data S12 to S14 and Data S16).

The second v4 locus of this type is Medtr4g088055 (*MtROP7*, Wang et al., 2021), which is now split into MtrunA17_Chr4g0046211 and MtrunA17_Chr4g0046221. These two loci are separated by only one nucleotide (A) in the plus-strand and share no sequence identity. We propose names *MtROP7a* and *MtROP7b*, respectively, for these loci. RNAi constructs used in the original study (Wang et al., 2021) target the first locus (*MtROP7a*) together with many other genes, including *MtROP8* and *MtROP10*. The alignment of these RNAi constructs with the second locus (*MtROP7b*) yields no obvious similarity, except for a 40-nt-long region having 88% identity with the RNAi construct produced by primers Forward (MROP10) 1 and Reverse (MROP10) 1 (Supplemental Table 2 in Wang et al., 2021). Is this very local similarity sufficient to affect *MtROP7b*? Probably not, because much longer sequences (>130 nt) are typically used for efficient RNAi-mediated silencing in plants (Kakiyama et al., 2019). Only the *MtROP7b*-specific qRT-PCR analysis can tell. With so little evidence linking this gene to the conditional phenotype reported by Wang et al. (2021), *MtROP7b* would be removed from our dataset as being unaffected by the loss-of-function experiment. However, the following consideration shows how important it is to look deeper into such examples and to inspect the original qRT-PCR data on a case-by-case basis. Analysis of *MtROP7*-designed qRT-PCR primers used by Wang et al. (2021) clearly shows that the new model is wrong and the old one (v4) is correct (Supplemental Table 2 in Wang et al., 2021). The forward primer (TTATGCTCCAAATGTGCCTA) is located in *MtROP7a* whereas the reverse primer (TGTTTTAGAGCTGCACTCA) resides in *MtROP7b*. They can form an amplicon of 269 bp exclusively from the cDNA of the old model. If the signal reported in Figure S5 of Wang et al. (2021) had come from contamination with genomic DNA, the amplicon would have been 401 bp long because of Intron 4. The melting curve analysis would certainly have alerted the researchers about such contamination. Thus, the reported data is very likely to be correct and unambiguous. This is not the only example of a “longer” original model making more sense because of being supported either by transcriptomic or by MS proteomic data (see the example of *MthGSHSb* in Data S16 Slide 7). Nevertheless, it is important to look deeper into the reason for rejecting the old model by the current genome annotation. The 58-bp-long junction between the ORFs of *MtROP7a* and *MtROP7b* is free of stop codons only in frame 3 relative to the sequences, which is not the reading frame of the annotated ORFs. Thus, the sequences are separated by in-frame stop codons. Possibly, the junction is a spliced-out intron, which could explain why in-frame stop codons in this region were tolerated by natural selection. It is worth noting that neither of these two loci were provided with the correct reference in the genome browser v. 5.1.9 (Data S6).

Another example of this sort is v4 locus Medtr4g073400 (*MtSYT1*, Gavrin et al., 2017), which is now split into MtrunA17_Chr4g0037111 and MtrunA17_Chr4g0037121 (*MtSYT1a* and *MtSYT1b*, respectively, in our dataset). As in the case of *MtROP7*, the loci are separated by a single nucleotide (A) in the plus-strand. Likewise, qRT-PCR primers MtSyt1-F AGGAGGCCCGTTGGAATTTT and MtSyt1-R GTGATGCTTCCCCTCAACA (Supplementary Table 1 in Gavrin et al., 2017) span the junction between the two loci, forming an amplicon of 625 bp from both cDNA and genomic DNA, according to the new model. The length of 625 bp is clearly suboptimal for qRT-PCR (normal range is between 60 and 150 bp, Udvardi et al., 2008). Evidently, the qRT-PCR signal shown in Supplementary Figure 7 of Gavrin et al. (2017) could come from the cDNA corresponding to the “old” model, if the junction that contains in-frame stop codons was spliced out. We doubt that the qRT-PCR settings used in that high-quality study would permit the generation of a signal exclusively from the contamination with genomic DNA, which would result in this unusually long amplicon (625 bp). For this reason, in our dataset, we consider *MtSYT1a* and *MtSYT1b* as part of the same functional unit, as it was described in the original study. If the new model is correct, it is impossible to find out if both loci were affected by the generic RNAi construct designed for simultaneous silencing of *MtSYT1* and *MtSYT3*, which share no sequence identity. This is because RNAi primers MtSyt1-F TGGATCCATCACAGGCCATG and MtSyt1-R CGCAAACGGATTTGTGTGGT (Supplementary Table 1 in Gavrin et al., 2017) do not perfectly match the cDNA of the split models MtrunA17_Chr4g0037111 and MtrunA17_Chr4g0037121. Thus, functional ambiguity does not permit exclusion of either locus from the reported RNAi phenotype even though only MtrunA17_Chr4g0037111 was provided with the correct loss-of-function reference in the genome browser v. 5.1.9 (Data S6).

The last example of a split locus is v4 model Medtr1g102190 (*MtLICK1*, Wang et al., 2025). It corresponds to two adjacent loci in the genome assembly v. 5.1.9: MtrunA17_Chr1g0204221 and MtrunA17_Chr1g0204231 (*MtLICK1a* and *MtLICK1b*, respectively, in our dataset). They are separated by a nucleotide triplet ACA in the minus-strand. If this newer model is correct, only MtrunA17_Chr1g0204221 was affected by the *Tnt1* insertion in Wang et al., 2025. Based on FST NF9810_low_9 (the *Tnt1* mutant collection, Tadege et al., 2008; Lee et al., 2018; Kaur et al., 2021), the insertion is located in the 5’-UTR of this gene. However, locus MtrunA17_Chr1g0204221 is a direct continuation of locus MtrunA17_Chr1g0204231. The high similarity of exon-intron structures shown in Extended Data Fig. 2k and i (Wang et al., 2025) suggests that the new model may be incorrect. Thus, the insertion is predicted to be in the last exon of the v4 model. If it were the 5’-UTR, the mutant phenotype would probably not differ from the wild type. The study provided further proof for the validity of the original model by MS proteomics. Specifically, Table 1 in Wang et al. (2025), which illustrates MtLICK1 phosphorylation sites, lists a peptide MSI(pT)VGAAEGLAYLHHEANPHIIHR with the original position of the phosphorylation site at T151. It maps to the 3’-UTR of MtrunA17_Chr1g0204231 in such a way that 306 nt located between MSITVGAAEGLA and YLHHEANPHIIHR in the current model are missing. The same concerns peptides RM(pS)ITVGAAEGLAYLHHEANPHIIHR (S149) and RMSITVGAAEGLA(pY)LHHEANPHIIHR (Y160). This indicates that this part of the junction between MtrunA17_Chr1g0204231 and MtrunA17_Chr1g0204221 is a spliced-out intron, like in the cases of *MtROP7* and possibly *MtSYT1*. In contrast, peptides DIKA(pS)NVLLDTEFQAK (S177) and ASNVLLD(pT)EFQAK (T183) map exactly to the junction site between MtrunA17_Chr1g0204231 and MtrunA17_Chr1g0204221, which means this region is a translated exon. Thus, we consider MtrunA17_Chr1g0204231 and MtrunA17_Chr1g0204221 as a single functional unit in our dataset, as it was described in the original study. Unfortunately, neither of these loci is provided with the correct reference in the genome browser v. 5.1.9 (Data S6).

### Duplicated gene models

A different type of challenge in assigning a mutant phenotype to an annotated locus is exemplified by a v4 gene Medtr4g125520. It corresponds to two entries in the genome assembly v. 5.1.9: MtrunA17_Chr4g0070611 and MtrunA17_Chr4g0070641. In contrast to the splitting of an old model into two, as described in the previous section, these two loci are separated by several other genes and repeat elements. Although the gDNA sequences of MtrunA17_Chr4g0070611 and MtrunA17_Chr4g0070641 are identical base-for-base over the shared region, the loci are located in separate genomic positions. The shorter locus (MtrunA17_Chr4g0070641) entirely lacks the region corresponding to the end of Exon 3 of MtrunA17_Chr4g0070611. The gene model presented in Figure 5a of Liu et al. (2014) corresponds to MtrunA17_Chr4g0070611. Thus, only this longer locus is listed in the current dataset. However, if both models are correct, it is impossible to tell which one was affected by the *Tnt1* insertions because the gDNA sequences around the insertion sites are identical in these two genes.

We attempted to solve this ambiguity by analyzing FST sequences of *Tnt1* insertions corresponding to lines *myb14-1* and *myb14-2* (Figure 5a of Liu et al., 2014). These are FSTs NF14565_low_42 and NF0084-INSERTION-2, respectively, both in reverse orientation. The first FST is not informative for our purpose because its end reaches only 14 bp upstream of the translational start (ATG) codon of MtrunA17_Chr4g0070611 and MtrunA17_Chr4g0070641. These 14 bp do not match the gDNA of either locus in the Jemalong A17-based genome browser v. 5.1.9. They are likely to present a sequencing error because they also do not match a corresponding model in the ecotype R108 (the background of the *Tnt1* mutant population; Tadege et al., 2008; Lee et al., 2018; Kaur et al., 2021). The second FST, NF0084-INSERTION-2, gives more information because its end reaches 612 bp upstream of the ATG. While corresponding regions in the Jemalong A17 genome are identical between MtrunA17_Chr4g0070611 and MtrunA17_Chr4g0070641, there is a single matching sequence in the R108 genome. It is immediately upstream of a gene nearly identical to MtrunA17_Chr4g0070611 at the gDNA level (only three mismatches over the entire length of MtrunA17_Chr4g0070611). Thus, we conclude that the second (short) copy may exist in the ecotype Jemalong A17 but is absent from the ecotype R108, which suggests a very recent duplication. This is further supported by the qRT-PCR-mediated test for gene copy number presented in Supplemental Figure S11 of Liu et al. (2014). Primers MtMYB14qPCR1-F AGGCAACATATCCTCAGATGAAG and MtMYB14qPCR1-RTCGTGTTCCAATAATTCTTGATTTCA (Supplemental Table 4, Liu et al., 2014) reside in a region shared between MtrunA17_Chr4g0070611 and MtrunA17_Chr4g0070641. The signal generated by these primers corresponds to a single genomic copy. This is an example leaving a big question about the functional status of both loci in Jemalong A17: which of them is a functional ortholog of the single copy in R108? Even with a targeted mutagenesis tool such as CRISPR/Cas9 it may be challenging to discriminate between these loci because of the high nucleotide identity within, upstream, and downstream of the sequences. Future functional annotation of such loci should probably indicate ambiguities related to the differences between the genomes of ecotypes A17 and R108.

### Loss-of-function methodology and biological processes studied in *M. truncatula*

Diverse methods were used for loss-of-function analyses of 673 *M. truncatula* genes (22 approaches, Figure S1). More than half of the studies applied one (or a combination) of just two methods: insertional mutagenesis of tobacco retrotransposon *Tnt1* (359 genes) (Tadege et al., 2008; Lee et al., 2018; Kaur et al., 2021) and post-transcriptional silencing mediated by RNAi (265 genes) (Sharp & Zamore, 2000; Baulcombe, 2004; Chaudhary et al., 2024). As a new method, gene editing with the aid of CRISPR/Cas9 (Doudna & Charpentier, 2014; Guo et al., 2023) was used for only 87 genes so far. The popularity of RNAi as a tool is remarkable given the known limitations and ambiguity associated with this method. Technically, it is a simple and inexpensive approach when the generation of transgenic roots (so-called hairy roots, Nilsson & Olsson, 1997; Limpens et al., 2004), rather than the entire transgenic plants, is sufficient for the analysis. Out of 117 genes first characterized in 2023-2025, 30 were studied by RNAi, which is almost the same as the number of genes studied by CRISPR/Cas9 in the same period (31 genes). *Tnt1* was used for more than half of the 117 genes (67 genes), followed by FNB (11 genes), EMS (four genes), and AS (one gene) in the same period. FNB, EMS, and AS stand for Fast Neutron Bombardment mutagenesis (Li et al., 2001), Ethyl Methanesulfonate mutagenesis (Sega, 1984), and Antisense-mediated gene silencing (Waterhouse et al.,1998), respectively. From this data, it is evident that the *Tnt1* resource plays a pivotal role in the functional genomics of *M. truncatula* (Data S6, Figure S1).

*M. truncatula* was used for studies on 97 biological processes (Data S9, Figure S2). However, this legume was primarily established as a model for SNF. More than half of 673 genes were tested for SNF (380 genes), followed by root development (166 genes), and AM (141 genes). Among non-symbiotic processes other than root development, *M. truncatula* was frequently tested for leaf development (104 genes), flower development (60 genes), control of body size (48 genes), floral transition (48 genes), and pod development (40 genes). Seed-related studies are also numerous in *M. truncatula* (93 genes): seed development (47 genes), seed composition (27 genes), seed germination (8 genes), seed longevity (5 genes), seed coat function (3 genes), and seed dormancy (3 genes).

### More than half of the genes studied in *M. truncatula* are transcription factors and enzymes

Transcription factors (TFs) and enzymes are two protein groups prevailing in the current dataset (224 and 205 genes, respectively). Counts of these and other protein groups are shown in Figure 3, Figure S3, Data S10, and Data S11. Among TFs, the top four most represented families are AP2-EREBP (21 genes), GRAS (19 genes), MYB-HB-like (18 genes), and SBP (17 genes). Among enzymes, the most represented classes are kinases (48 genes), oxidoreductases (45 genes), transferases (43 genes), and hydrolases (39 genes). Given the pivotal role of transport proteins in SNF, AM, and other biological processes, transporters (73 genes) and channels (13 genes) are also well-represented in the current dataset. Remarkably, transport substrates were found for many of these proteins (60 transporters and 7 channels), which is usually a challenging task.

**Figure 3.**
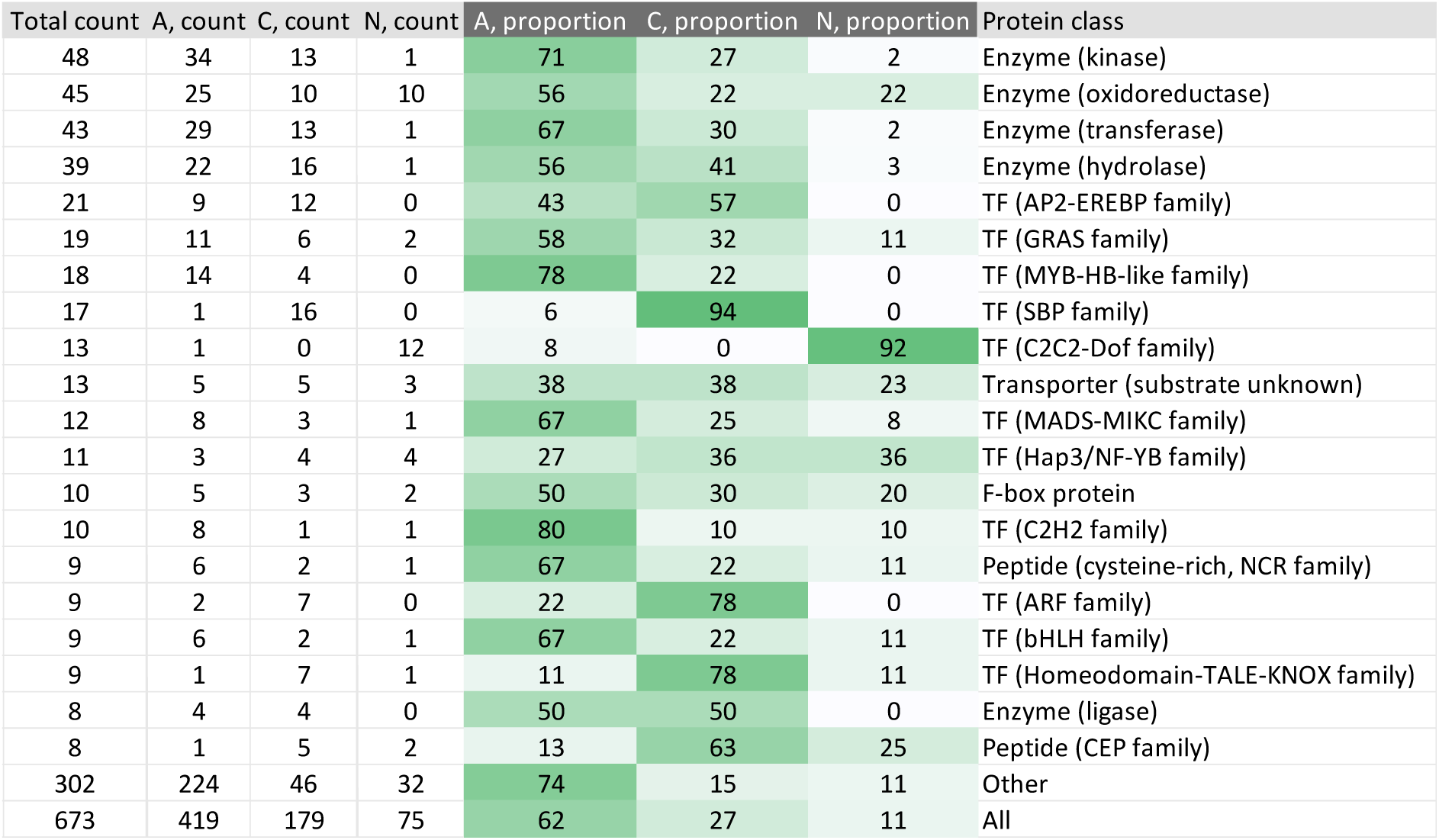
Different protein classes have different proportions of phenotypes, which may reflect their relative functional importance. Letters A, C, and N stand for altered, conditional, and neutral phenotypes, respectively. The graph shows only the top-20 protein classes. The complete list of protein classes and corresponding statistical analysis are available in Data S10. The summary on phenotypes of combined protein groups is available in Data S11.

### Non-conditional loss-of-function phenotypes are found in 62% of genes studied in *M. truncatula*

We have categorized phenotypes of 673 genes studied by loss-of-function approaches into three categories. This classification was based exclusively on the genetic conditions. Specifically, it did not consider conditions such as different growth media or temperature regimes. Phenotypes caused by single genes were categorized as non-conditional (A for altered). Phenotypes caused by more than one gene were regarded as conditional (C for conditional). Neutral phenotypes (N for neutral) were those that did not differ from the wild type, regardless of the number of genes affected in each line. The overall proportions of A-, C-, and N-phenotypes were 62%, 27%, and 11%, respectively (Figure S4). However, these proportions differed slightly between SNF phenotypes and non-SNF phenotypes (compare Figures S4a and S4b). The proportion of C-phenotypes was higher among genes tested for SNF (31%), compared to non-SNF phenotypes (26%). At the same time, the proportions of N-phenotypes did not differ much (16% and 15% for SNF and non-SNF, respectively). Only 53% of genes tested for SNF gave non-conditional phenotypes (A-phenotypes), which suggests functional redundancy due to the robust structure of the genetic program controlling SNF.

This relatively low “success rate” of loss-of-function studies on SNF reflects the challenge in identifying crucial genes that can be used for the improvement of SNF in legume crops and synthetic biology efforts to make non-legume plants undergo SNF. With some effort, Data S6 can help predict the likely “success rate” of planned loss-of-function studies for specific biological processes, as we showed for SNF. For example, all 11 genes tested for cold stress showed an A-phenotype for this biological process (higher sensitivity or lower sensitivity). Hence, other genes possibly involved in cold stress are likely to have scorable A-phenotypes, too. In contrast, out of 60 genes tested for flower development, not more than half exhibited A-phenotypes. Because Data S6 organizes phenotypes at the gene level, additional processing is required to examine them in the context of individual biological processes tested in each study. In the next section, we focus on statistically significant differences in this “success rate” among different protein classes and groups of classes. Extracting this information from Data S6 is simple because phenotypic categories are linked to genes. Moreover, we find this type of data more relevant for academic research and breeding efforts because it can tell which gene groups are more likely to yield a scorable phenotype.

In the remaining part of this section, we present data on the agreement between phenotypic categories when the same genes are tested for SNF and non-SNF phenotypes (Figure S5). Among 219 genes tested for SNF and at least one non-SNF phenotype (Data S5), 40% had A-phenotypes for both processes (group AA). For conditional (CC) and neutral (NN) phenotypes, this proportion was 25% and 5%, respectively (Figure S5). Cases of discordance regarding phenotypic categories were not very prominent among genes tested for both types of biological processes. The largest discordance group (10%) was represented by genes that exhibited A-phenotypes when tested for non-SNF processes and N-phenotypes when tested for SNF (group AN in Figure S5). The reciprocal group, NA, accounts for only 5% of 219 genes, which emphasizes that purely SNF-specific genes are not very abundant. These observations suggest the relatively large number of genes with pleiotropic effects, as can be seen also for various non-SNF phenotypes in Data S6. In summary, the chance that a gene selected for analysis will give an A-phenotype for at least one biological process is 62% (Figure S4c). If a gene shows an A-phenotype for SNF, in 40% of cases, it will also have a scorable A-phenotype for at least one non-SNF process (Figure S5b). This may reflect the adoption of non-symbiotic genes for symbiotic processes reported in many studies (Limpens et al., 2003; Soyano et al., 2019; Huisman & Geurts, 2019; Liu & Bisseling, 2020; Hawkins & Oresnik, 2022).

### Unique proportions of phenotypic categories in different protein classes may reflect their relative functional importance

As shown in Figure 3, the proportions of phenotypic categories are not the same for different protein classes. Likewise, different groups of combined protein classes have different proportions of phenotypic categories (Figure S3). The natural question is whether these differences are statistically significant. Data S10 and S11 present statistical analyses of these differences for protein classes and larger groups, respectively. Sheet “Sorted by count” in Data S10 shows which classes have A:C:N ratios significantly different from the overall ratio of 62:27:11 (see also Figure S4c). Four classes differ significantly from that ratio: TFs of the SBP family and the C2C2-Dof family, channels with unknown substrates, and protease inhibitors. TFs of the SBP family have the ratio of 6:94:0, which corresponds to 16 out of 17 tested SBP genes having C-phenotypes. This, however, is a special case because nearly all these SBP genes were affected by miRNA in a single study (Wang et al., 2019). A somewhat similar situation is observed in TFs of the C2C2-Dof family (the ratio of 8:0:92), where 12 out of 13 genes were targeted in a single study (Zhang et al., 2019). In contrast to the SBP family, these genes were targeted in separate *Tnt1* lines and exhibited N-phenotypes. Channels with unknown substrates fall into the same category (the ratio of 17:0:83), with four out of seven genes targeted in a single study (Charpentier et al., 2016). Protease inhibitors have the ratio of 20:0:80, with four out of five genes exhibiting N-phenotypes. Much more interesting differences are observed when individual classes are cross-compared with each other rather than with the overall ratio of 62:27:11. Sheet “Example comparisons” in Data S10 highlights highly significant differences between related classes such as kinases and oxidoreductases, transferases and hydrolases, TFs of the AP2-EREBP family and the MYB-HB-like family, etc. For example, kinases have a much higher proportion of A-phenotypes compared to oxidoreductases and a much lower proportion of N-phenotypes, which reflects their crucial roles in various processes. Likewise, TFs of the MYB-HB-like family have a much higher proportion of A-phenotypes compared to other TF groups, such as the AP2-EREBP family and the GRAS family. TFs of the AP2-EREBP family have an unusually high proportion of C-phenotypes (57%) but no N-phenotypes among 21 tested genes. This information can be of use for practical purposes because it provides a quantitative measure of relative functional importance and redundancy.

When combined functional groups are considered (Data S11), similar significant differences are evident. Channels stand out as a group with the A:C:N ratio of 62:0:38, which is significantly different from the overall ratio of 62:27:11. They have an abnormally low proportion of C-phenotypes and an abnormally large proportion of N-phenotypes. Likewise, the combined group of protein classes designated as “Other” in Data S11 (see Data S10 for the classes included in this group) has the ratio of 73:12:15, which is significantly different from the overall ratio of 62:27:11. It has an abnormally high proportion of A-phenotypes. Sheet “Example comparisons” in Data S11 exemplifies significant differences between combined groups. For example, the ratio of 71:18:11 for transporters is significantly different from the ratio of 53:34:13 for TFs, which literally translates into a higher chance of observing a scorable A-phenotype when genes under study are transporters.

It should be noted that 73% of A-phenotypes (group Other) is the largest proportion only within Dataset S11 (differences between combined groups). Much larger proportions can be found in individual classes with at least 5 genes per class (Data S10). The following classes have proportions of A-phenotypes larger than 73%: Cytochrome P450 (100%, n=6), TF (bZIP family) (100%, n=5); TF (NAM family) (83%, n=6), TF (C2H2 family) (80%, n=10); TF (Homeobox-WOX family) (80%, n=5); and TF (MYB-HB-like family) (78%, n=18). TFs exhibit a very large variation in proportions of A-phenotypes. For example, the following TF families have rather low proportions: TF (SBP family) (6%, n=17); TF (C2C2-Dof family) (8%, n=13); TF (Homeodomain-TALE-KNOX family) (11%, n=9); TF (ARF family) (22%, n=9); TF (Hap3/NF-YB family) (27%, n=11); TF (HD-SAD family) (40%, n=5); and TF (AP2-EREBP family) (43%, n=21).

Lastly, it is worth noting protein classes with the largest proportions of N-phenotypes (Data S10): TF (C2C2-Dof family) (92%, n=13); Channel (substrate unknown) (83%, n=6); Protease inhibitor (80%, n=5); Transporter (proton efflux) (60%, n=5); Membrane protein (PLAT domain containing) (40%, n=5); TF (Hap3/NF-YB family) (36%, n=11); Peptide (CEP family) (25%, n=8); Transporter (substrate unknown) (23%, n=13); and Enzyme (oxidoreductase) (22%, n=45). Channels, regardless of the substrate, are also prominent in Dataset 11, with the largest proportion of N-phenotypes among combined groups (38%, n=13).

In practical terms, the meta-analysis presented in this section may encourage the research community to focus on classes with the higher “success rate”. However, it should not bias researchers and funding agencies against candidate genes with relatively low “success rates” as they can still be highly informative and useful in breeding programs and synthetic biology efforts (e.g. *MtDASH*, *MtDMI1*, *MtHA1*, and many other genes in Data S6).

### NcORFs in *M. truncatula* genes targeted by loss-of-function approaches

Due to space limitations, the entire Results section dedicated to ncORFs was placed into Supporting Results, which are available in the online version of this article as a separate file. This section is crucial to the message of this paper, which can be summarized as follows: ncORFs cannot be ignored in functional studies as they introduce ambiguity regarding the interpretation of mutant phenotypes. The section references Data S12-S18 and has its own list of literature sources.

## DISCUSSION

### The genome annotation of *M. truncatula* requires better integration of loss-of-function data

The latest release of the *M. truncatula* genome (v. 5.1.9) is an indispensable reference for plant biologists, especially for studies that take advantage of the symbiotic properties of this model organism (Pecrix et al., 2018). However, the resource lacked proper integration of information on loss-of-function phenotypes. This concerns not only the detailed description of phenotypes but also relevant references to loss-of-function studies (Figure 2). Consequently, a researcher who considers targeting a locus of interest may be unaware of any previous work on that locus. This limitation has multiple negative effects on the progress of functional genomics in *M. truncatula*, including duplication of efforts, assigning multiple names to the same genes, and being unable to benefit from the evidence generated in previous experiments. It could be argued that obtaining such information is the responsibility of researchers. However, a well-annotated and user-friendly resource should aim to facilitate information retrieval, not merely leave it to the users. The tool should minimize not only the effort but also the ambiguity as to which locus was affected, with little space left for misinterpretation. During our meta-analysis, we have noticed that establishing a correct link between a locus described in a loss-of-function study and the latest identifier of that locus is not always a trivial task. Often, it requires careful sequence analysis of primers, RNAi segments, and even the conversion of DNA/protein sequences from image to text. This concerns not only old studies conducted before the advent of the genome browser but also new studies. On 10 April 2026, our dataset was integrated into the genome browser of *M. truncatula* for the maximal benefit of the research community. It will be maintained and updated by the corresponding author (see Data Availability).

### Keeping the loss-of-function domain of the browser up to date may require a human-curated protocol

Once the data becomes part of the genome browser, updating it in real time, or at least frequently, presents a considerable challenge. We have noticed that many details of the phenotypes and other relevant information summarized in Data S6 may be extremely difficult to extract automatically. A human-curated protocol allowing direct submission may be the only adequate way to record and consolidate error-free data on loss-of-function phenotypes. This, however, is not a task that can be handled in a background mode by a dedicated staff member of the genome browser. Integrating numerous functional genetic studies (Figure 1) can be highly time-consuming, particularly given the steadily increasing number of publications each year. Therefore, we propose that such information be requested directly from the authors of loss-of-function studies. A convention can be established as to the rules and the format of entering the information on newly characterized genes directly into the browser shortly before a publication is transferred to the print stage by a journal. Each submission can have a unique identifier that researchers can cite in their publications. If successful, this concept can become a standard not only for *M. truncatula* but also for other species. Generalizing such a submission system, however, would be the next challenge, this time of an organizational type. It would require coordination between multiple institutions, resources, and publishers in such a way that the journals make it compulsory for authors to register their mutant data. Importantly, this should concern not only phenotypes of genes that were clearly in the focus of a study (for example, their names were mentioned in a title or in an abstract) but also phenotypes of mutants that had already been published in the same or a different context earlier. Typically, such mutants have an auxiliary role in a study. However, they often provide additional details unnoticed or not addressed in the original articles. Many phenotypes in our dataset fall into this category. Data S1-S6 also list genes that were mutagenized only once; however, they did not constitute the core of articles describing their phenotypes. Thus, such genes can be overlooked in the literature because the fact of their mutagenesis was not highlighted anywhere except for a few sentences and/or supplementary figures. Extracting such information and assigning it accurately to specific loci requires a human effort that cannot be easily automated.

The network linking the genome browsers and the journals should be the future of functional genomics. It would eliminate many limitations mentioned in the previous section, thus facilitating the progress of biological research. At the same time, one format described in the next section can serve as a model for other resources (Baldarelli et al., 2024). It was successfully implemented with an option of direct submission for mouse mutants.

The only significant problem with the direct submission system proposed here is cybersecurity. A very considerable effort is required to ensure that direct submissions do not open the door to malware intended to damage the resource. This is the only reason why our recently opened loss-of-function track in the *M. truncatula* genome browser will continue to function through one curator. In the Data Availability section, we encourage the *M. truncatula* community to send their pre-formatted data to the corresponding author. IK will mediate the integration of the information into the browser.

### Strong points and limitations of existing loss-of-function databases

Historically, the first attempt to create a multi-species genotype/phenotype database was made at the dawn of the genomic era, before the *M. truncatula* genome became well-annotated. This database, called PhenomicDB, included mouse, zebrafish, fruit fly, *Caenorhabditis elegans*, yeast, and *A. thaliana* (Kahraman et al., 2005; Groth et al., 2007). It contained 399,772 phenotypes associated with 77,400 genes. Unfortunately, the URL of PhenomicDB, http://www.phenomicdb.de, has been offline since 2010. It is not available in an alternative form.

Currently, the most advanced loss-of-function resource for animal organisms is Mouse Genome Informatics (MGI) (Baldarelli et al., 2024). It is closely associated with MUTAGENETIX^TM^, a database of N-ethyl-N-nitrosourea (ENU)-induced mutations that cause phenotypes in mice (Wang et al., 2015). MGI is a well-structured, comprehensive database containing information on 20,583 genes with phenotype annotations (as of 25 February 2026). It enables filtering phenotypes by the mutagenesis method (the methods are not limited to the treatment with ENU), allele attribute (conditional, dominant negative, no functional change, etc.), type of mutation (deletion, insertion, duplication, etc.), and even the anatomical system affected by the phenotype. It shows human homologs of affected genes and homologs in other mammalian organisms. Importantly, the genome browser of MGI has a comprehensive portfolio for each locus describing all possible aspects of corresponding mutants, including references to original experimental studies. The resource accepts direct submissions of phenotype data, which makes it ideal for efficient functional analysis. Its format and depth should be the model for browsers of other organisms. The only drawback of the MGI browser is the absence of a track for ncORFs (as of 25 February 2026), which does not permit convenient checking for ncORFs potentially co-affected in mutagenesis studies. The *M. truncatula* has such a track, with information on chimeric and non-chimeric ncORF MS-validated in our study (Çakır et al., 2025b).

The most comprehensive analysis of loss-of-function phenotypes in a plant organism was conducted in *A. thaliana* (Lloyd and Meinke, 2012). There is little to add to the deep coverage of that study, which is much more versatile compared to the basic analysis we present in this manuscript. The current genome browser of *A. thaliana* (Araport11, Cheng et al., 2017) contains a field that summarizes loss-of-function and other important data for each affected locus (Curator_summary). Unfortunately, that field contains no references to original studies that established the link between a locus and its phenotype. Likewise, no details are provided as to the genes potentially contributing to the phenotype. Thus, a user does not learn if the phenotype is neutral, genetically conditional (more than one gene involved), or caused by a single gene. Regarding the references, the *M. truncatula* genome browser has been one step ahead of the *A. thaliana* browser as it has been providing relevant references to at least some genes targeted by loss-of-function approaches (Figure 2). Like the genome browser of *M. truncatula*, the *A. thaliana* browser has a separate track for ncORFs of the following types: uORFs, dORFs, and lncRNA-ORFs, according to the system of Mudge et al. (2022). In contrast to the *M. truncatula* browser, the ncORF track does not include intORFs, uoORFs, and doORFs (as of 25 February 2026), presumably because they are difficult to discriminate from translated refORFs by ribosome profiling (Ribo-Seq). The ncORF data in the *A. thaliana* browser were generated by Wu et al. (2024a).

Gephebase is an excellent resource for evolutionary studies (Courtier-Orgogozo et al., 2020). It focuses on genotype-phenotype relationships in natural variation observed in animals, yeast, and plants. Unfortunately, the database excludes lab mutants. Other databases are dedicated to mutations associated with human diseases or are much smaller in their scope compared with the resources mentioned above.

Several platforms were developed for *in silico* mutant screening in plants. They include, but are not limited to, the database of *M. truncatula* mutants (Tadege et al., 2008; Lee et al., 2018; Kaur et al., 2021), TOMATOMA for tomato (Saito et al., 2011; Shikata et al., 2016), and MiRiQ for rice (Kubo et al., 2024). However, these platforms cannot substitute genome browsers with integrated loss-of-function data, such as MGI and Araport11, because most of the mutants listed in those databases have not been characterized yet.

These considerations position our resource as smaller in scale than the *A. thaliana* database, but still essential for plant biologists, as it represents the most comprehensive loss-of-function dataset available after *A. thaliana*. Importantly, the integration of ncORF information with functional data gives our dataset a unique value among comparable resources. Specifically, among all standard genome browsers, only the browsers of *A. thaliana* and *M. truncatula* have the ncORF tracks (Wu et al., 2024a; Çakır et al., 2025b). In contrast to the *Arabidopsis* browser, our ncORF track in the *Medicago* browser includes MS-validated ncORFs that are embedded in refORFs or overlap with refORFs (ncORF designated as “intORFs”, “uoORFs”, and “doORFs” in the system of Mudge et al., 2022), along with all other major ncORF types. Lastly, the *Medicago* ncORF track is currently the only one (globally) that includes chimeric ncORFs (ncORFs that are translated to chimeric peptides via programmed ribosomal frameshifting, PRF).

### About the content of Supporting Discussion

Due to space limitations, many important considerations about loss-of-function data, as well as about ambiguities associated with ncORFs and *Tnt1*, were placed into a separate file named Supporting Discussion. The file references Figure 4, supporting Figures S6-S9, supporting Tables S2 and S3, Data S19, and has its own list of literature sources. Figure 4 introduces seven types of ambiguity associated with translated ncORFs in loss-of-function studies, with real examples.

**Figure 4.**
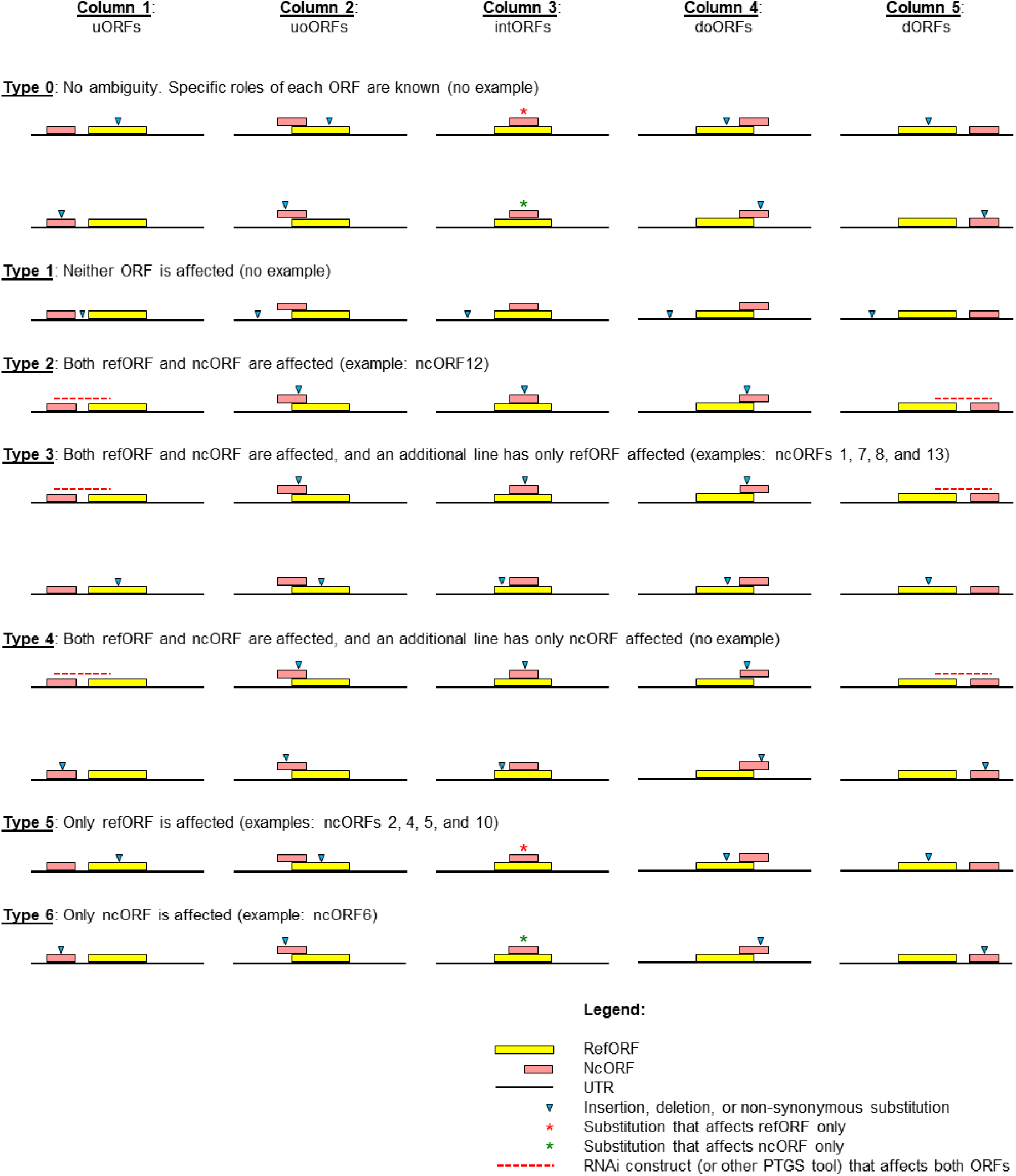
Main types of ambiguity associated with translated ncORFs in loss-of-function studies. Ambiguity types are exemplified based on the assumption that currently annotated gene models are correct. Each column corresponds to a specific location of a ncORF relative to its refORF (uORFs, uoORFs, intORFs, doORFs, and dORFs, respectively, Mudge et al., 2022). Each row corresponds to a specific scenario leading to the inactivation of one or two ORFs. The graph is a simplified illustration of the most common scenarios. It does not apply to the following situations: (1) a mutation outside both ORFs affects their transcription or translation without affecting their amino acid sequences; (2) an insertion at one location interferes with splicing at a different location, thus affecting ORFs in trans. Abbreviations: UTR, untranslated region; RNAi, RNA interference; PTGS, post-transcriptional gene silencing.

### The take-home message

Functional genomics relies upon comprehensive and accurate annotation of genes available as a genome browser. At the same time, a truly comprehensive genome browser should encompass the information on loss-of-function phenotypes. While the mouse, *A. thaliana*, and *M. truncatula* genomes are among the few to link such information to specific loci, there is currently no standard pipeline for systematic recording, storage, and displaying phenotypic data for other organisms. Here, we propose developing a centralized system that should make it obligatory for researchers to report such information at the publication stage. We also showed the limitations of the most comprehensive genome browsers, which can be summarized as the lack of information on genes co-affected in original studies (multiple knockouts or knockdowns) and missing references to such studies. Besides, the main purpose of this work was to show that two fundamental aspects of functional genome annotation have been largely overlooked in standard genome browsers. These aspects are (1) translated and conserved ncORFs and (2) trans effects of insertional mutagenesis on splicing. Although both phenomena have been well documented, they remain beyond consideration in nearly all functional analyses. We provided examples of how overlooking such information may result in major misinterpretation of loss-of-function phenotypes.

Importantly, this ambiguity is not unique to *M. truncatula*. It represents a pervasive, often ignored variable in plant functional genomics. Whether using RNAi, which inevitably targets co-transcribed ncORFs, or gene editing via CRISPR/Cas9, which can disrupt overlapping coding sequences or splicing motifs, the “one gene, one phenotype” assumption is frequently violated. Our data highlight that a significant proportion of “verified” mutants likely requires re-evaluation to rule out ncORF contributions. Thus, the framework presented here serves not merely as a curatorial resource, but as a necessary quality-control step for distinguishing causal refORF mutations from ncORF-mediated artifacts in genetic studies.

We also showed how ncORFs can help rectify doubtful gene models. Despite the relatively modest scope of the genome browser of *M. truncatula*, the integration of loss-of-function and ncORF data in one resource makes it unique and more informative than other standard browsers. In addition to summarizing the information on phenotypes of 673 genes in the present manuscript, we mapped 805 non-chimeric and 156 chimeric MS-validated ncORFs to corresponding loci in the genome browser. To the best of our knowledge, after the ncORF study conducted in *Arabidopsis* (Wu et al., 2024a), this is the second study to merge loss-of-function data with ncORFs in any organism. At the same time, it is the first attempt to reconsider published loss-of-function phenotypes in the light of knowledge about translated ncORFs expressed from the same loci. We expect that this resource will serve as the model for browsers of other organisms, which will ultimately improve the quality of functional analyses for the maximal benefit of biomedical research, sustainable agriculture, and biotechnology.

Our meta-analysis of loss-of-function studies produced unique data on family-wise proportions of altered, conditional, and neutral phenotypes in different protein classes and families. This type of resolution is so far unprecedented because other studies mostly reported frequencies of essential genes only for heterogeneous classes that combine members of diverse families. Our list of tripartite phenotype proportions can be used by legume biologists and breeders as a cheat sheet to focus on candidate genes that are likely to give a scorable phenotype. We explicitly state that such a resource may have validity only within the legume family or even *M. truncatula* alone. Like any cheat sheet, it should be used constructively: gene families with low proportions of altered phenotypes should not be deprived of due attention.

The manuscript also coins several ideas in the field of ncORFomics. To the best of our knowledge, these ideas are novel. First, based on earlier reviews and our new data, we developed, discussed, and exemplified the classification of ambiguities associated with ncORFs in loss-of-function studies. Second, we explained why the analysis of taxonomic ranges of ncProts’ conservation should become a standard component of genome annotations. Third, we advocated the potential biological relevance of ncProts whose conservation signatures are limited to the species of origin. Fourth, we showed how certain translated ncORFs can systematically remain undetected due to regions of 100% similarity to refProts in the species of origin. Lastly, we provided examples in which very strong conservation of ncORFs at the amino acid level may not be a reliable marker of their translation. These considerations will help better understand the biological roles of ncORFs, separately from refORFs, as an integral component of the organismal translatome.

## MATERIALS AND METHODS

### Meta-analysis of loss-of-function studies

Literature on loss-of-function studies in *M. truncatula* was collected manually using Google Scholar, PubMed, and other search engines. Main texts and supplementary data were examined for information on locus identifiers, short and long forms of locus names, mutagenesis systems, loss-of-function phenotypes, and protein classes. The protein class of each entry was reconfirmed with the *M. truncatula* genome browser (Pecrix et al., 2018) and the UniProt reference protein database (UniProt Consortium, 2019). The correspondence between the latest locus identifiers and identifiers from earlier versions of the genome release was established in various ways, depending on the availability of the primary information in original studies. For some entries, the current locus identifiers were deduced using BLASTN (Altschul et al., 1990; Boratyn et al., 2013) analysis of primers and/or RNAi segments, BLASTP (Altschul et al., 1990; Boratyn et al., 2013) analysis of protein sequences, and sometimes required the conversion of images to text for the extraction of sequence information. All steps were manually curated, free of artificial intelligence.

### Detection of ncORFs

Conserved and MS-validated ncORFs were detected using MosaicProt (Çakır et al., 2025c). In short, each annotated transcript was scanned for the presence of stop-free regions longer than 59 nt in all forward reading frames. Such regions were *in silico* translated, supplied with unique informative identifiers, and saved in FASTA format. RefProts were separated from ncProts by matching each sequence to the currently annotated list of refProts.

For conservation analysis, all ncProts were compared with UniProt v. 2020_02 entries (UniProt Consortium, 2019) using DIAMOND v. 0.9.14 (Buchfink et al., 2021). NcProts with at least one match, including matches to *M. truncatula* sequences only, were categorized as conserved if the e-value of the top match was not more than 0.001. We explicitly declare that our definition of conserved ncProts is conditional in this study. It does not imply multi-species conservation, although some ncProts are conserved across hundreds of species.

For MS-validation of ncProts, they were compared to MS peptides from three publicly deposited proteomic datasets (PXD002692, Marx et al., 2016; PXD013606, Shin et al., 2021; and PXD022278, Castañeda et al., 2021), 16 samples in total, using SearchGUI v. 4.0.41 (Barsnes & Vaudel, 2018) and its partner tool PeptideShaker v. 2.0.33 (Vaudel et al., 2015). Each sample was searched independently using the target-decoy approach with FDR below 1%. MS peptides matching refProts and/or common contaminants were filtered out. The detection of NCR169 serves as a positive control for our procedure. NCR169 is a well-characterized peptide crucial for SNF (Starker et al., 2006; Domonkos et al., 2013; Farkas et al., 2014; Horváth et al., 2015, 2023; Güngör et al., 2023; Li et al., 2025).

MS-validated non-chimeric and chimeric ncProts (805 and 156, respectively) were mapped to the current genome and are available in the genome browser as two separate tracks (Çakır et al., 2025b). For chimeric MS peptides, not only 145 primary sources but also 101 potential alternative sources were mapped. The primary sources are loci in which chimeric MS peptides were identified in our analysis. Alternative sources of the same peptides were found using the ChiMSource software (Çakır et al., 2025a). Which of the loci are the true sources of chimeric MS-validated peptides was discussed in Çakır et al. (2025b). For 805 non-chimeric MS-validated peptides, we found 833 potential sources, which are mapped in the genome browser.

### Locating mutations and insertions relative to refORFs and ncORFs of affected genes

To determine whether refORFs and selected ncORFs were affected in the original studies, we analyzed images and sequence data presented by the authors. In some cases, the exact positions of *Tnt1* insertions were deduced from FST sequences downloaded from the *Tnt1* mutant database of *M. truncatula* (Tadege et al., 2008; Lee et al., 2018; Kaur et al., 2021). Exon-intron maps of gene models were prepared by aligning cDNA to genomic DNA in Geneious^®^ (MUSCLE alignment), with the manual correction of alignment artifacts at the splicing sites.

## Supporting information

Data S1 Master table non-SNF refORFs

Data S2 Master table non-SNF ncORFs

Data S3 Master table SNF refORFs

Data S4 Master table SNF ncORFs

Data S5 Common genes

Data S6 All genes

Data S7 References for LOF studies refORFs and ncORFs

Data S8 References for GOF studies ncORFs only

Data S9 Biological processes

Data S10 Protein classes individual groups

Data S11 Protein classes combined groups

Data S12 Master table loci with ncORFs

Data S13 Conservation at the aa level

Data S14 dNdS MEGA

Data S15 Short alignments

Data S16 Origin of conserved ncORFs

Data S17 Long alignments

Data S18 Gene and altORF models

Data S19 NcProts grouped with refProts during the MS analysis

Graphical abstract

Supporting Discussion

Supporting Results

## Abbreviations

ncORF: non-canonical open reading frame
refORF: reference open reading frame
ncProt: non-canonical protein
refProt: reference protein
mRNA: messenger
RNA: ncRNA, non-coding
RNA: rRNA, ribosomal
RNA: tRNA, transfer
RNA: CDS, coding sequence
UTR: untranslated region
Aa: amino acids
Bp: base pairs
Nt: nucleotides
MS: mass spectrometry

## AUTHOR CONTRIBUTIONS

**Umut Çakır**: Conceptualization, Methodology, Software, Validation, Formal analysis, Resources, Data Curation, Investigation, Writing - Original Draft, Writing - Review & Editing, Visualization, Funding acquisition. **Noujoud Gabed**: Conceptualization, Validation, Formal analysis, Resources, Investigation, Writing - Review & Editing. **Selen Kaya**: Investigation, Data Curation. **Vagner A. Benedito**: Conceptualization, Writing - Review & Editing. **Marie Brunet**: Methodology. **Xavier Roucou**: Methodology, Writing - Review & Editing. **Igor S. Kryvoruchko**: Conceptualization, Methodology, Validation, Formal analysis, Resources, Data Curation, Investigation, Software, Writing - Original Draft, Writing - Review & Editing, Visualization, Supervision, Project administration, Funding acquisition.

## ACKNOWLEDGMENTS

This work was supported by the Scientific and Technological Research Council of Turkey (TÜBİTAK) grants and Boğaziçi University standard research grant (BAP-P) to UÇ and IK (TÜBİTAK 1001 120Z514, TÜBİTAK 1002 120Z247, and BAP-P 18841), a Canada Research Chair in functional proteomics and discovery of novel proteins to XR, and a Canadian Institutes of Health Research Project Grant PJT-175322 to XR and MB. MB was supported by a Fonds de Recherche du Québec en Santé (FRQS) Junior 1 award (307936). Computational analysis was conducted using the server of the Turkish National e-Science e-Infrastructure (TRUBA) center. We sincerely thank Dr. Yun Kang for her critical feedback on the manuscript. The integration of the ncORF tracks and the loss-of-function track into the genome browser of *M. truncatula* would be impossible without continuous support from Dr. Sebastien Carrere. The completion of this study was made possible by support for IK from the United Arab Emirates University and for UÇ from the IMPRS-Genome Science PhD program.

## DATA AVAILABILITY STATEMENT

The annotated models of 123 genes listed in Data S12 are available in Geneious^®^ format in the online version of this article (Data S18). They include cDNA and gDNA sequences, refORFs and ncORFs mapped to cDNA sequences, and alignments of cDNA and gDNA, which outline exon-intron structures of the models. The entire set of files for the conservation evidence and MS-validation of ncProts will be published later in a paper describing all conserved and translated ncProts detected *in M. truncatula*. On 8 January 2025, our ncORF data were integrated into the *M. truncatula* genome portal MtrunA17r5.0-ANR as two separate tracks. In total, 156 MS-validated chimeric peptides were mapped to 246 primary and alternative source loci in the genome browser (non-repeat-element alternative sources only). In addition, 805 MS-validated non-chimeric ncProts were mapped to 833 genomic loci. On 10 April 2026, our loss-of-function dataset was integrated into the browser as a new track. It will be maintained and updated by the corresponding author. Users are encouraged to send their relevant data to IK in the format adopted in this article. Likewise, requests to update or edit the existing information will be considered. The complete set of data related to this article is available from the digital repository Zenodo (record 19398421). The codes used in this study are accessible from the following GitHub repository: github.com/umutcakir/ncORFs_in_studied_genes_of_Medicago_truncatula

## SUPPORTING INFORMATION

Additional Supporting Information may be found in the online version of this article.

**Figure S1**. Loss-of-function approaches used for studying 673 genes in *Medicago truncatula*.

**Figure S2**. Biological processes examined in loss-of-function studies on 673 genes in *Medicago truncatula*.

**Figure S3**. Different groups of protein classes have different proportions of phenotypes, which may reflect their relative functional importance.

**Figure S4**. Partitioning of phenomics data into altered (A), conditional (C), and neutral (N) phenotypes.

**Figure S5**. Phenotypes of 219 genes studied in the context of both symbiotic nitrogen fixation (SNF) and other processes.

**Figure S6**. Aberrant transcript forms recovered from two *Tnt1* mutant lines of *MtSWEET11* (Kryvoruchko et al., 2016).

**Figure S7**. Frequencies of aberrant transcript forms recovered from two *Tnt1* mutant lines shown in Figure S6.

**Figure S8**. A ClustalW protein alignment of the mutant version of *MtSWEET11* (Type X) with the wild-type version from ecotype R108 (the background of the *Tnt1* mutant population).

**Figure S9**. Biological and biosafety implications of induced insertional mutagenesis.

**Table S1**. Loci incorrectly named in the genome browser v. 5.1.9 (an extended version of Table 3).

**Table S2**. Five v4 gene models that may be correct without splitting.

**Table S3**. MS-validated ncORFs that introduce ambiguity into the interpretation of loss-of-function studies.

**Data S1**. A master table that summarizes loss-of-function studies on mRNA-refORFs and ncRNAs tested for phenotypes other than symbiotic nitrogen fixation.

**Data S2**. A master table that summarizes loss-of-function and gain-of-function studies on ncORFs tested for phenotypes other than symbiotic nitrogen fixation.

**Data S3**. A master table that summarizes loss-of-function studies on mRNA-refORFs and ncRNAs tested for symbiotic nitrogen fixation phenotypes.

**Data S4**. A master table that summarizes loss-of-function and gain-of-function studies on ncORFs tested for symbiotic nitrogen fixation phenotypes.

**Data S5**. A master table that summarizes loss-of-function studies on mRNA-refORFs, ncRNAs, and ncORFs tested for both symbiotic nitrogen fixation (SNF) and non-SNF phenotypes.

**Data S6**. A master table that summarizes loss-of-function studies on 673 loci in *Medicago truncatula*.

**Data S7**. A list of 565 loss-of-function studies on mRNA-refORFs, ncRNAs, and ncORFs in *Medicago truncatula*.

**Data S8**. A list of 37 gain-of-function and other non-mutant studies on ncORFs/peptides in *Medicago truncatula*.

**Data S9**. The complete list of 97 biological processes examined in loss-of-function studies on 673 genes in *Medicago truncatula*.

**Data S10**. The complete list of protein classes, corresponding phenotype counts, and statistical analysis of proportions.

**Data S11**. The complete list of combined protein groups, corresponding phenotype counts, and statistical analysis of proportions.

**Data S12**. A master table that summarizes various parameters of 148 ncORFs found in 123 genes targeted by functional analyses in *Medicago truncatula*.

**Data S13**. Visual demonstration of conservation at the amino acid level for eight selected ncORFs.

**Data S14**. Evidence for purifying selection of eight selected ncORFs.

**Data S15**. 24 nucleotide alignments that demonstrate evidence for purifying selection of eight selected ncORFs.

**Data S16**. Sequence analysis demonstrating the origin of highly conserved ncORFs in five selected genes.

**Data S17**. Nucleotide alignments used in Data S16.

**Data S18**. Gene models and mapped ncORFs of 123 loci targeted by functional analyses in *Medicago truncatula* (Data S12).

**Data S19**. NcProts grouped with refProts during the MS analysis.

