## Supplementary material for "Loss-of-function phenomics, ncORFs, and ambiguity of mutant phenotypes in *Medicago truncatula*": Data S13 Conservation at the aa level

**MtrunA17_Chr7g0273141**, MthGSHSb, putative homoglutathione synthase

**<u>NcORF11</u>**: MtrunA17_Chr7g0273141_2F_2-136_135

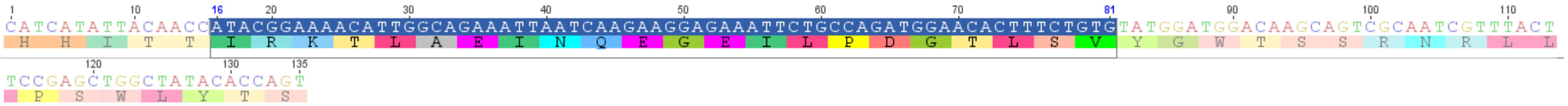


Species: *Phaseolus vulgaris*

Taxonomic group: eudicots, legumes

Annotation: homoglutathione synthetase (hgshs)

Protein ID: AAF98157.1

Nucleotide seq. ID: AF258320.1

**<u>Conservation</u>**:

amino acid level: 91%

nucleotide level: 89%

dN/dS: 0.111, p-value: 0.037

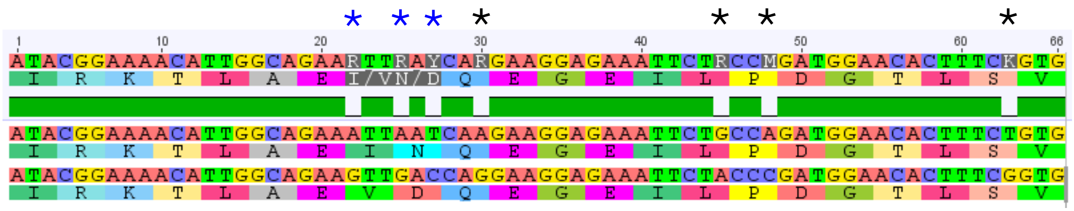


Consensus

*Mt*

*Pv*

**MtrunA17_Chr7g0273141**, MthGSHSb, putative homoglutathione synthase

**<u>NcORF11</u>**: MtrunA17_Chr7g0273141_2F_2-136_135

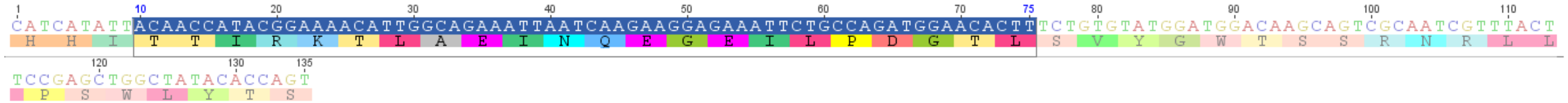

Species: *Salix viminalis*

Taxonomic group: eudicots, non-legumes

Annotation: glutathione synthetase

Protein ID: KAJ6737531.1

Nucleotide seq. ID: JAPFFL010000003.1

**<u>Conservation</u>**:

amino acid level: 91%

nucleotide level: 85%

dN/dS: 0.056, p-value: 0.018

Consensus

*Mt*

*Sv*

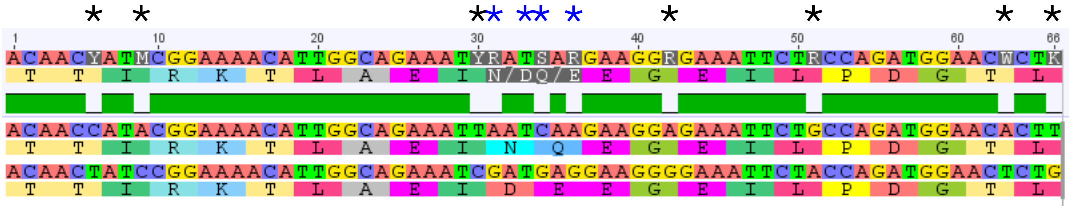

**MtrunA17_Chr7g0273141**, MthGSHSb, putative homoglutathione synthase

**NcORF11**: MtrunA17_Chr7g0273141_2F_2-136_135

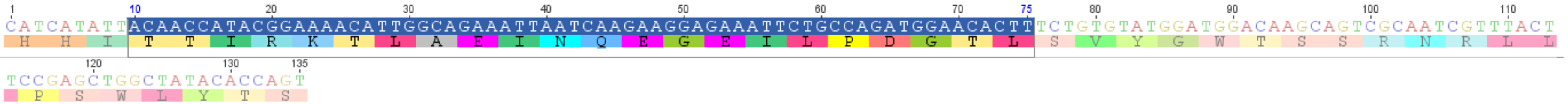


Species: *Populus trichocarpa*

Taxonomic group: eudicots, non-legumes

Annotation: glutathione synthetase, chloroplastic

Protein ID: XP_024456199.1

Nucleotide seq. ID: XM_024600431.2

**Conservation**:

amino acid level: 91%

nucleotide level: 83%

dN/dS: 0.045, p-value: 0.019

Consensus

*Mt*

*Pt*

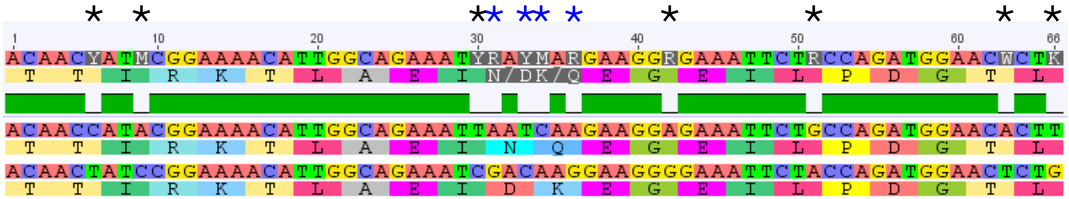

**MtrunA17_Chr7g0273141**, MthGSHSb, putative homoglutathione synthase

**<u>NcORF12</u>**: MtrunA17_Chr7g0273141_1F_625-816_192

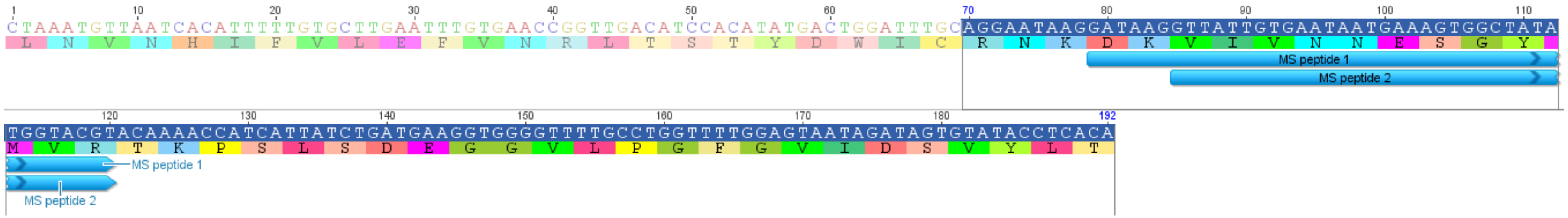


Species: *Cicer arietinum*

Taxonomic group: eudicots, legumes

Annotation: glutathione synthetase, chloroplastic-like isoform X1

Protein ID: XP_027188838.1

Nucleotide seq. ID: XM_027333037.2

**<u>Conservation</u>**:

amino acid level: 98%

nucleotide level: 93%

dN/dS: 0.051, p-value: 0.009

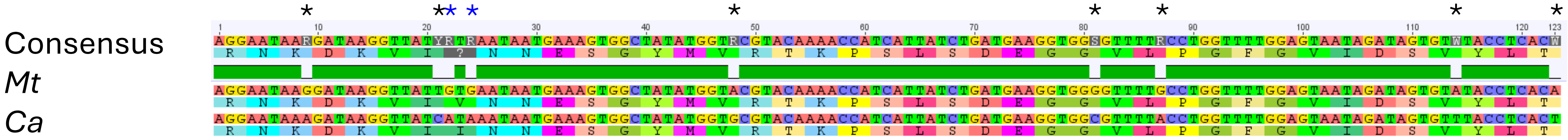

**MtrunA17_Chr7g0273141**, MthGSHSb, putative homoglutathione synthase

**NcORF12**: MtrunA17_Chr7g0273141_1F_625-816_192

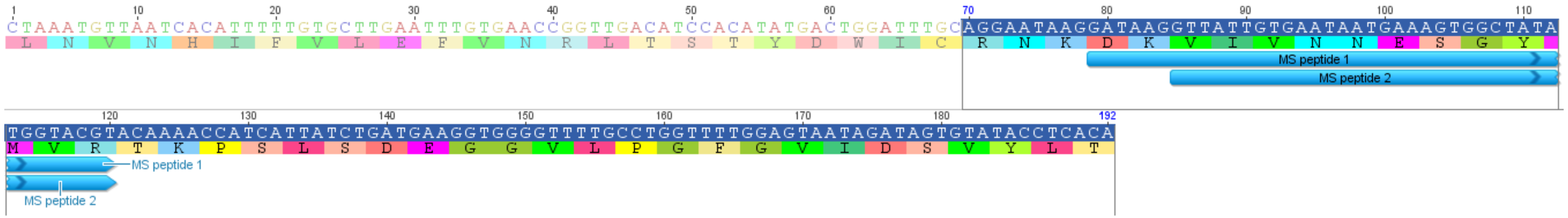


Species: *Cucumis sativus*

Taxonomic group: eudicots, non-legumes

Annotation: glutathione synthetase (GSH2)

Protein ID: ADM53349.1

Nucleotide seq. ID: HM230748.1

**Conservation**:

amino acid level: 71%

nucleotide level: 67%

dN/dS: n/c, p-value: n/c

Consensus

*Mt*

*Cs*

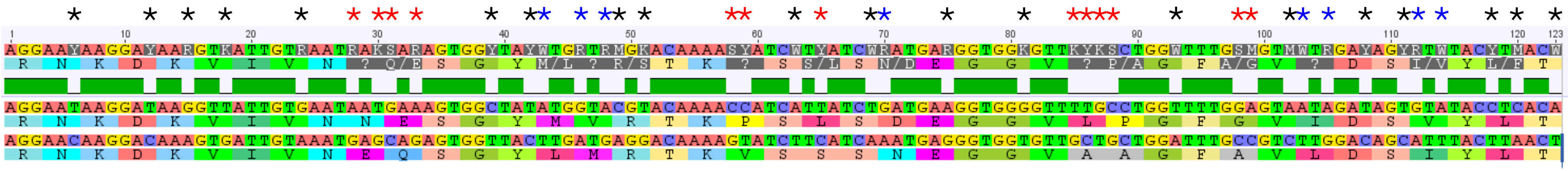

**MtrunA17_Chr4g0009054**, MtPHO2-A, ubiquitin-conjugating enzyme E2

**NcORF51**: MtrunA17_Chr4g0009054_1F_1300-1440_141

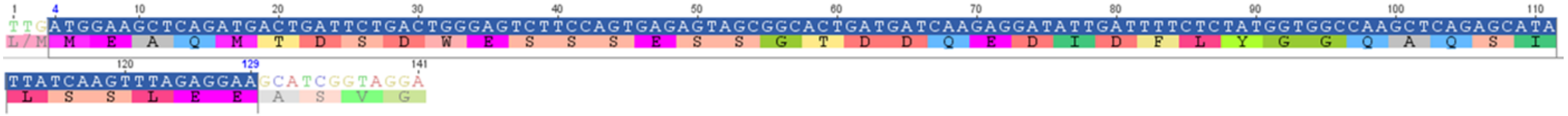

Species: *Vicia villosa*

Taxonomic group: eudicots, legumes

Annotation: probable ubiquitin-conjugating enzyme E2

Protein ID: XP_058771571.1

Nucleotide seq. ID: XM_058915588.1

**Conservation**:

amino acid level: 95%

nucleotide level: 90%

dN/dS: 0.025, p-value: 0.006

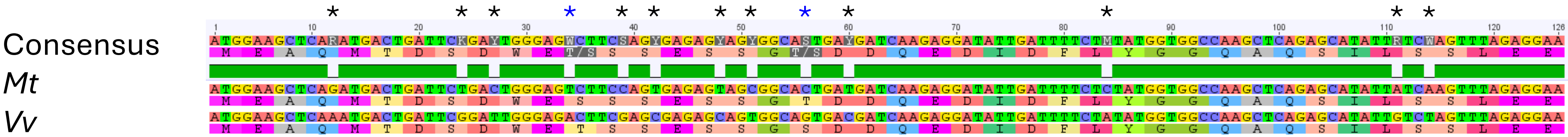

**MtrunA17_Chr4g0009054**, MtPHO2-A, ubiquitin-conjugating enzyme E2
**NcORF53**: MtrunA17_Chr4g0009054_1F_3370-3927_558

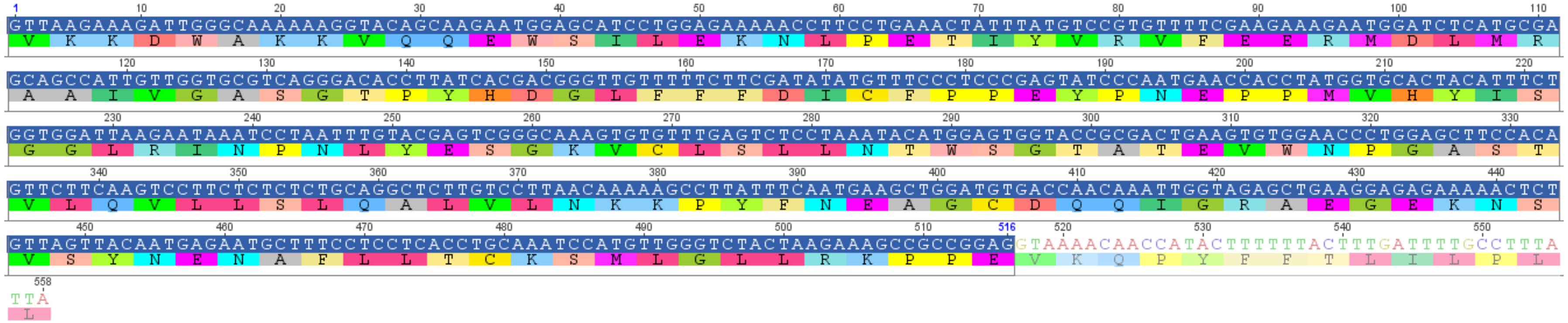

Species: *Cicer arietinum*
Taxonomic group: eudicots, legumes
Annotation: probable ubiquitin-conjugating enzyme E2
Protein ID: XP_004485781.1
Nucleotide seq. ID: XM_004485724.4
**Conservation**:
amino acid level: 96%
nucleotide level: 92%
dN/dS: 0.060, p-value: 1.110E-06

(continued on the next slide)

**MtrunA17_Chr4g0009054**, MtPHO2-A, ubiquitin-conjugating enzyme E2

**NcORF53**: MtrunA17_Chr4g0009054_1F_3370-3927_558

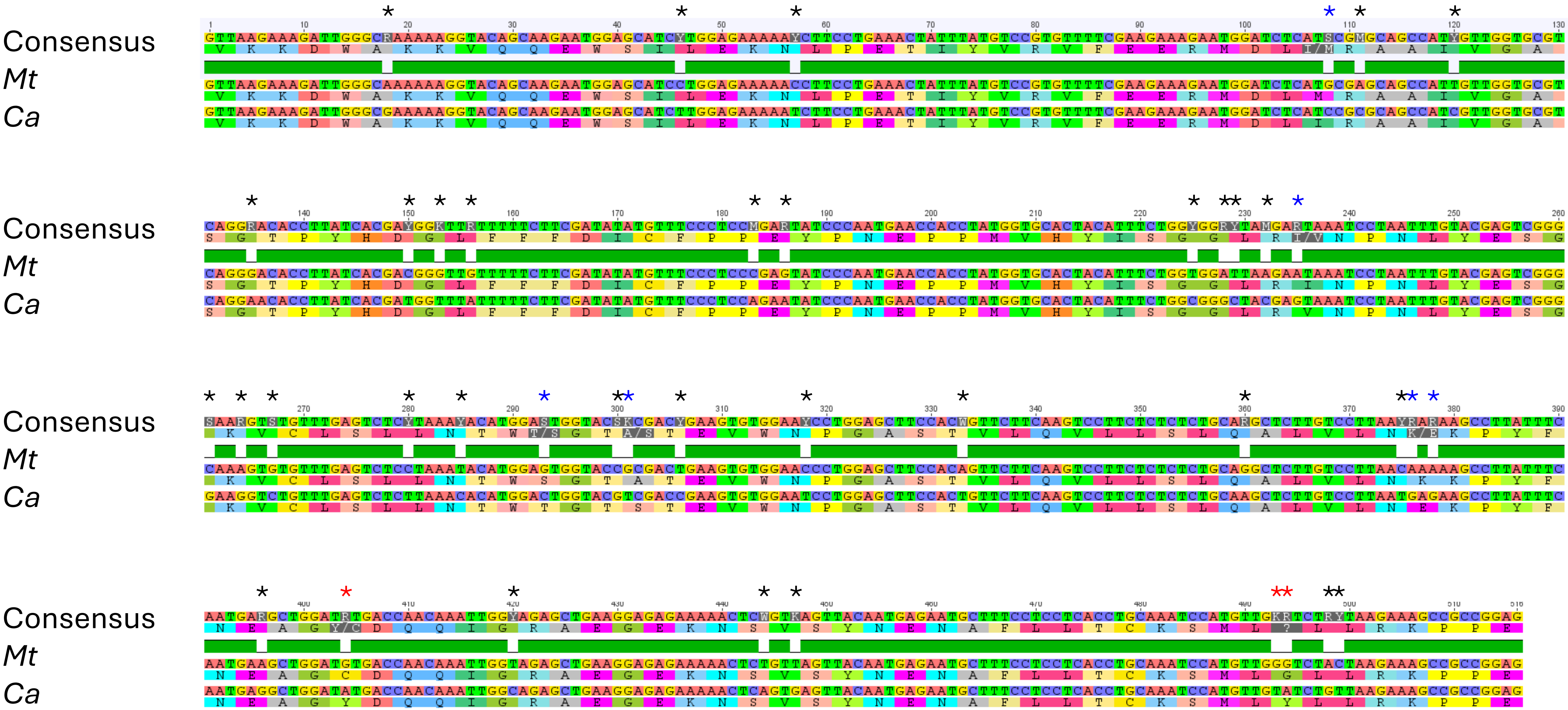

**MtrunA17_Chr4g0009054**, MtPHO2-A, ubiquitin-conjugating enzyme E2

**<u>NcORF53</u>**: MtrunA17_Chr4g0009054_1F_3370-3927_558

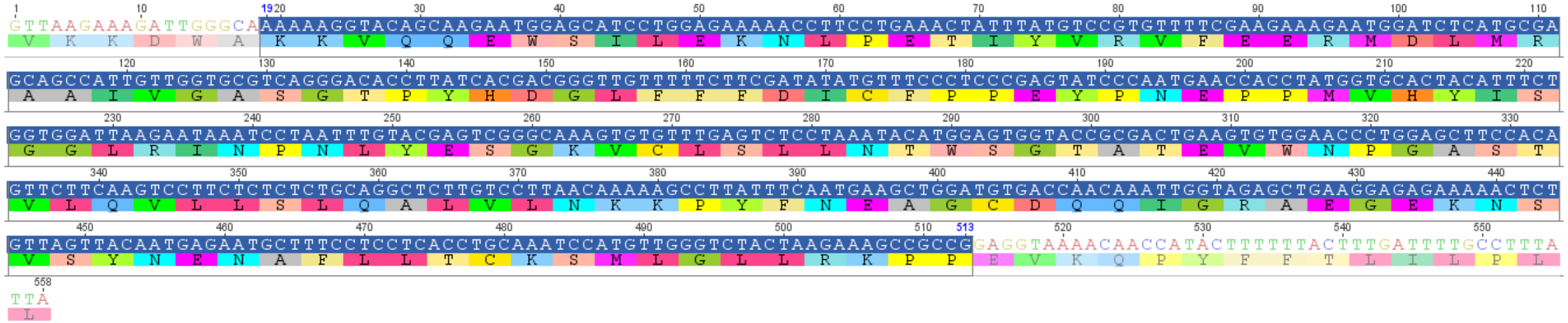

Species: *Morus notabilis*

Taxonomic group: eudicots, non-legumes

Annotation: probable ubiquitin-conjugating enzyme E2

Protein ID: XP_010108186.1

Nucleotide seq. ID: XM_010109884.2

**<u>Conservation</u>**:

amino acid level: 88%

nucleotide level: 77%

dN/dS: 0.029, p-value: 0.094

(continued on the next slide)

**MtrunA17_Chr4g0009054**, MtPHO2-A, ubiquitin-conjugating enzyme E2

**<u>NcORF53</u>**: MtrunA17_Chr4g0009054_1F_3370-3927_558

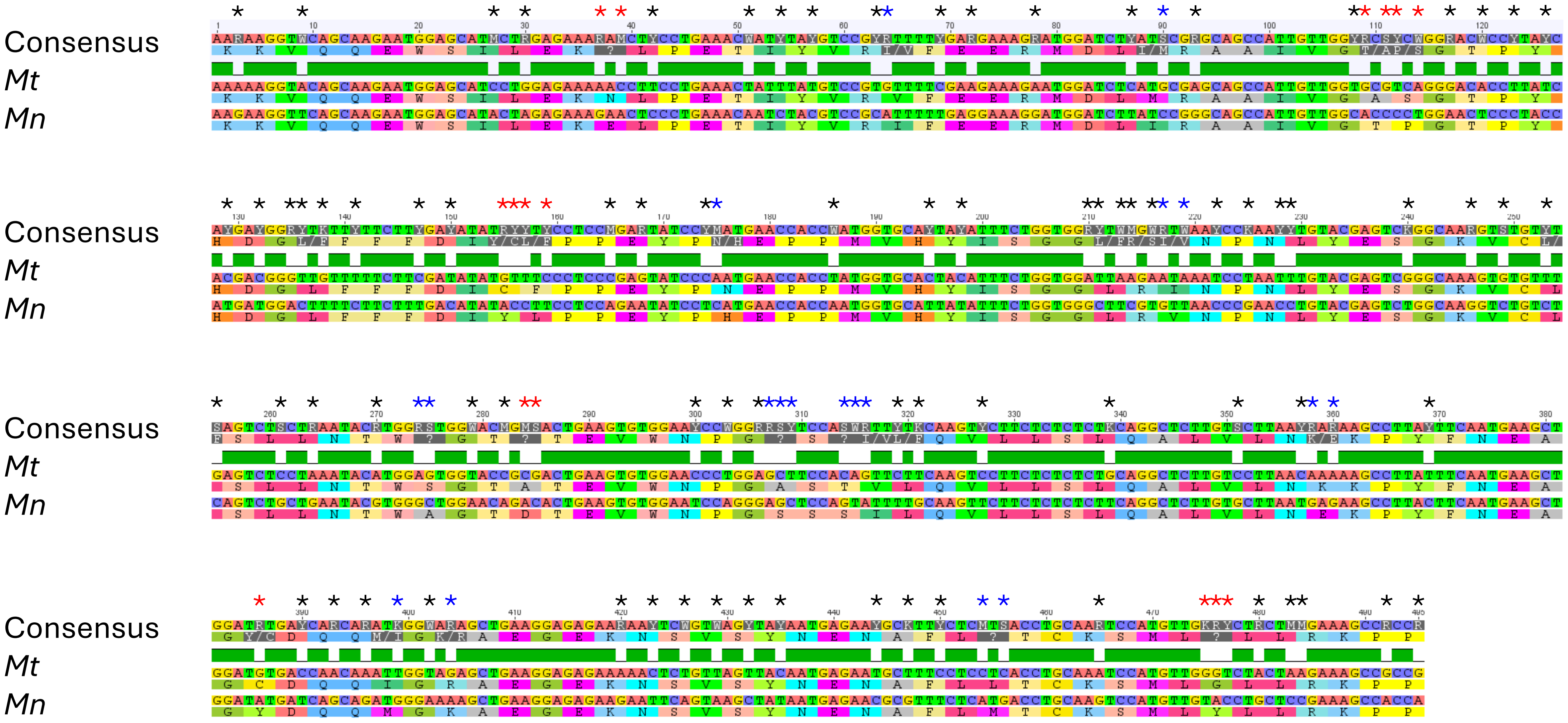

**MtrunA17_Chr4g0009054**, MtPHO2-A, ubiquitin-conjugating enzyme E2

**<u>NcORF53</u>**: MtrunA17_Chr4g0009054_1F_3370-3927_558

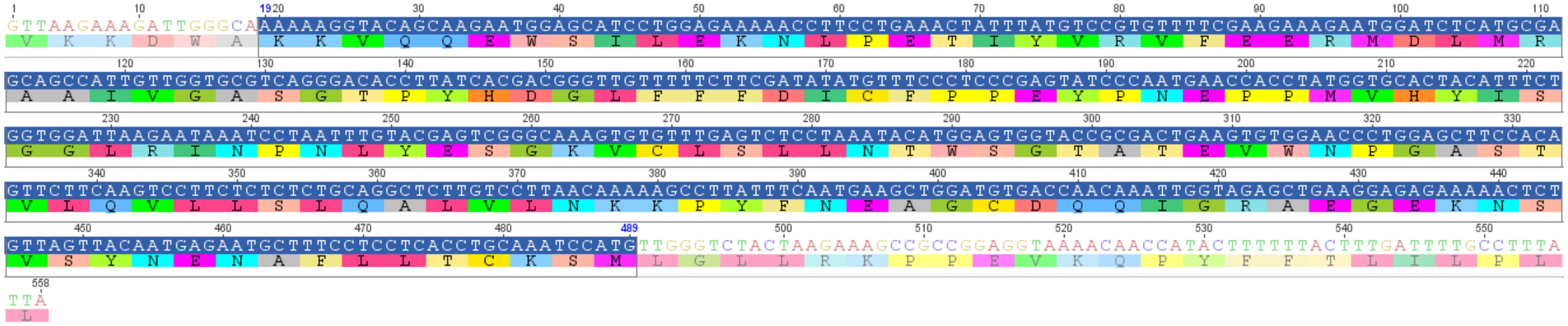

Species: *Rhynchospora pubera*

Taxonomic group: monocots

Annotation: ubiquitin-conjugating enzyme family protein

Protein ID: KAJ4731383.1

Nucleotide seq. ID: JAMFTS010007231.1

**<u>Conservation</u>**:

amino acid level: 82%

nucleotide level: 72%

dN/dS: n/c, p-value: n/c

(continued on the next slide)

**MtrunA17_Chr4g0009054**, MtPHO2-A, ubiquitin-conjugating enzyme E2

**<u>NcORF53</u>**: MtrunA17_Chr4g0009054_1F_3370-3927_558

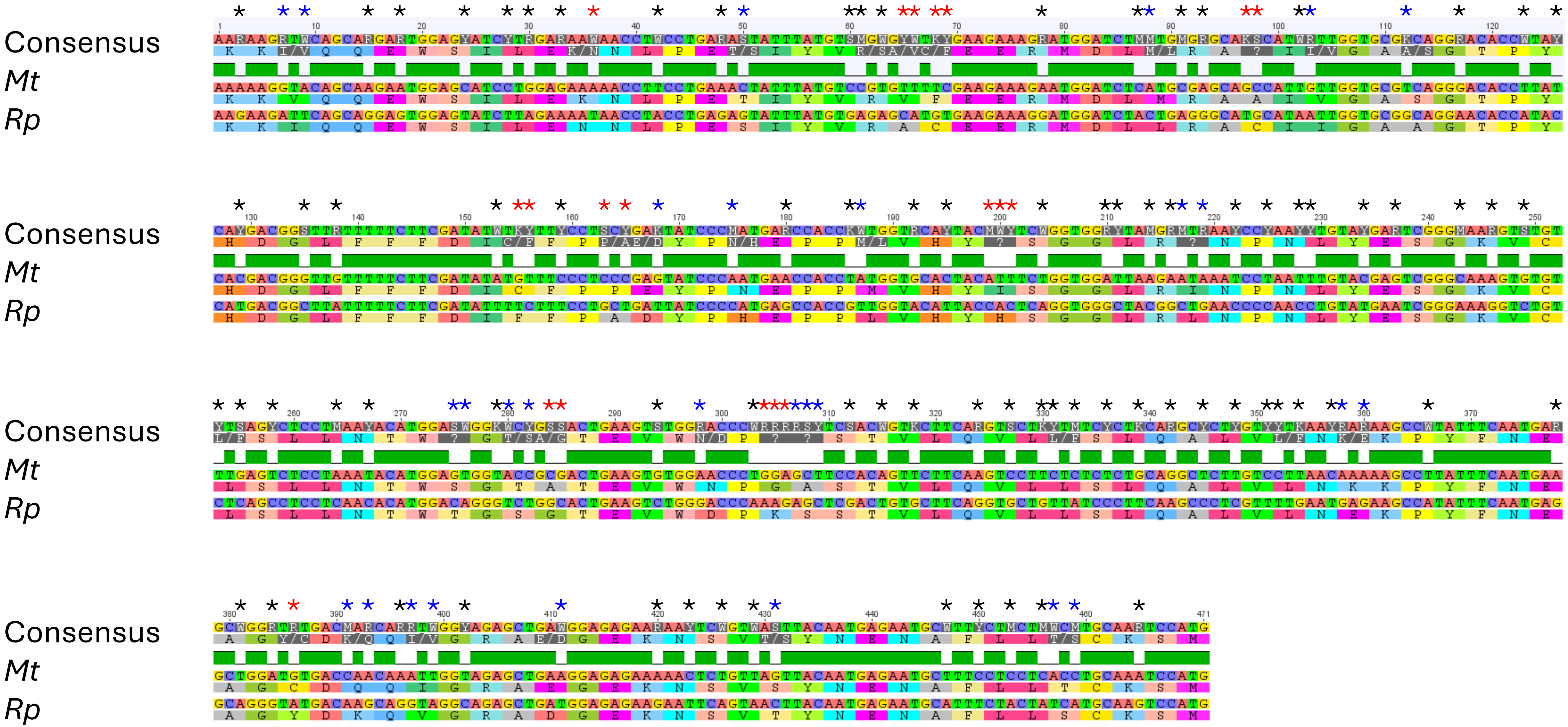

**MtrunA17_Chr4g0009054**, MtPHO2-A, ubiquitin-conjugating enzyme E2

**<u>NcORF54</u>**: MtrunA17_Chr4g0009054_2F_4052-4372_321

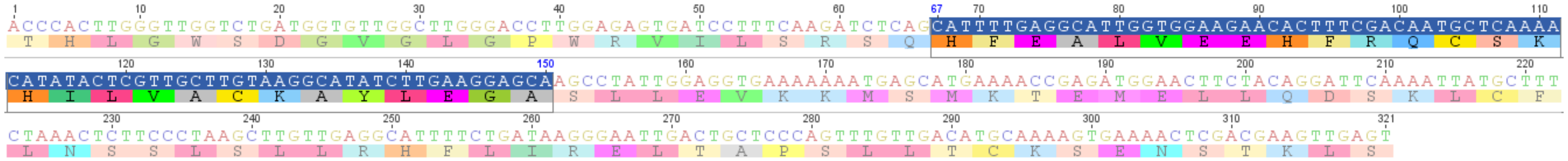

Species: *Glycine max*

Taxonomic group: eudicots, legumes

Annotation: probable ubiquitin-conjugating enzyme E2

Protein ID: XP_014621290.1

Nucleotide seq. ID: XM_014765804.3

**<u>Conservation</u>**:

amino acid level: 96%

nucleotide level: 92%

dN/dS: 0.061, p-value: 0.024

Consensus

*Mt*

*Gm*

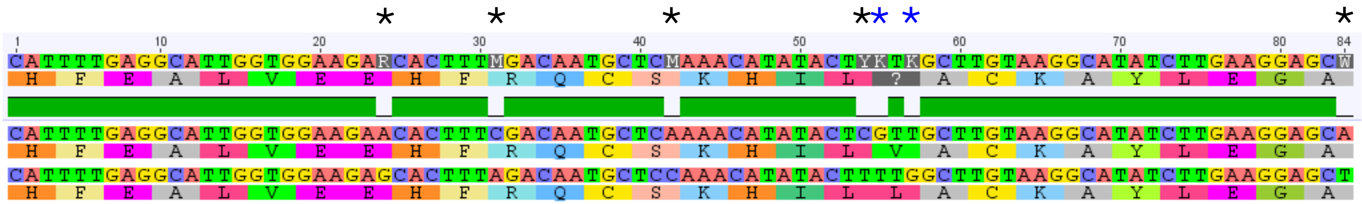

**MtrunA17_Chr4g0009054**, MtPHO2-A, ubiquitin-conjugating enzyme E2
**<u>NcORF54</u>**: MtrunA17_Chr4g0009054_2F_4052-4372_321

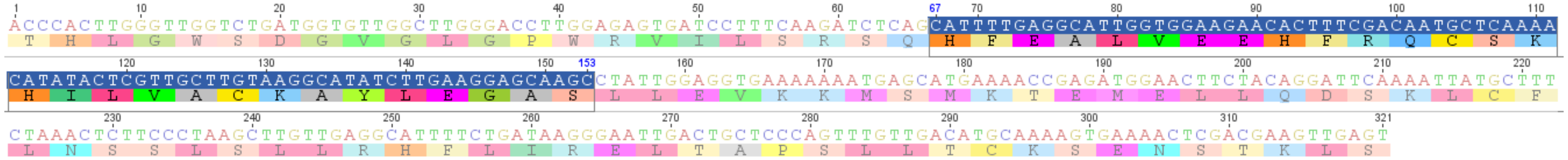

Species: *Alnus glutinosa*
Taxonomic group: eudicots, non-legumes
Annotation: probable ubiquitin-conjugating enzyme E2
Protein ID: XP_062161121.1
Nucleotide seq. ID: XM_062305137.1
**<u>Conservation</u>**:
amino acid level: 83%
nucleotide level: 75%
dN/dS: 0.057, p-value: 0.175

Consensus
*Mt*
*Ag*

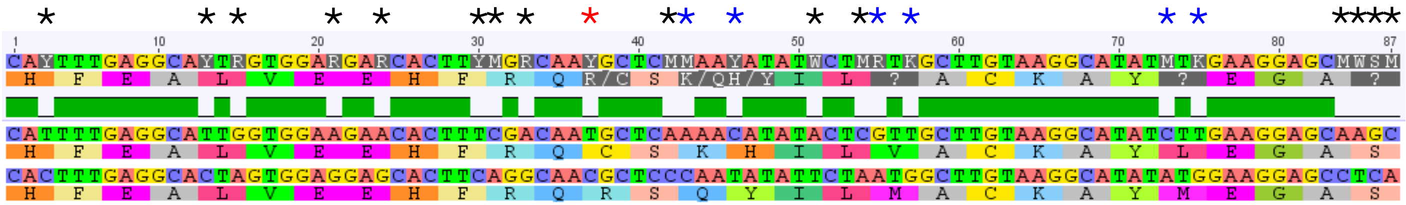

**MtrunA17_Chr4g0009054**, MtPHO2-A, ubiquitin-conjugating enzyme E2
**NcORF54**: MtrunA17_Chr4g0009054_2F_4052-4372_321

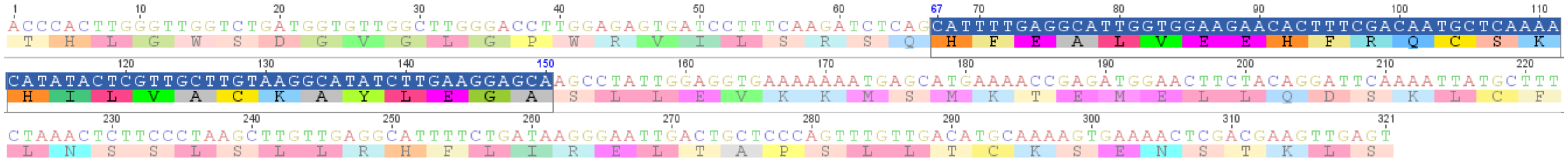

Species: *Elaeis guineensis*
Taxonomic group: monocots
Annotation: probable ubiquitin-conjugating enzyme E2
Protein ID: XP_073115209.1
Nucleotide seq. ID: XM_073259108.1
**Conservation**:
amino acid level: 82%
nucleotide level: 69%
dN/dS: n/c, p-value: n/c

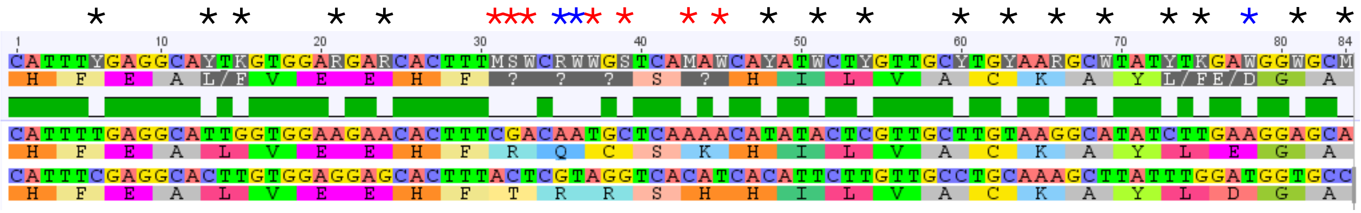

Consensus
*Mt*
*Eg*

**MtrunA17_Chr4g0009054**, MtPHO2-A, ubiquitin-conjugating enzyme E2
**NcORF55**: MtrunA17_Chr4g0009054_1F_4195-4350_156

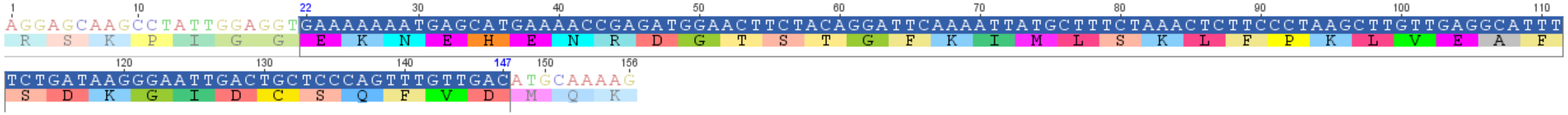

Species: *Glycine max*
Taxonomic group: eudicots, legumes
Annotation: putative ubiquitin-conjugating enzyme E2
Protein ID: KAH1242978.1
Nucleotide seq. ID: JAGRRG010000007.1
**Conservation**:
amino acid level: 93%
nucleotide level: 87%
dN/dS: 0.096, p-value: 0.007

Consensus
*Mt*
*Gm*

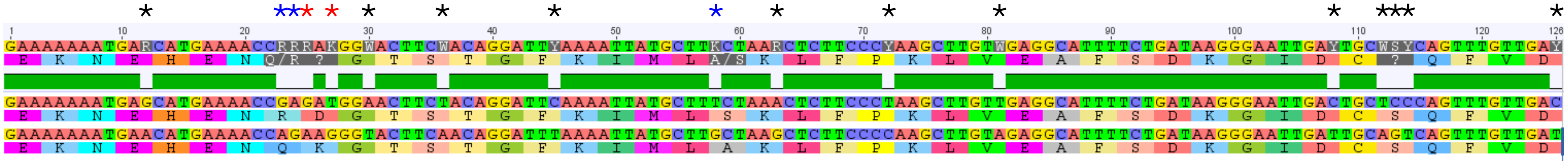

**MtrunA17_Chr4g0009054**, MtPHO2-A, ubiquitin-conjugating enzyme E2
**NcORF55**: MtrunA17_Chr4g0009054_1F_4195-4350_156

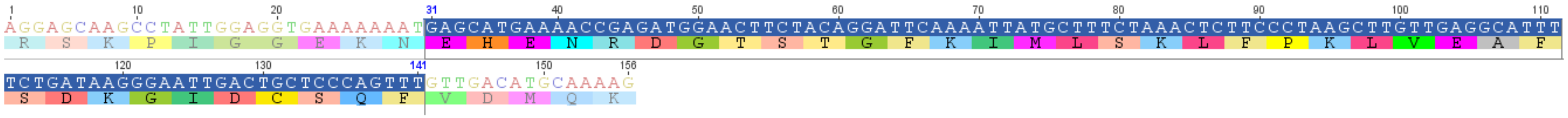

Species: *Quillaja saponaria*
Taxonomic group: eudicots, non-legumes
Annotation: ubiquitin-conjugating enzyme
Protein ID: KAJ7968561.1
Nucleotide seq. ID: JARAOO010000005.1
**Conservation**:
amino acid level: 92%
nucleotide level: 84%
dN/dS: 0.058, p-value: 0.009

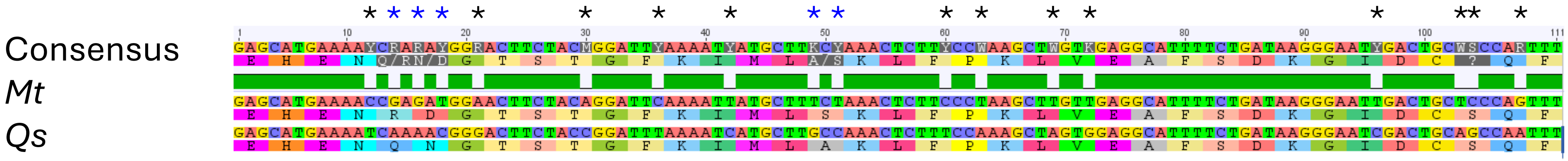

**MtrunA17_Chr4g0009054**, MtPHO2-A, ubiquitin-conjugating enzyme E2

**<u>NcORF55</u>**: MtrunA17_Chr4g0009054_1F_4195-4350_156

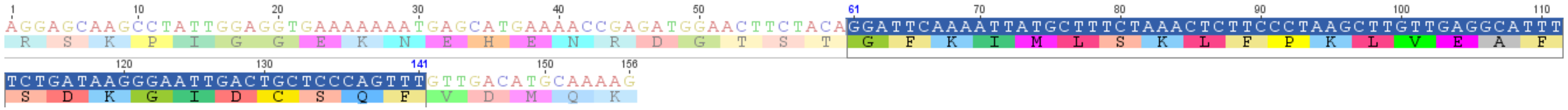

Species: *Nelumbo nucifera*

Taxonomic group: basal eudicots

Annotation: probable ubiquitin-conjugating enzyme E2

Protein ID: XP_010260198.1

Nucleotide seq. ID: XM_010261896.2

**<u>Conservation</u>**:

amino acid level: 85%

nucleotide level: 79%

dN/dS: 0.099, p-value: 0.041

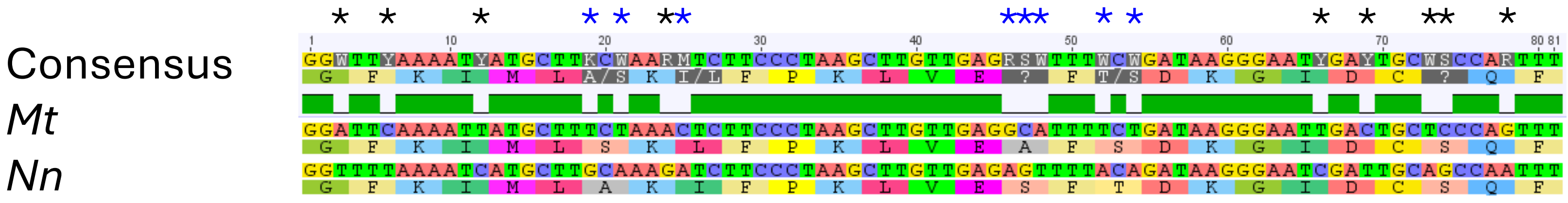

**MtrunA17_Chr4g0035414**, MtbZIP60b, S class bZIP transcription factor
**NcORF60**: MtrunA17_Chr4g0035414_3F_168-335_168

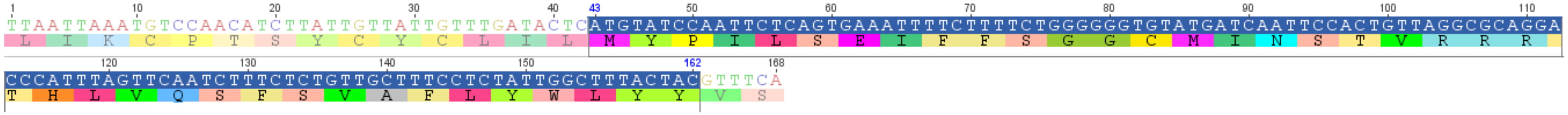

Species: *Glycine max*
Taxonomic group: eudicots, legumes
Annotation: bZIP transcription factor
Protein ID: KAH1220101.1
Nucleotide seq. ID: JAGRRG010000012.1
**Conservation**:
amino acid level: 98%
nucleotide level: 92%
dN/dS: 0.043, p-value: 0.015

Consensus
*Mt*
*Gm*

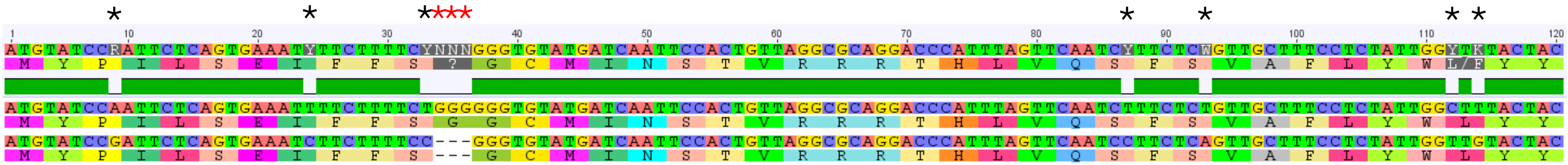

**MtrunA17_Chr4g0035414**, MtbZIP60b, S class bZIP transcription factor
**NcORF60**: MtrunA17_Chr4g0035414_3F_168-335_168

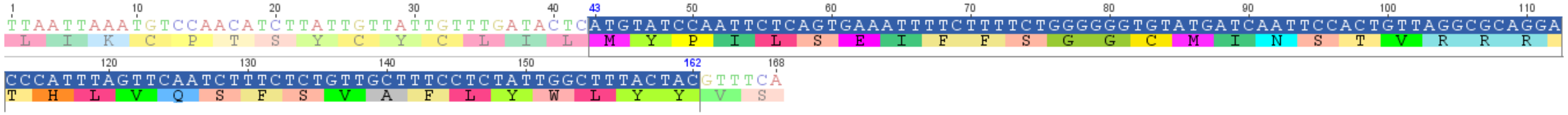

Species: *Mucuna pruriens*
Taxonomic group: eudicots, legumes
Annotation: bZIP transcription factor
Protein ID: RDY01331.1
Nucleotide seq. ID: QJKJ01002791.1
**Conservation**:
amino acid level: 98%
nucleotide level: 91%
dN/dS: 0.036, p-value: 0.010

Consensus
*Mt*
*Mp*

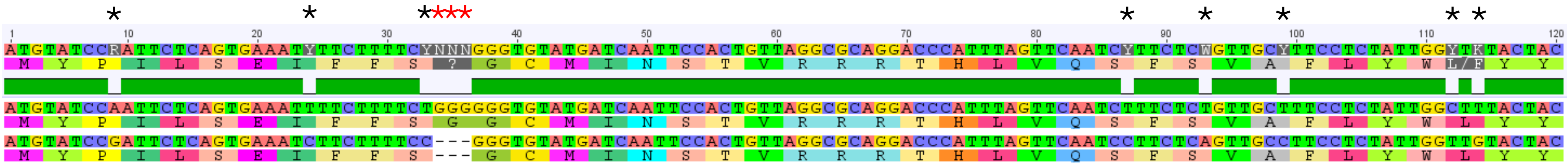

**MtrunA17_Chr4g0035414**, MtbZIP60b, S class bZIP transcription factor

**<u>NcORF60</u>**: MtrunA17_Chr4g0035414_3F_168-335_168

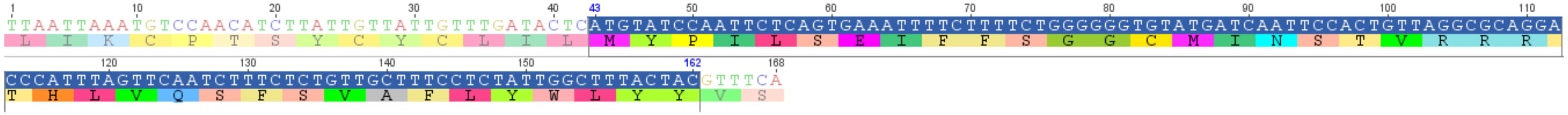

Species: *Lupinus albus*

Taxonomic group: eudicots, legumes

Annotation: putative transcription factor bZIP family

Protein ID: KAE9609489.1

Nucleotide seq. ID: WOCE01000008.1

**<u>Conservation</u>**:

amino acid level: 90%

nucleotide level: 85%

dN/dS: 0.099, p-value: 0.010

Consensus

*Mt*

*La*

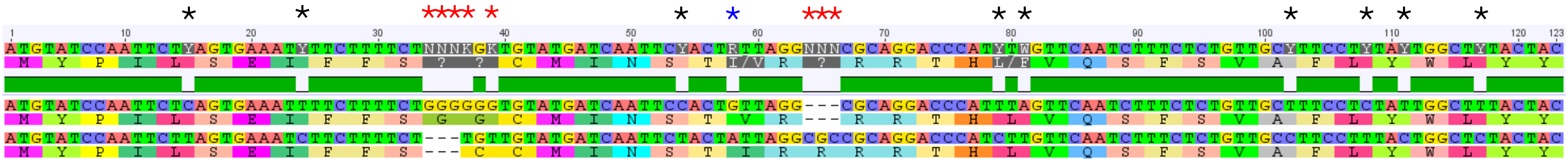

**MtrunA17_Chr4g0052121**, MtKAS II, ketoacyl-ACP synthase II
**NcORF65**: MtrunA17_Chr4g0052121_1F_982-1161_180

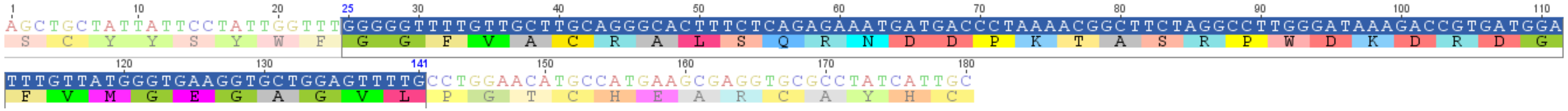

Species: *Glycine soja*
Taxonomic group: eudicots, legumes
Annotation: 3-oxoacyl-[acyl-carrier-protein] synthase I, chloroplastic
Protein ID: RZC12225.1
Nucleotide seq. ID: QZWG01000005.1
**Conservation**:
amino acid level: 100%
nucleotide level: 91%
dN/dS: 0.000, p-value: 0.003

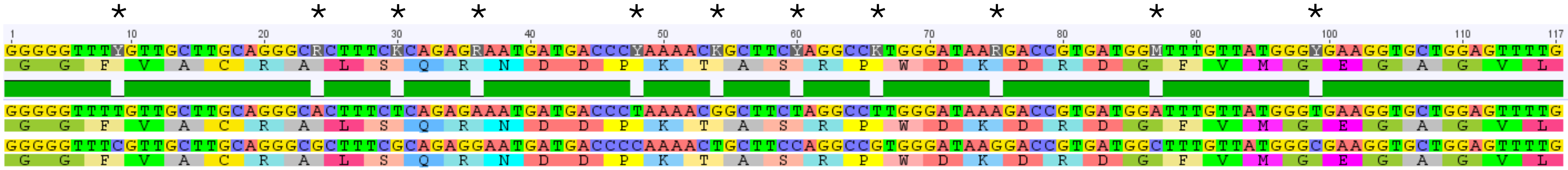

**MtrunA17_Chr4g0052121**, MtKAS II, ketoacyl-ACP synthase II
**NcORF65**: MtrunA17_Chr4g0052121_1F_982-1161_180

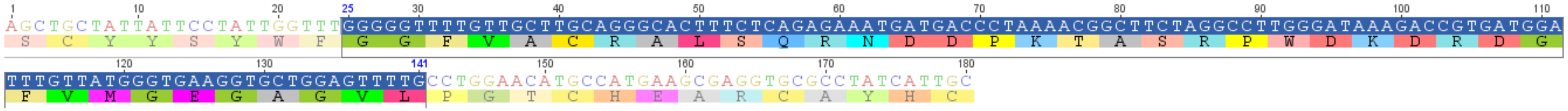

Species: *Cephalotus follicularis*
Taxonomic group: eudicots, non-legumes
Annotation: ketoacyl-synt domain-containing protein/MCM domain-containing protein
Protein ID: GAV66455.1
Nucleotide seq. ID: BDDD01000481.1
**Conservation**:
amino acid level: 100%
nucleotide level: 85%
dN/dS: 0.000, p-value: 0.013

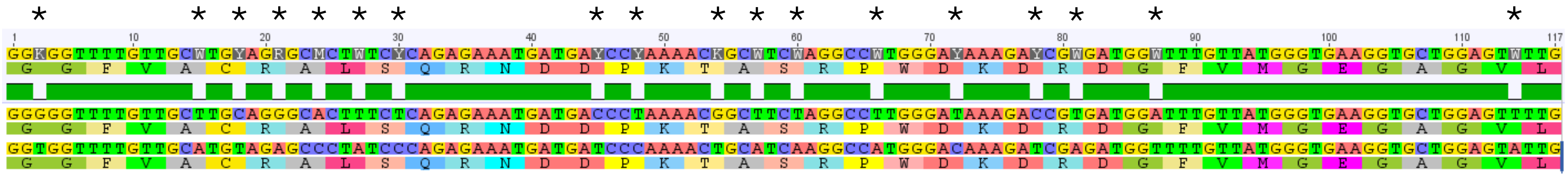

**MtrunA17_Chr4g0052121**, MtKAS II, ketoacyl-ACP synthase II
**<u>NcORF65</u>**: MtrunA17_Chr4g0052121_1F_982-1161_180

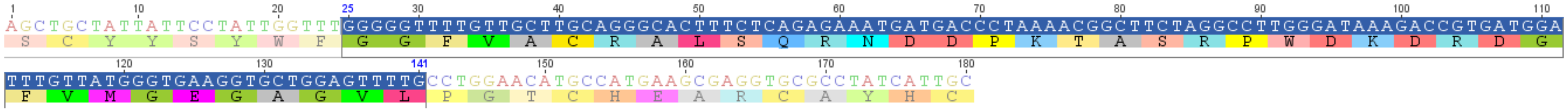

Species: *Rhynchospora pubera*
Taxonomic group: monocots
Annotation: 3-oxoacyl-[acyl-carrier-protein] synthase
Protein ID: KAJ4754275.1
Nucleotide seq. ID: JAMFTS010000005.1
**<u>Conservation</u>**:
amino acid level: 100%
nucleotide level: 86%
dN/dS: 0.000, p-value: 0.004

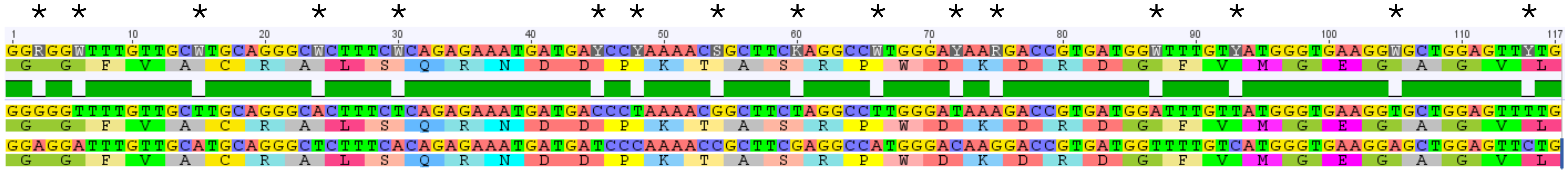

**MtrunA17_Chr4g0052121**, MtKAS II, ketoacyl-ACP synthase II

**NcORF65**: MtrunA17_Chr4g0052121_1F_982-1161_180

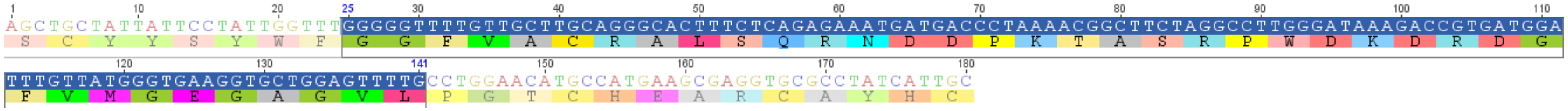

Species: *Magnolia sinica*

Taxonomic group: magnoliids

Annotation: 3-oxoacyl-[acyl-carrier-protein] synthase I, chloroplastic

Protein ID: XP_058109397.1

Nucleotide seq. ID: XM_058253414.1

**Conservation**:

amino acid level: 97%

nucleotide level: 85%

dN/dS: 0.009, p-value: 0.007

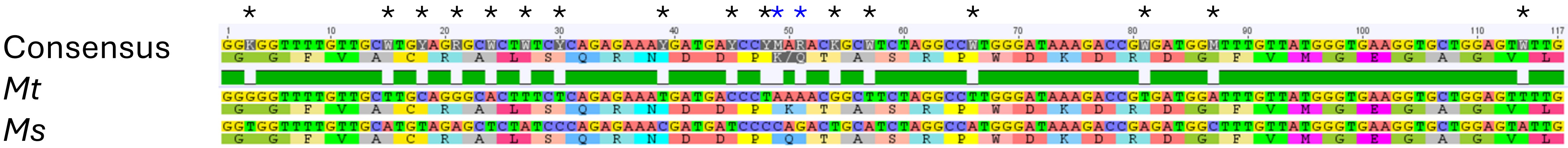

**MtrunA17_Chr4g0052121**, MtKAS II, ketoacyl-ACP synthase II

**<u>NcORF65</u>**: MtrunA17_Chr4g0052121_1F_982-1161_180

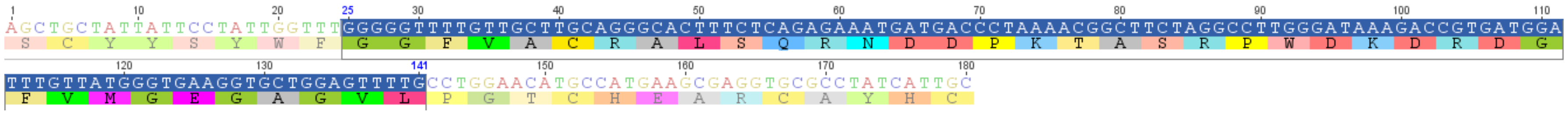

Species: *Physcomitrium patens*

Taxonomic group: bryophytes

Annotation: 3-oxoacyl-[acyl-carrier-protein] synthase II, chloroplastic

Protein ID: XP_024391557.1

Nucleotide seq. ID: XM_024535789.2

**<u>Conservation</u>**:

amino acid level: 95%

nucleotide level: 73%

dN/dS: n/c, p-value: n/c

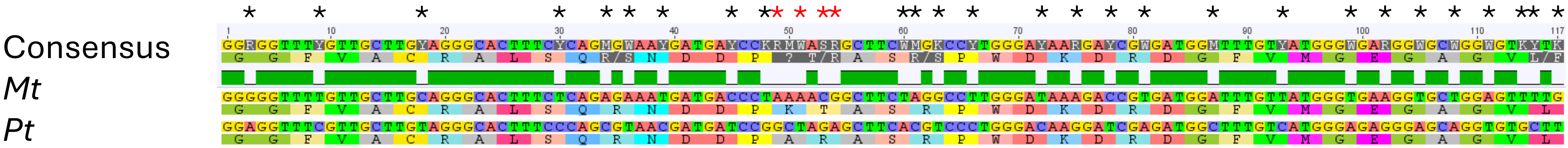

**MtrunA17_Chr4g0052121**, MtKAS II, ketoacyl-ACP synthase II

**<u>NcORF65</u>**: MtrunA17_Chr4g0052121_1F_982-1161_180

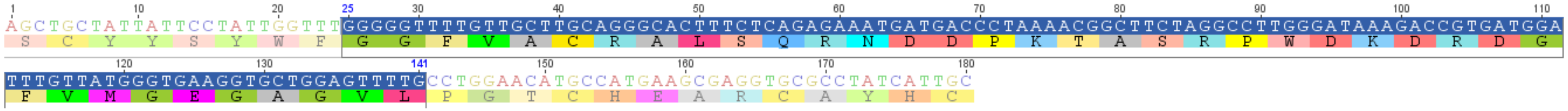

Species: *Pseudomonadota bacterium*

Taxonomic group: bacteria

Annotation: beta-ketoacyl-ACP synthase II

Protein ID: MFZ5502609.1

Nucleotide seq. ID: JBMOPZ010000030.1

**<u>Conservation</u>**:

amino acid level: 95%

nucleotide level: 68%

dN/dS: n/c, p-value: n/c

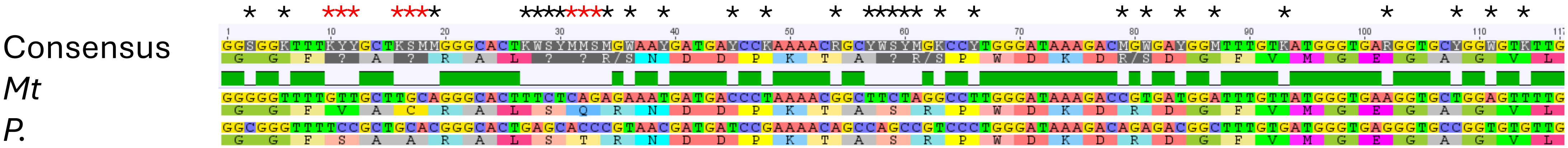


**Note**: despite the difference in the codon tables between plants and bacteria, amino acid sequences shown in the alignment using the standard codon table correspond to the actual amino acid sequences of corresponding source organisms. No dN/dS analysis is possible across different tables.

**Data S13**. Visual demonstration of conservation at the amino acid level for eight selected ncORFs. The dataset shows evidence for purifying selection (dN/dS ratios and their statistical significance computed in MEGA-X v10.2.6). Details of the calculations can be found in Data S14. Corresponding alignments in MEGA format can be extracted from Data S15). Color of asterisks shown above the alignments: black – a synonymous mutation; blue – a non-synonymous mutation that preserves properties of an amino acid in the chain (corresponds to a plus-sign in NCBI-BLASTN alignments); red – a non-synonymous mutation that does not preserve properties of an amino acid in the chain (corresponds to an empty space in NCBI-BLASTN alignments). Graphical data were generated in Geneious® R7.1.9. P-values are from Z-test for significance of dN/dS deviation from one. Only p-values without Bonferroni adjustment are shown. P-values after the adjustment are above 0.05, except for Slide 7. Even very strong conservation signatures obvious in some of the alignments in this dataset may not signify translation of a ncORF if it originates from a very recent frameshift that had no immediate effect on fitness (see Discussion).

**Contents**

Slides 1-5: *MthGSHSb*
Slides 6-18: *MtPHO2-A*
Slides 19-21: *MtbZIP60b*
Slides 22-27: *MtKASII*
