## Supplementary material for "Loss-of-function phenomics, ncORFs, and ambiguity of mutant phenotypes in *Medicago truncatula*": Data S15 Short alignments: Legend.docx

**Data S15**. 24 nucleotide alignments that demonstrate evidence for purifying selection of eight selected ncORFs. The alignments were constructed in Geneious® R7.1.9 using the MUSCLE tool and exported in MEGA format. Alignment names are the same as in Data S14.
