## Supplementary material for "Loss-of-function phenomics, ncORFs, and ambiguity of mutant phenotypes in *Medicago truncatula*": Data S16 Origin of conserved ncORFs

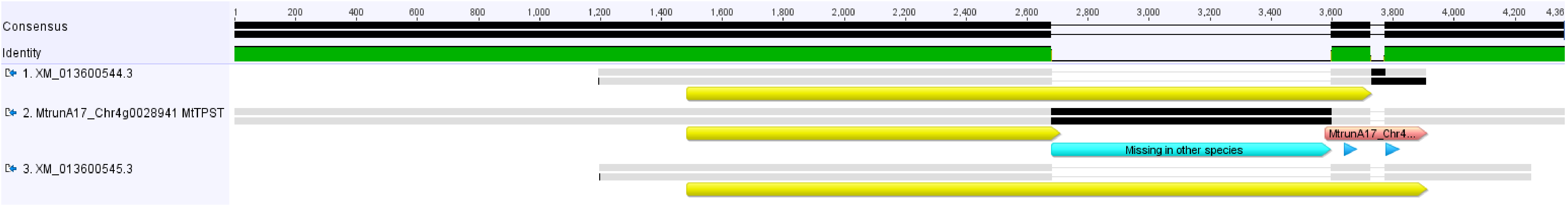


**Slide 1**. A MUSCLE alignment of the *MtTPST* cDNA and two putative homologs/sequence variants from *Medicago truncatula*. Yellow and pink arrow bars indicate refORFs and ncORFs, respectively. A cyan arrow bar indicates a region missing or heavily modified in most of legume species (see Slide 2). Blue arrows indicate positions of two MS peptides, SLNDLDLELYEYAR and WDNISSSEYVIIMWK, that we identified in a proteomic sample of 10-day-old root nodules in study PXD002692 (Marx et al., 2016). They are unique to ncProt6 (MtrunA17_Chr4g0028941_1F_3577-3867_291) and two refProts, XM_013600544.3 and XM_013600545.3. The complete absence of neutral polymorphic sites within the refORFs of these sequences indicates their origin from a single genetic locus. XM_013600544.3 and XM_013600545.3 have no corresponding annotated loci in the current version of the *M. truncatula* genome (v. 5.1.9). There are no other high-scoring *M. truncatula* subject sequences with similarity to *MtTPST* in the BLASTN search using the NCBI non-redundant nucleotide database (as of 25 February 2026).

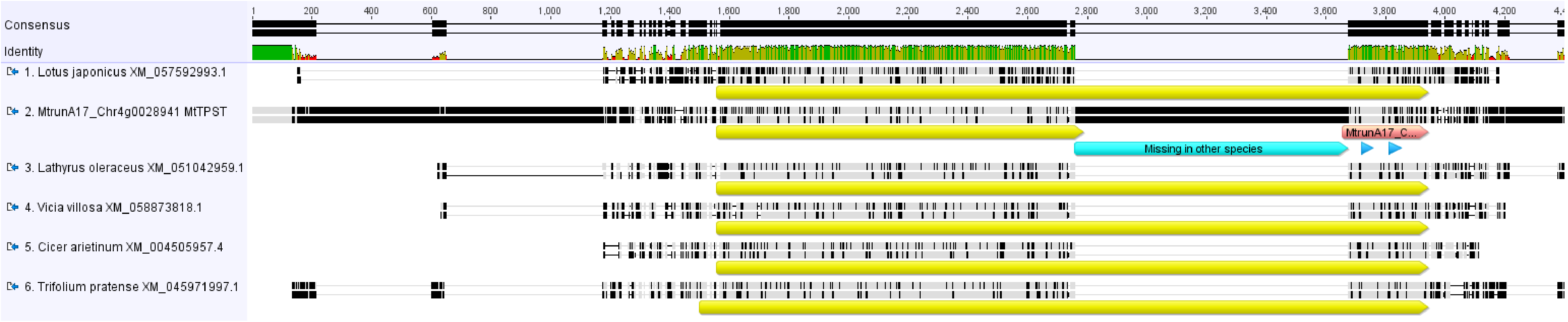


**Slide 2**. A MUSCLE alignment of the *MtTPST* cDNA and five putative homologs from other legumes species. Yellow and pink arrow bars indicate refORFs and ncORFs, respectively. Blue arrows indicate positions of two MS peptides, SLNDLDLELYEYAR and WDNISSSEYVIIMWK (see Slide 1). NcORF6 in *M. truncatula* corresponds to the end of refORF sequences in other legume species. It was disconnected from the refORF through an early in-frame stop codon but kept in the same reading frame by a putative insertion of 915 base pairs (cyan arrow bar), which is present in a slightly modified form in *M. arabica* and *M. lupulina* (93 and 94% identity, respectively) but has limited query coverage (9-70%) and similarity (77-92%) to other sequences in GenBank (as of 25 February 2026). This means that ncORF6 is either translated as a discrete unit or as a C-terminal part of a refProt, which corresponds to one valid gene model missed in the current version of the *M. truncatula* genome (v. 5.1.9). NcORF6 is conserved but is not under purifying selection, which indicates that its amino acid sequence is repurposed compared to the C-terminal refProt portions of other species.

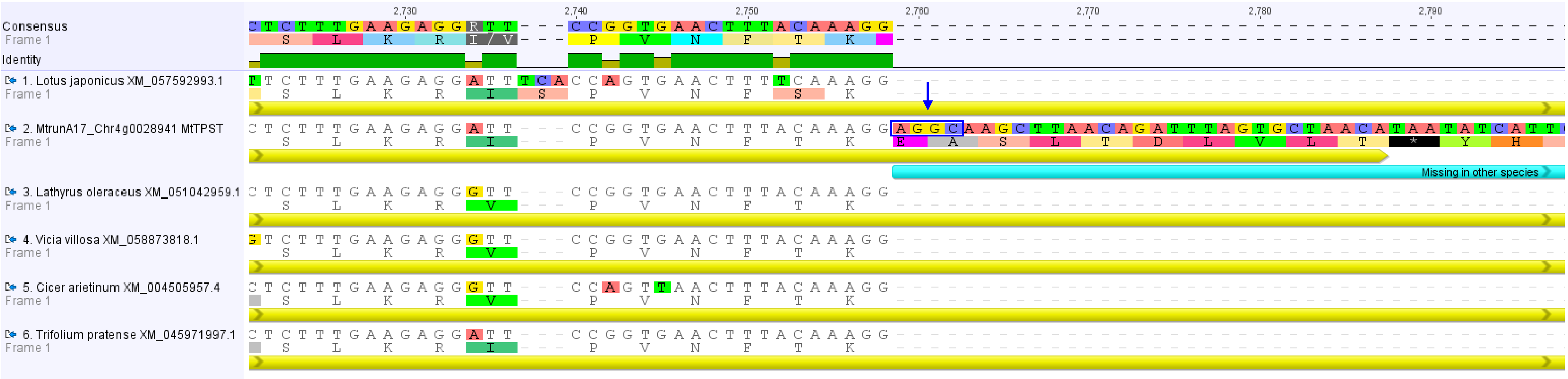


**Slide 3**. A fragment of the MUSCLE alignment shown in Slide 2 focusing on the left side of the 915-base-pair-long insertion. Because of a sequence AGGC (blue box and vertical arrow), which is very similar to the classical exon-intron border motif, the insertion is likely to act as a retained intron in the actual gene model, which means there must be an alternative splicing form in which this intron is removed. The same motif is found in the right side of this insertion (see Slide 4). Translation is shown relative to the refORFs of all six sequences.

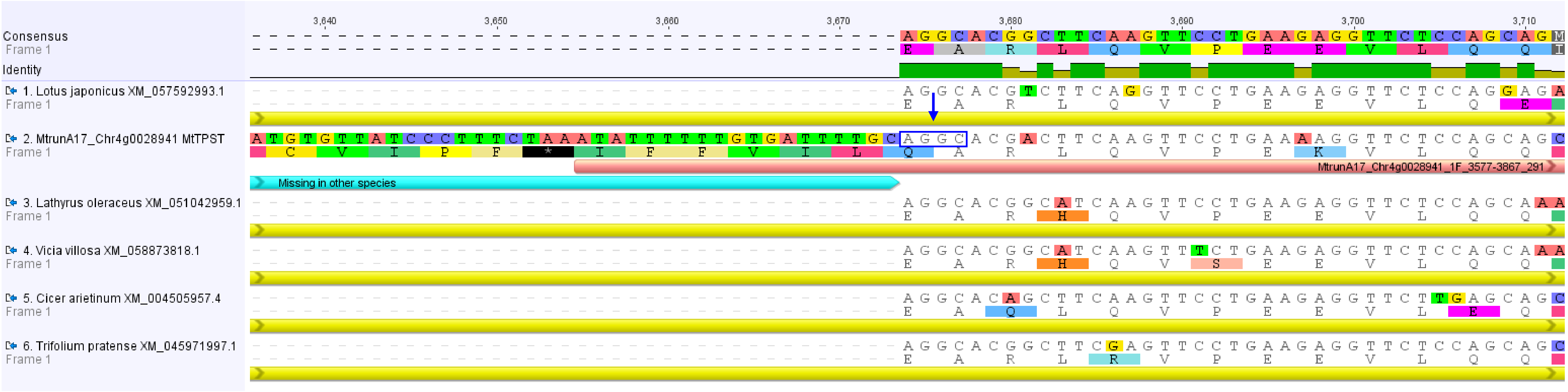


**Slide 4**. A fragment of the MUSCLE alignment shown in Slide 2 focusing on the right side of the 915-base-pair-long insertion. Because of the presence of a motif AGGC (blue box and vertical arrow), the insertion is likely to act as a retained intron in the current gene model, which means there must be an alternative splicing form in which this intron is removed. Translation is shown relative to the refORFs of all six sequences.

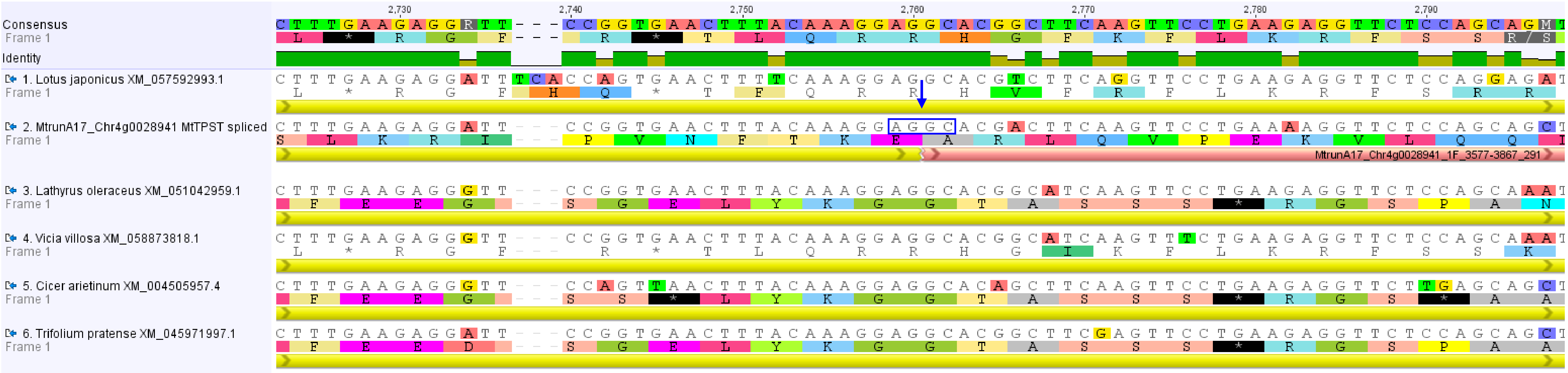


**Slide 5**. A fragment of the MUSCLE alignment of the *MtTPST* cDNA and five putative homologs from other legumes species after removing the putative intron from *MtTPST*. Yellow and pink bars indicate refORFs and ncORFs, respectively. The sequence AGGC (blue box and vertical arrow) corresponds to the putative exon-exon junction. After this simulated splicing, the *MtTPST* cDNA becomes highly similar to the cDNA sequences of other legumes species, with the splicing site 100% conserved in this alignment. This indicates that the annotated model of *MtTPST* is only a splicing variant, which may exist along with but not instead of the fully spliced form, the latter one being supported by MS data (two peptides, see Slides 1 and 2). Translation is shown relative to the refORFs of all six sequences.

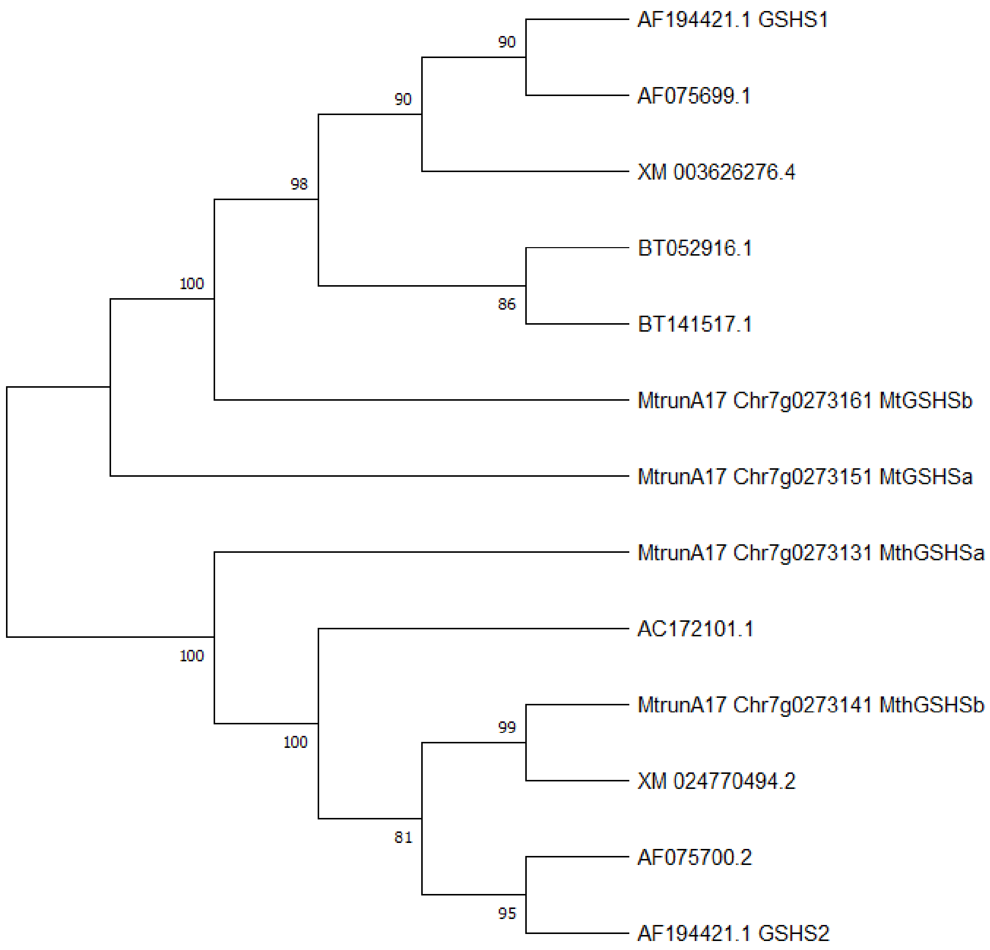


**Slide 6**. An unrooted UPGMA tree generated from a MUSCLE alignment of *MthGSHSb* and its 12 putative homologs/sequence variants from *M. truncatula*. *MthGSHSb* appears to be most related to XM_024770494.2, which has no corresponding annotated locus in the current version of the *M. truncatula* genome (v. 5.1.9). The tree was constructed using 1000 bootstrap replicates on cDNA or CDS sequences without trimming to preserve the information on 5'- and 3'-untranslated regions, where two putative ncORFs are located in *MthGSHSb*.

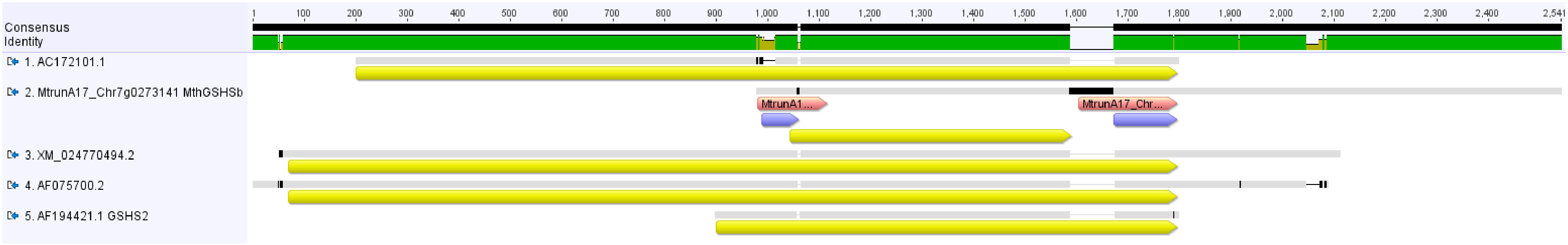


**Slide 7**. A MUSCLE alignment of the *MthGSHSb* cDNA and four sequences from the same clade (Slide 6). Yellow, pink, and violet arrow bars indicate refORFs, ncORFs, and the strongly conserved regions, respectively. The complete absence of neutral polymorphic sites within the refORFs of these sequences indicates their origin from a single genetic locus. Only the sequence of *MthGSHSb* has a corresponding annotated locus in the current version of the *M. truncatula* genome (v. 5.1.9). Although there is no gene model for a long form combining the upstream *MtGSHSa/MthGSHSa*-like refORF with the downstream *MtGSHSb/MthGSHSb*-like refORF (Slide 9) in a single cDNA in *M. truncatula*, such forms are ubiquitous in other eudicots. This suggests that the sequences shown here are either splicing variants of the same locus or artifacts of the genome assembly. It should be noted that two MS peptides, VIVNNESGYMVR and DKVIVNNESGYMVR, that we identified in six proteomic samples of three independent studies (PXD002692, Marx et al., 2016; PXD013606, Shin et al., 2021; PXD022278, Castañeda et al., 2021) are unique to the downstream ncORF12 (MtrunA17_Chr7g0273141_1F_625-816_192) and the refProts of AC172101.1, AF075700.2/XM_024770494.2, and AF194421.1 GSHS2. This means that ncORF12 is either translated as a discrete unit or as a C-terminal part of a refProt, which corresponds to one valid gene model missed in the current version of the *M. truncatula* genome.

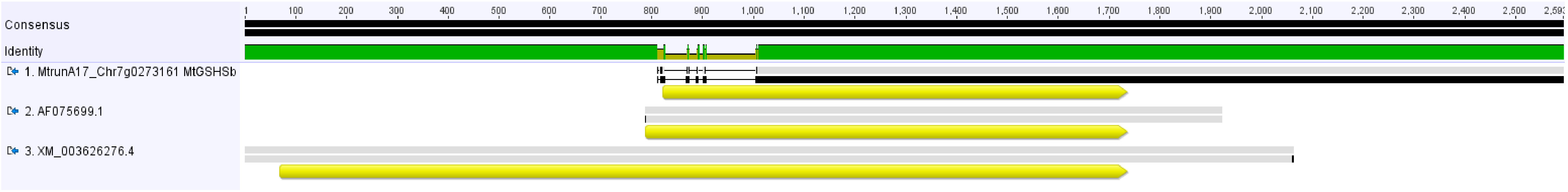


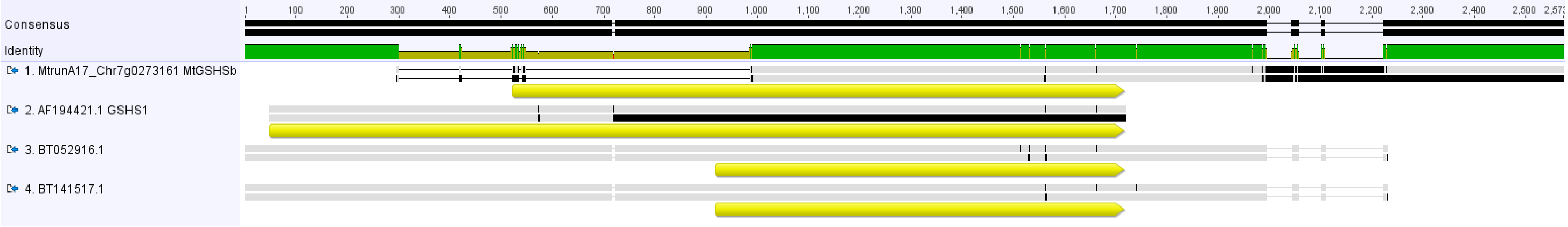


**Slide 8**. MUSCLE nucleotide alignments of the *MthGSHSb* homolog, *MtGSHSb*, with other sequences from its clade. Yellow arrow bars indicate refORFs. While sequences shown in the top alignment are likely to originate from a single genetic locus (no neutral polymorphisms in their refORFs), the bottom alignment contains neutral, missense, and frameshifting single-nucleotide differences, which indicates that the sequences can either have a heterogenous origin or may be artifacts of the genome assembly/sequencing errors. These sequences (except for *MtGSHSb*) have no corresponding annotated loci in the current version of the *M. truncatula* genome (v. 5.1.9). Validation of these putative ORFs with MS proteomics is necessary to ascertain their status.

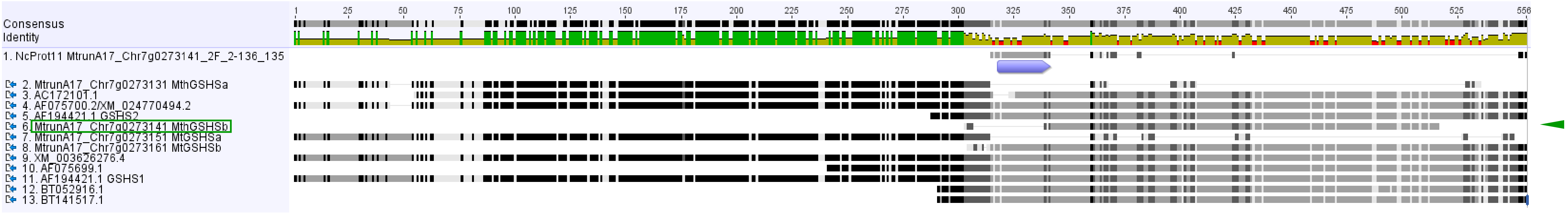


**Slide 9**. A MUSCLE protein alignment of MthGSHSb, its upstream CDS-overlapping putative ncProt11 (MtrunA17_Chr7g0273141_2F_2-136_135), and 11 putative homologs/proteoforms of MthGSHSb from *M. truncatula*. The strongly conserved region of the ncProt aligns with refProts of other cDNA sequences (shown as a violet arrow bar). This region is absent from the refProt of *MthGSHSb*. The latter aligns with the central or C-terminal portions of longer sequences. It is the shortest refProt in the alignment (green box and arrowhead).

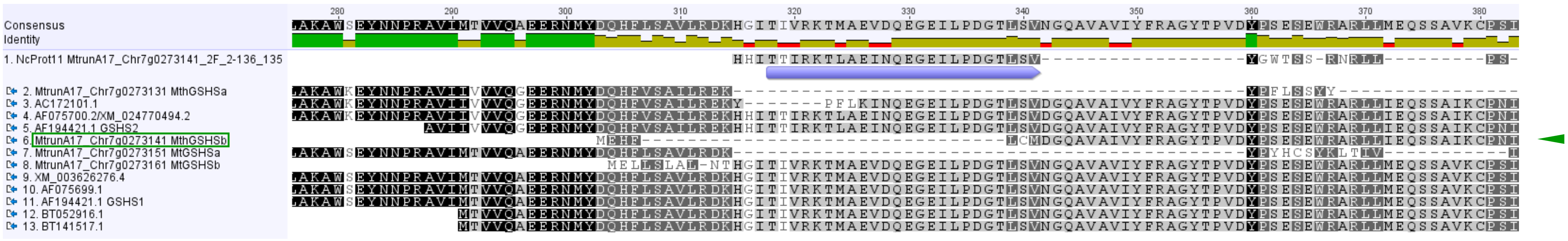


**Slide 10**. Details of the protein alignment shown in the previous slide. The refProt of *MthGSHSb* starts with the sequence MEHFLC, which is found neither in the ncProt nor in other refProts. This is because the reading frame of *MthGSHSb* in this region is shifted forwardly by one nucleotide relative to the ncProt and refProts. The MEHFLCM translation in the refProt of *MthGSHSb* corresponds to the region translated to DGTLSV in refProts of other sequences (see the next slide). Thus, the partial alignment of MEHF with DQHF is purely artifactual. The return of the reading frame of *MthGSHSb* to the same frame with other refProts is achieved by an addition of a four-nucleotide sequence TATG at the end of the first exon, which is translated to CM. Most likely, TATG comes from the retention of the first four nucleotides of Intron 1 (genomic DNA in Slide 12). It remains to be shown if a sequence variant without this retained intron is translated.

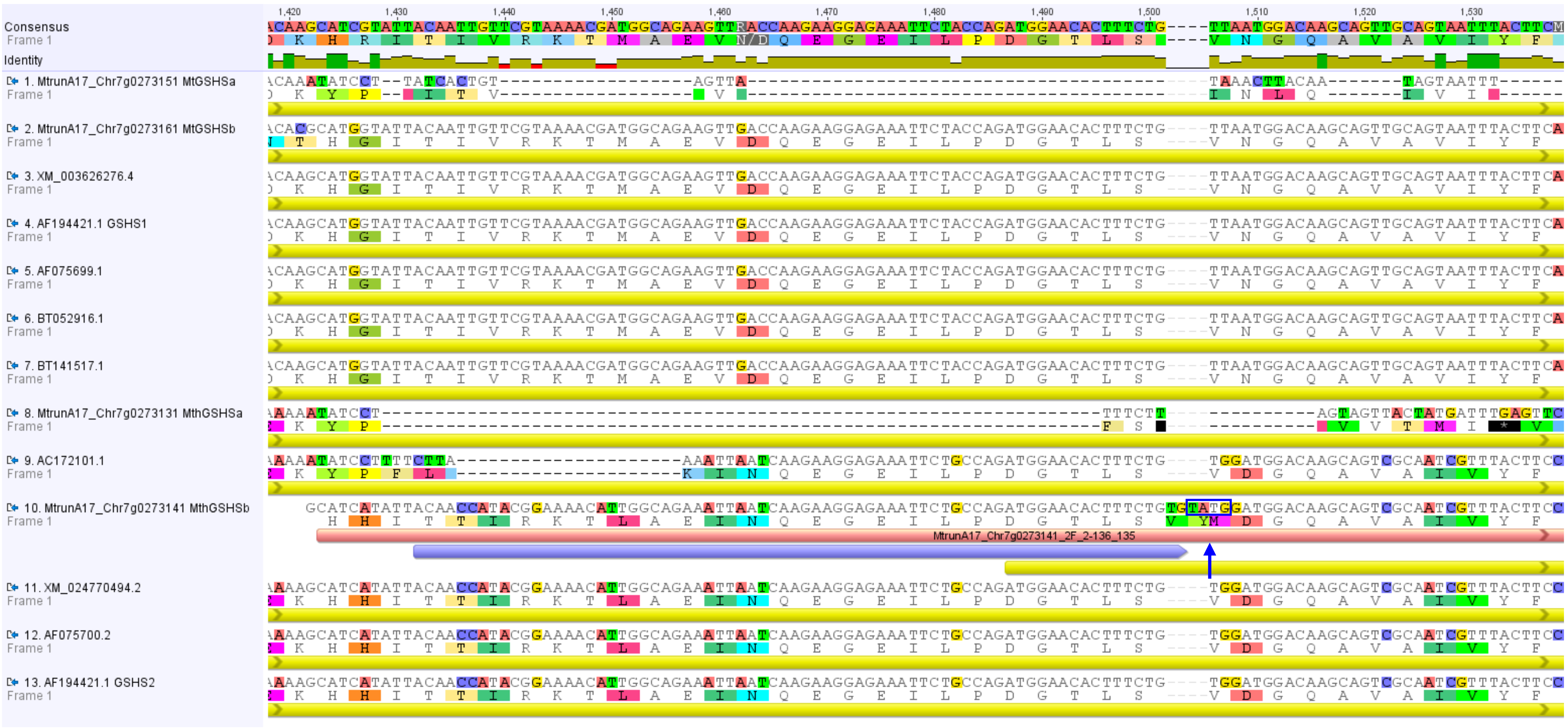


**Slide 11**. A fragment of a MUSCLE alignment of the *MthGSHSb* cDNA and 12 putative homologs/sequence variants of *MthGSHSb* from *M. truncatula*. This alignment is the basis for the tree in Slide 6. NcORF11 (MtrunA17_Chr7g0273141_2F_2-136_135) is shown as a pink bar. The strongly conserved region of the ncProt (violet arrow bar) aligns with refProts and is highly similar to them at the amino acid level. Translation is shown relative to the consensus, which is the frame of most refProts (yellow bars), but not the frame in which the refProt of *MthGSHSb* starts. The return to the same frame with other refProts was achieved by an addition of a four-nucleotide sequence TATG shown as blue box with a vertical arrow (see the next slide).

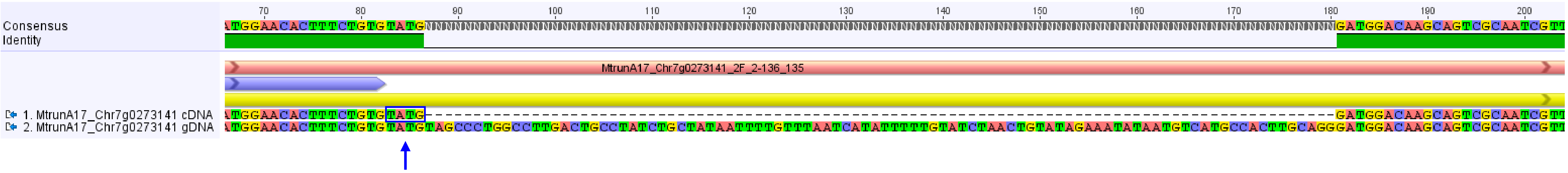


**Slide 12**. A fragment of a MUSCLE alignment of the cDNA (top) and the gDNA (bottom) of *MthGSHSb*. The refORF, ncORF11, and the strongly conserved region are shown as yellow, pink, and violet bars/arrow bars, respectively. The blue box indicates the last four nucleotides of Exon 1, which were presumably retained from Intron 1 to produce the annotated version of *MthGSHSb*. This splicing event could be unique to *MthGSHSb*. It switched translation from the alternative frame (MEHFLCM) back to the frame of most other refProts (DGQAVAI…), as can be seen in Slide 10. It remains to be shown if a sequence variant without this retained intron is translated.

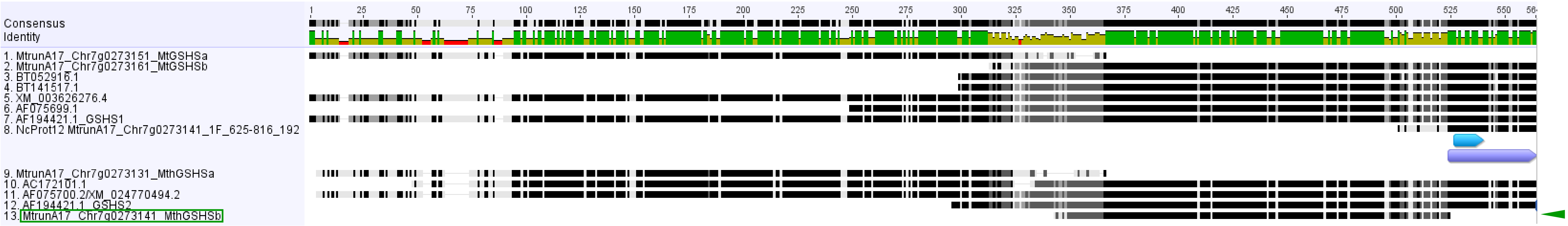


**Slide 13**. A Clustal Omega protein alignment of MthGSHSb, its downstream putative ncProt12 (MtrunA17_Chr7g0273141_1F_625-816_192), and 11 putative homologs/proteoforms of MthGSHSb from *M. truncatula*. The strongly conserved region of the ncProt, shown as a violet arrow bar, aligns with refProts of other cDNA sequences. This region is absent from the refProt of *MthGSHSb* (green box and arrowhead). The region matching the MS peptide DKVIVNNESGYMVR is shown in blue. It is unique to the ncProt and sequences 10, 11, and 12 in the alignment.

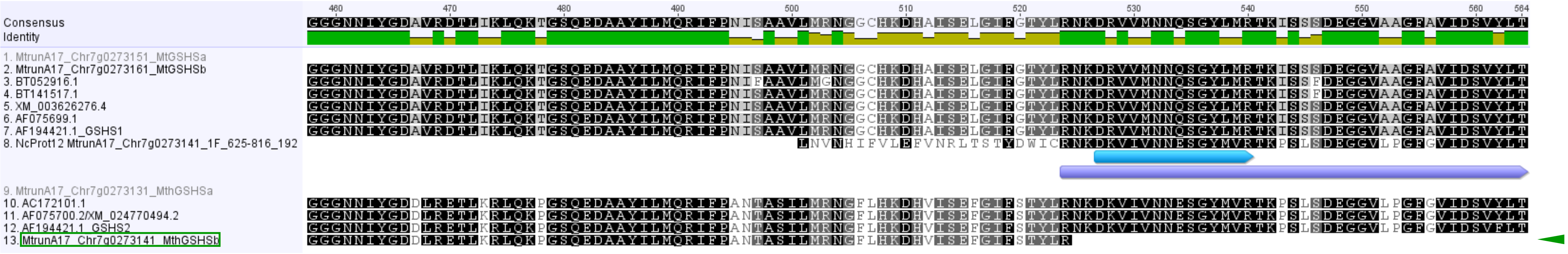


| | Description | Scientific Name | Max Score | Total Score | Query Cover | E value | Per. Ident | Acc. Len | Accession |
|---|---|---|---|---|---|---|---|---|---|
| | Medicago truncatula homoglutathione synthetase GSHS2 (gshs2) gene, partial cds; and glutathione syntheta… | Medicago trunca… | 158 | 158 | 100% | 5e-34 | 100.00% | 9597 | AF194421.1 |
| | Medicago truncatula chromosome 7 clone mth2-29f3, complete sequence | Medicago trunca… | 158 | 158 | 100% | 5e-34 | 100.00% | 88313 | AC172101.1 |
| | Medicago lupulina genome assembly, chromosome: 2 | Medicago lupulina | 130 | 130 | 100% | 1e-25 | 94.19% | 73454271 | OY283140.1 |
| | Medicago arabica genome assembly, chromosome: 6 | Medicago arabica | 124 | 124 | 100% | 5e-24 | 93.02% | 61901166 | OX326969.1 |
| | Trifolium dubium genome assembly, chromosome: 11 | Trifolium dubium | 69.4 | 69.4 | 79% | 2e-07 | 85.29% | 38738077 | OX638072.1 |
| | Trifolium dubium genome assembly, chromosome: 4 | Trifolium dubium | 62.1 | 62.1 | 74% | 4e-05 | 84.38% | 51571781 | OX638065.1 |

**Slide 14**. Top panel: Details of the protein alignment shown in the previous slide. The strongly conserved portion of ncProt12 corresponds to the refProts of other cDNA sequences. The ncProt starts with the sequence LNVNHIFVLEFVNRLTSTYDWIC, which has very weak similarity to bacterial proteins annotated as ABC transporter substrate-binding proteins (e-value of the top hit: 2.7, as of 25 February 2026). This region corresponds to an insertion of 85 nucleotides, which truncates the refProt of *MthGSHSb* and frameshifts the downstream portion of the ancestral refORF. Thus, ncORF11 is the remnant of the original longer refORF. Bottom panel: The sequence of 85 nucleotides has only a few hits in the BLASTN search using the NCBI non-redundant nucleotide database (as of 25 February 2026). It is likely to act as a retained intron in the current gene model (see the next slide).

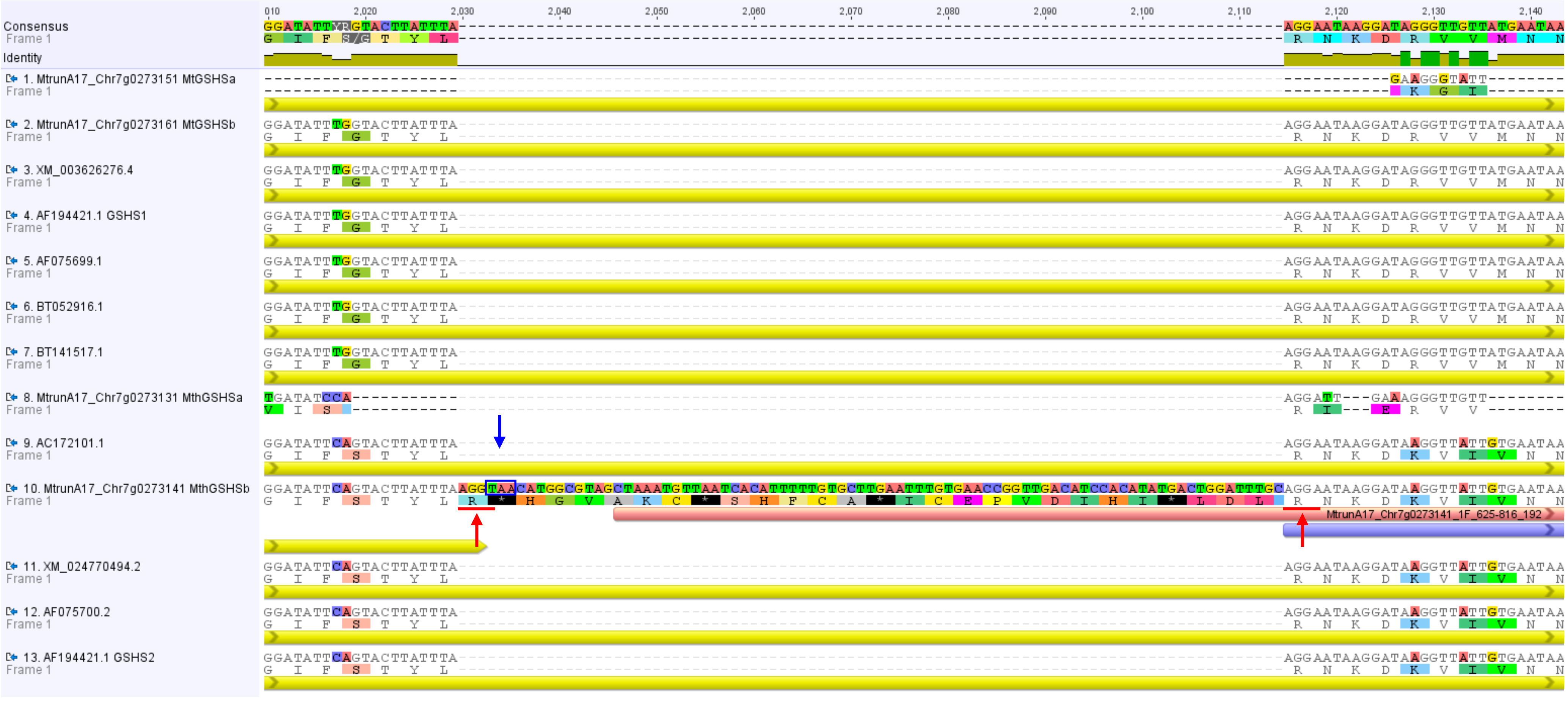


**Slide 15**. A fragment of the MUSCLE alignment from Slide 11 focusing on ncORF12. It aligns the *MthGSHSb* cDNA with 12 putative homologs/sequence variants of *MthGSHSb* from *M. truncatula*. Yellow, pink, and violet bars indicate refORFs, the ncORF, and the strongly conserved region, respectively. Translation is shown relative to the consensus. The sequence of 85 inserted nucleotides has an in-frame stop codon (blue box and vertical arrow) that truncates the refORF of *MthGSHSb*. The insertion frameshifts the downstream portion of the ancestral refORF. It is conceivable that the insertion may act as a retained intron, as follows from the presence of two sites very similar to the classical exon-intron and intron-exon borders (red-underlined AGGT and AGGA, respectively). When these 85 nucleotides are removed exactly at the putative splicing sites (red vertical arrows), the cDNA of *MthGSHSb* perfectly aligns with the other sequences listed here.

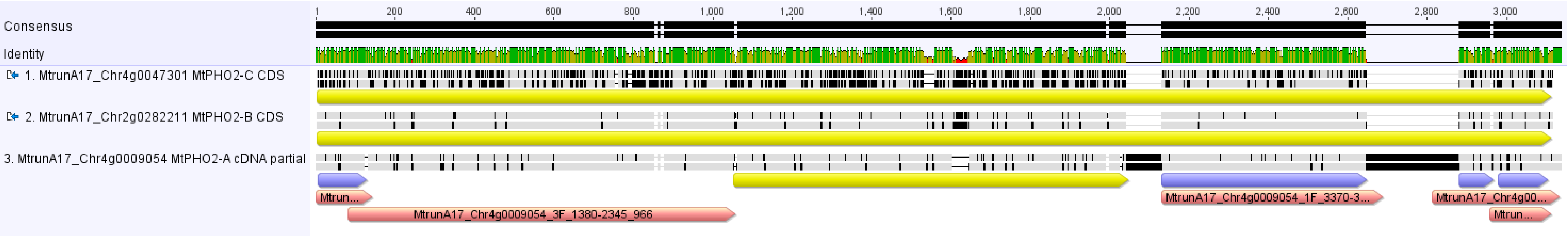


**Slide 16**. A MUSCLE alignment of the *MtPHO2-A* cDNA (with UTRs truncated) and the CDSs of two homologs. The refORFs, the ncORFs, and the strongly conserved regions are shown as yellow, pink, and violet arrow bars, respectively. The short refORF of *MtPHO2-A* and four strongly conserved regions in ncORFs 51, 53, 54, and 55 correspond to modified portions of the ancestral, longer refORF found in *MtPHO2-B* and *MtPHO2-C*. These five regions largely preserved the original amino acid sequence of the ancestral refProt. In contrast, ncORF52 (MtrunA17_Chr4g0009054_3F_1380-2345_966) is out of frame relative to the long refProts of *MtPHO2-B* and *MtPHO2-C*. This analysis indicates that *MtPHO2-A* can be categorized as a functional unitary pseudogene according to the definition of Cheetham et al. (2020).

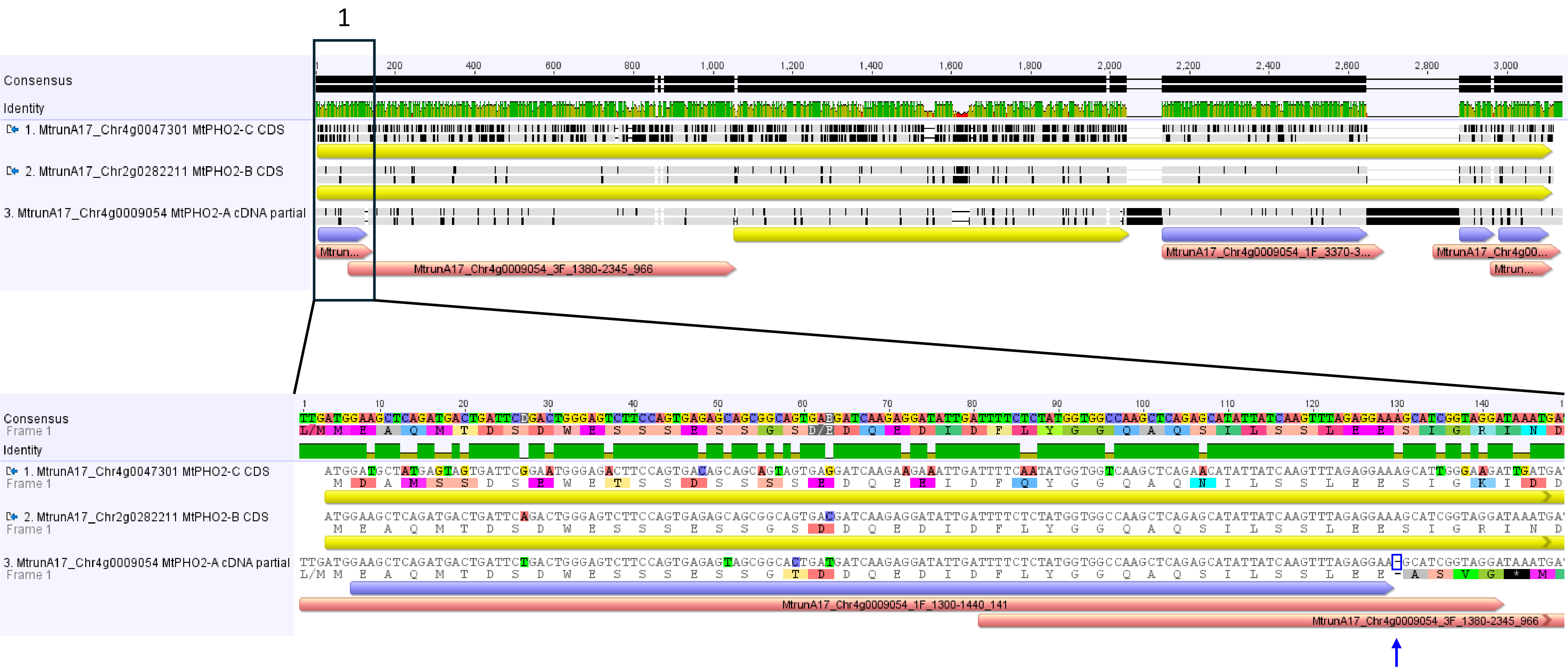


**Slide 17**. The first fragment of the alignment shown in the previous slide. The conserved region of ncProt51 (MtrunA17_Chr4g0009054_1F_1300-1440_141) is nearly identical to the beginning of MtPHO2-B. The original frame is shifted via the loss of a single nucleotide (presumably A; blue box and vertical arrow).

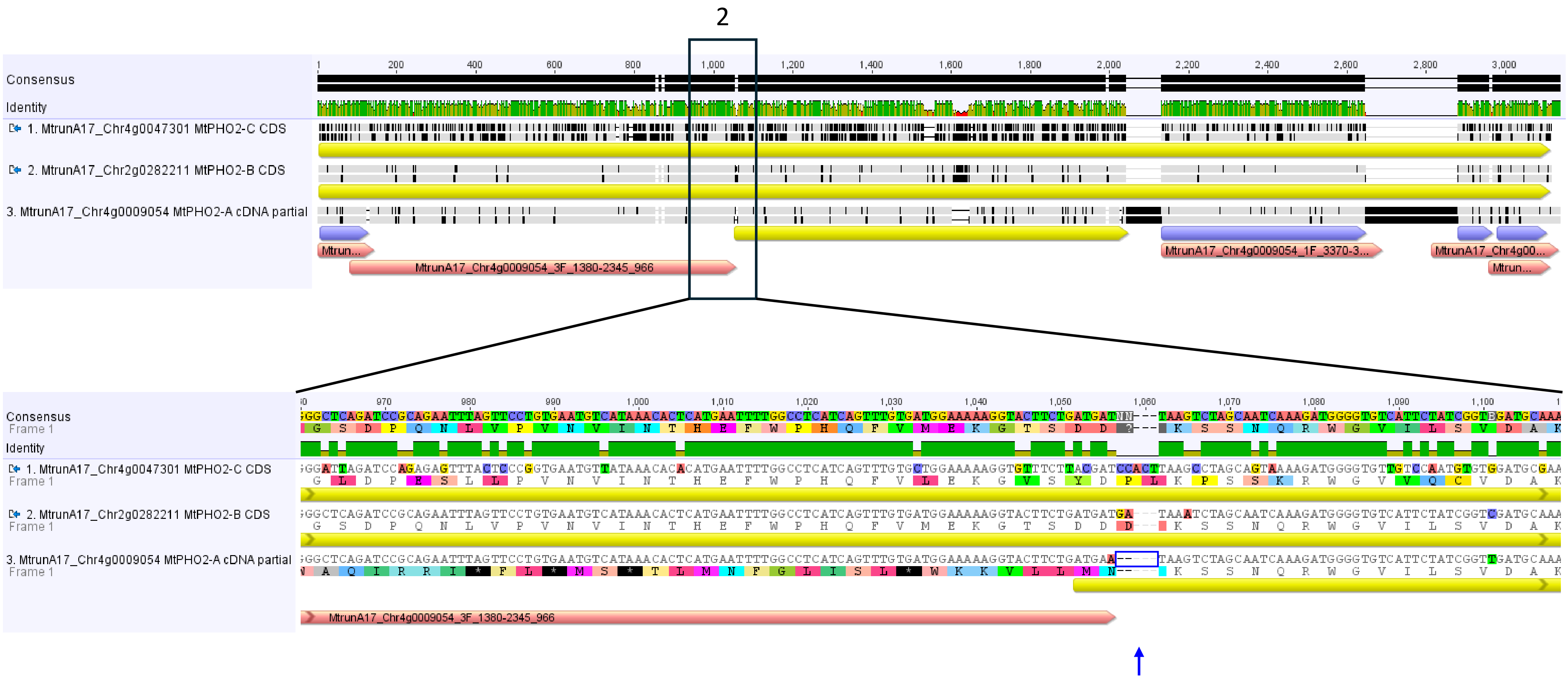


**Slide 18**. The second fragment of the same alignment. The original reading frame was regained by the modern version of *MtPHO2-A* via the deletion of multiple nucleotides at the end of ncORF52 (MtrunA17_Chr4g0009054_3F_1380-2345_966).

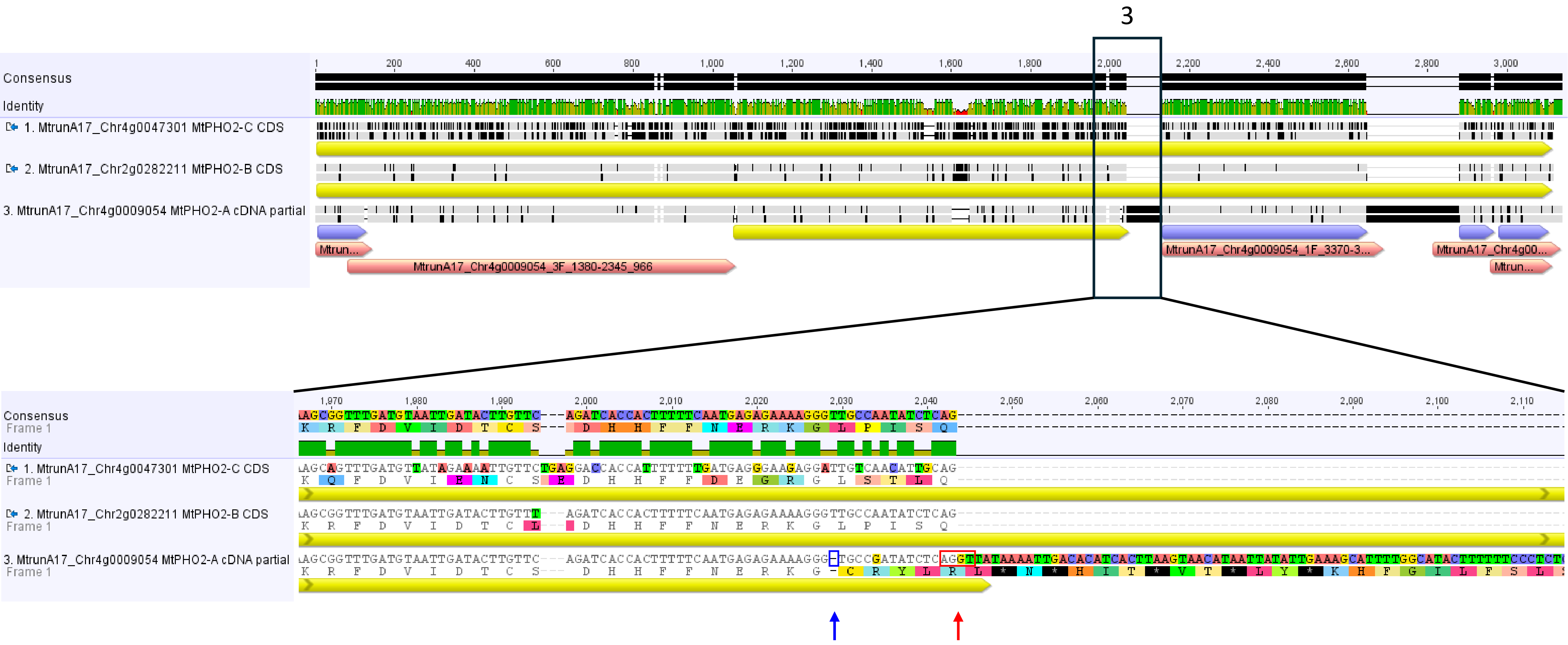


**Slide 19**. The third fragment of the same alignment. The refORF of *MtPHO2-A* was prematurely terminated by a single-nucleotide deletion (presumably T, blue box and vertical arrow), which caused a frameshift and introduced an early in-frame stop codon. However, the downstream cDNA sequence until the beginning of ncORF53 (MtrunA17_Chr4g0009054_1F_3370-3927_558) is not present in the cDNA of *MtPHO2-B* and *MtPHO2-C.* Most likely, it originates not from a recent insertion but from an intron retention event because it is found in the genomic DNA of all three genes (see the next slide). This possibility is further supported by the presence of a sequence AGGT precisely at the end of the aligned region (red box and vertical arrow), which is very similar to the classical exon-intron border motif.

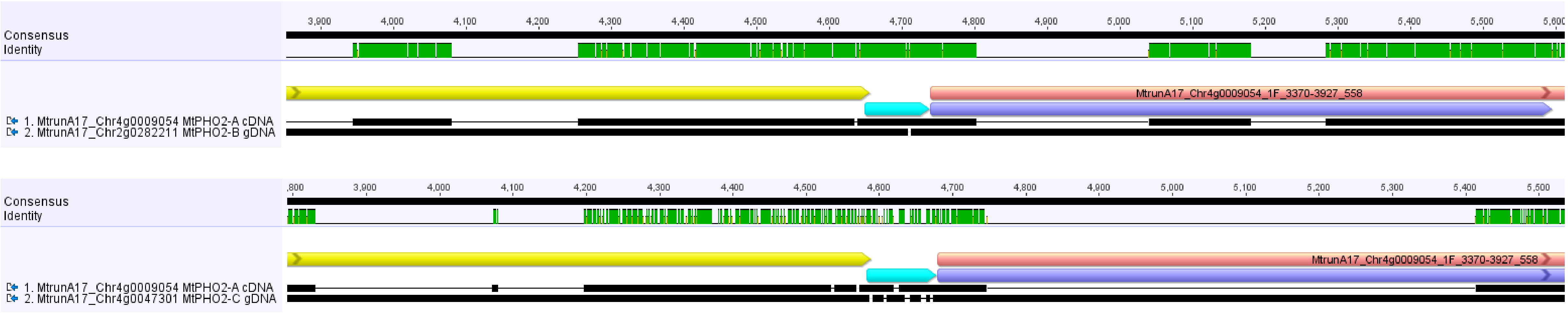


**Slide 20**. Fragments of two MUSCLE alignments of the *MtPHO2-A* cDNA and the genomic DNA of its two homologs. The region shown as a cyan arrow bar between the refORF and ncORF53 (MtrunA17_Chr4g0009054_1F_3370-3927_558) is present in the genomic DNA of these genes. Thus, it is an intron retained by *MtPHO2-A* but spliced out in the other two genes.

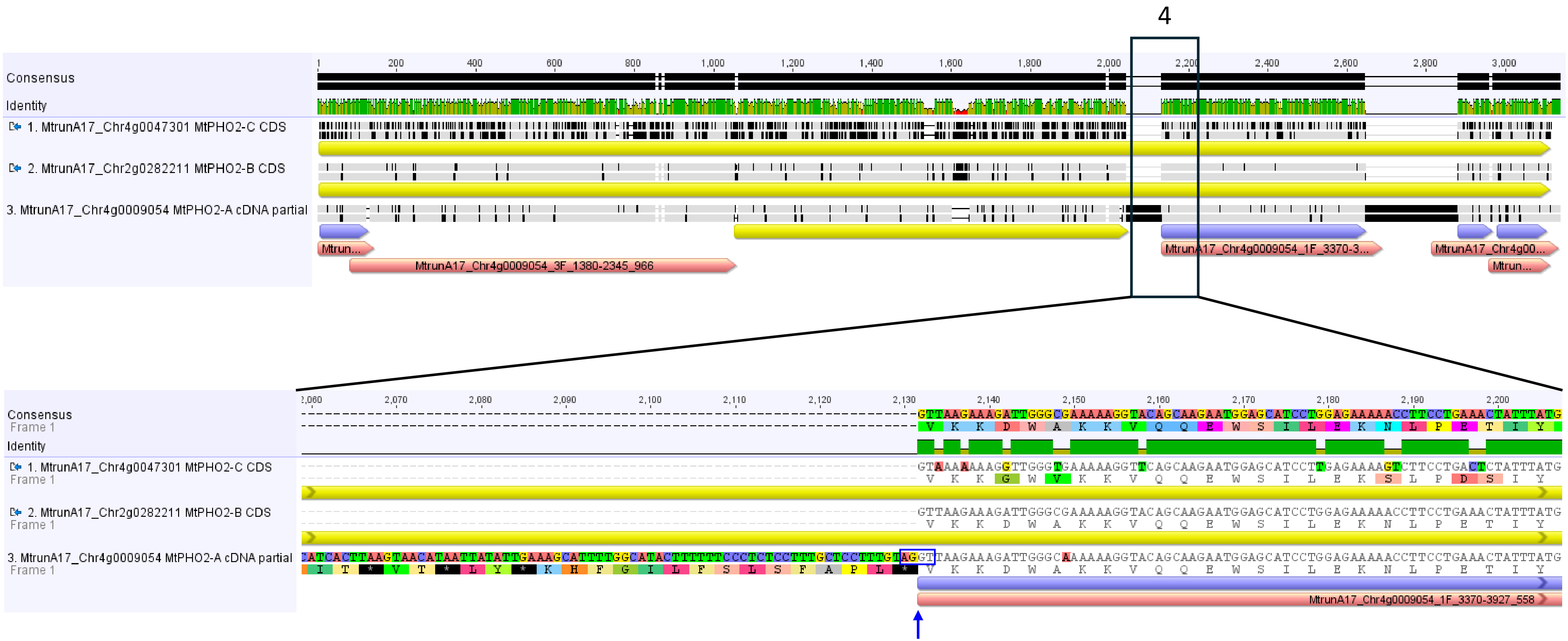


**Slide 21**. The fourth fragment of the alignment shown earlier. The intron retention event terminated the refORF of *MtPHO2-A* but brought the downstream ncORF53 (MtrunA17_Chr4g0009054_1F_3370-3927_558) back to the same frame with refORFs of the other genes. Note the presence of a sequence AGGT precisely at the end of the putative retained intron (blue box and vertical arrow), which is very similar to the classical intron-exon border motif.

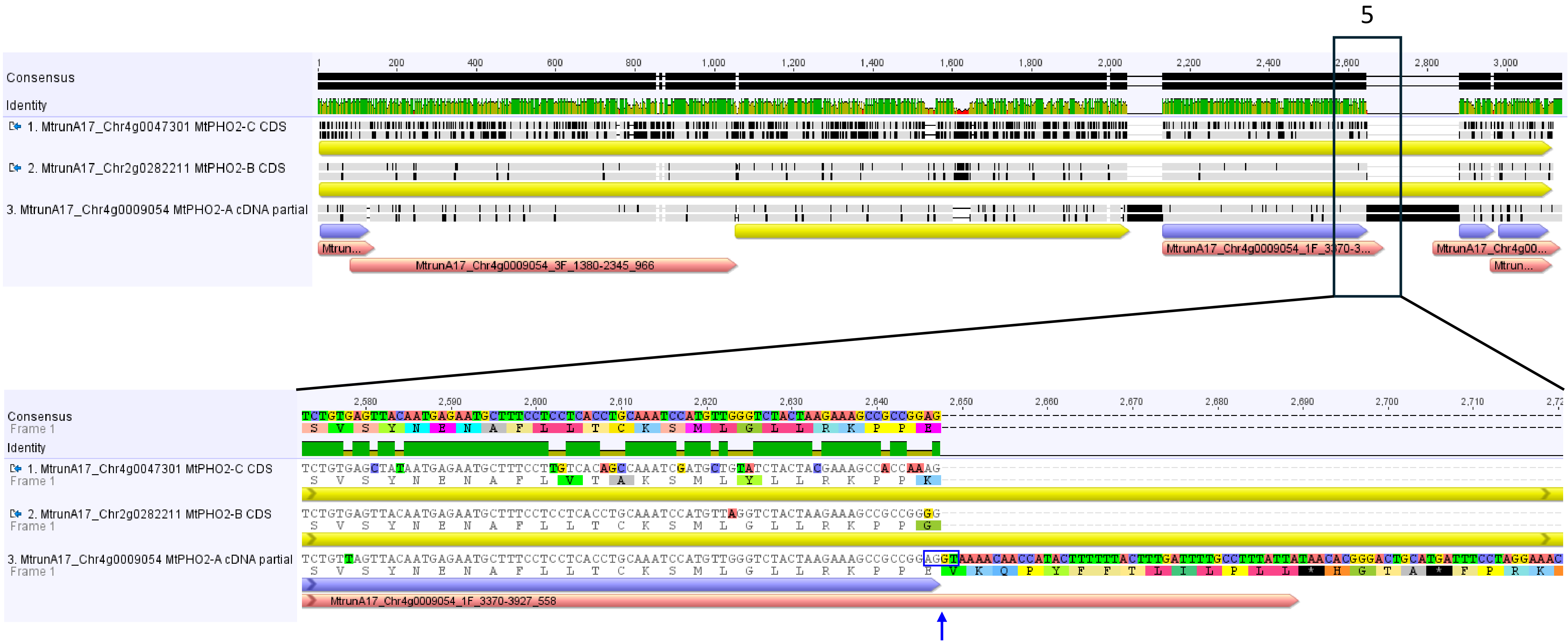


**Slide 22**. The fifth fragment of the same alignment. The sequence downstream of the conserved region of ncORF53 (MtrunA17_Chr4g0009054_1F_3370-3927_558) is not present in the cDNA of *MtPHO2-B* and *MtPHO2-C*. Most likely, it originates from an incomplete intron retention event because it is partially found in the genomic DNA of *MtPHO2-A* and *MtPHO2-B* (see the next slide). This possibility is further supported by the presence of a sequence AGGT precisely at the end of the aligned region (blue box and vertical arrow), which is very similar to the classical exon-intron border motif.

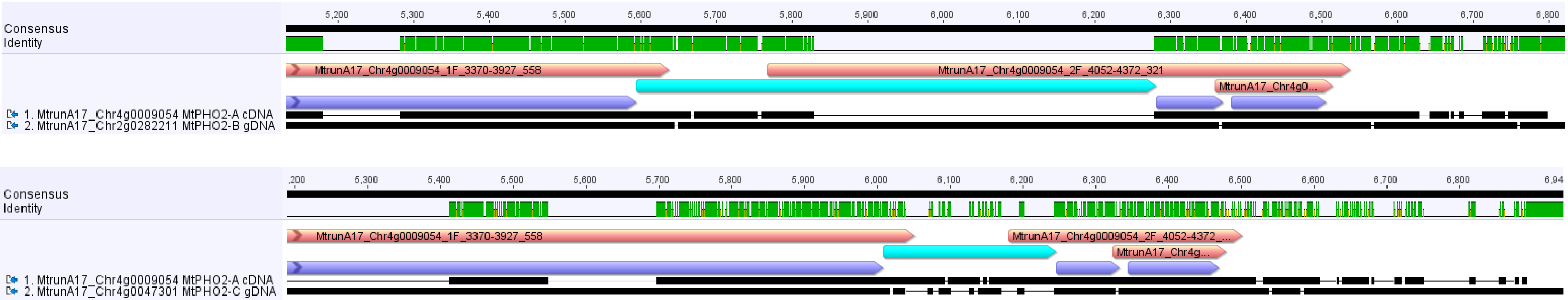


**Slide 23**. Fragments of two MUSCLE alignments of the *MtPHO2-A* cDNA and the genomic DNA of its two homologs (the same alignments as used in Slide 20). The region shown as a cyan arrow bar between conserved regions of ncORF53 (MtrunA17_Chr4g0009054_1F_3370-3927_558) and ncORF54 (MtrunA17_Chr4g0009054_2F_4052-4372_321) is present partially in the genomic DNA of *MtPHO2-B*. It is lost or heavily modified in *MtPHO2-C*. Thus, it is an intron incompletely retained by *MtPHO2-A* but spliced out by *MtPHO2-B*.

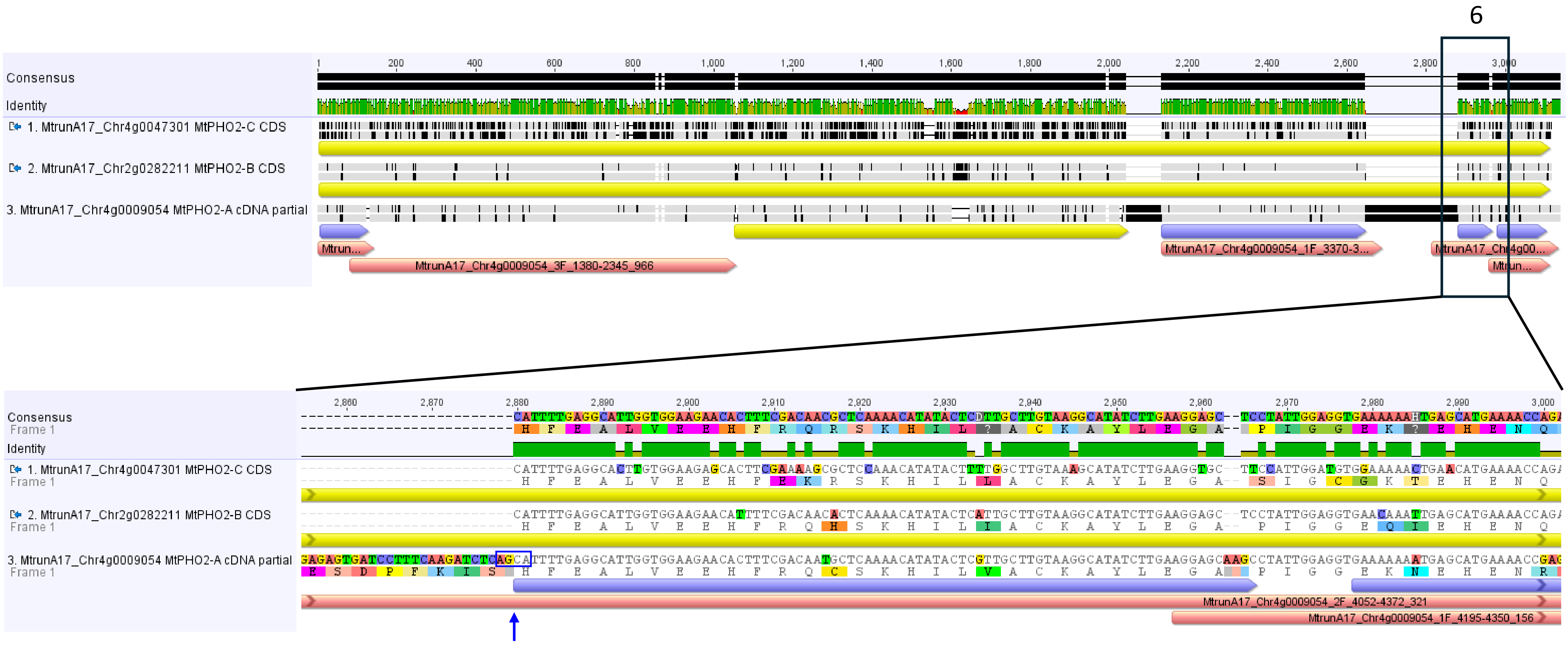


**Slide 24**. The sixth fragment of the same alignment. Because of the incomplete retention of an ancestral intron, the conserved region of ncORF54 (MtrunA17_Chr4g0009054_2F_4052-4372_321) is out of frame with the refProts of *MtPHO2-B* and *MtPHO2-C*. However, it is nearly identical to the refProts at the amino acid level. Note the presence of a sequence AGCA precisely at the end of the putative retained intron (blue box and vertical arrow), which is somewhat similar to the classical intron-exon border motif.

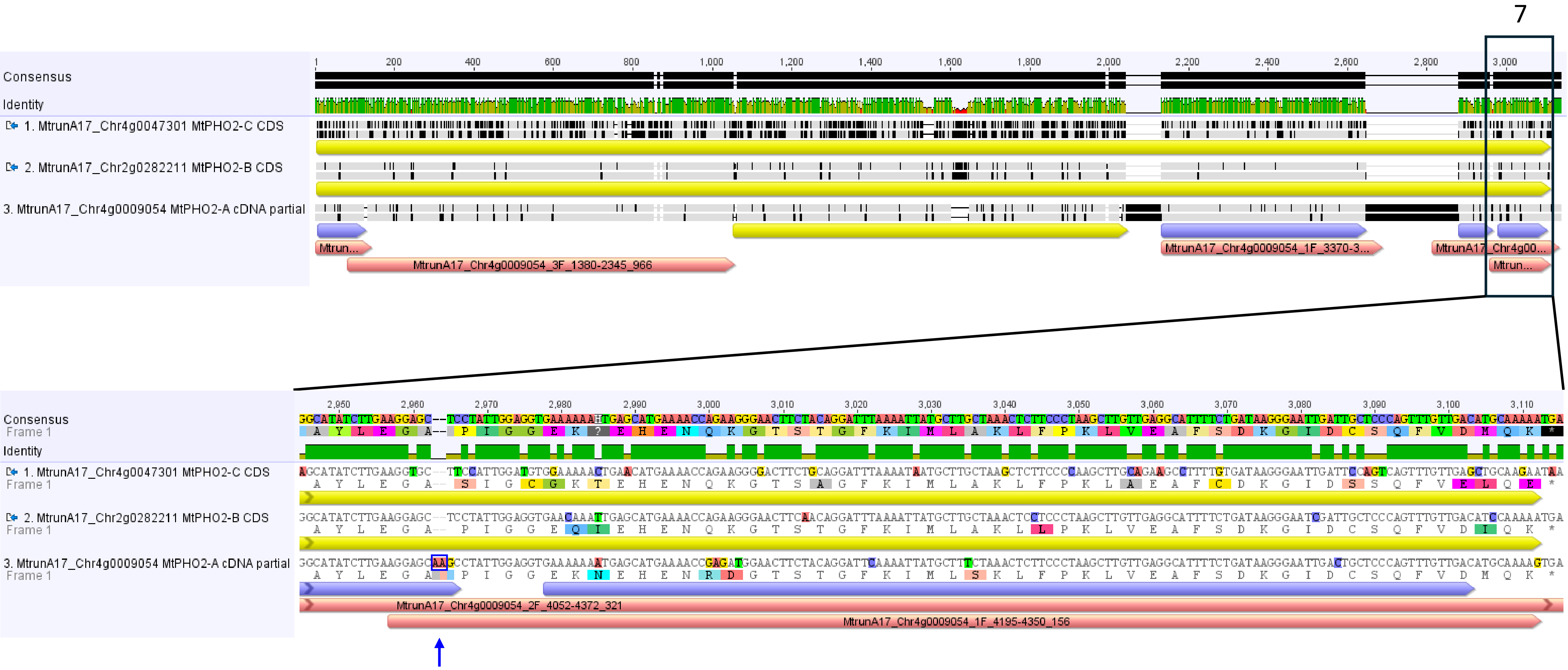


**Slide 25**. The seventh fragment of the same alignment. Because of a two-nucleotide insertion indicated with a blue box and a vertical arrow, the strongly conserved region of the last ncORF in this gene (atlORF55, MtrunA17_Chr4g0009054_1F_4195-4350_156) regains the same reading frame with refProts of *MtPHO2-B* and *MtPHO2-C*. Thus, the ncORF terminates exactly at the same position with the refORFs.

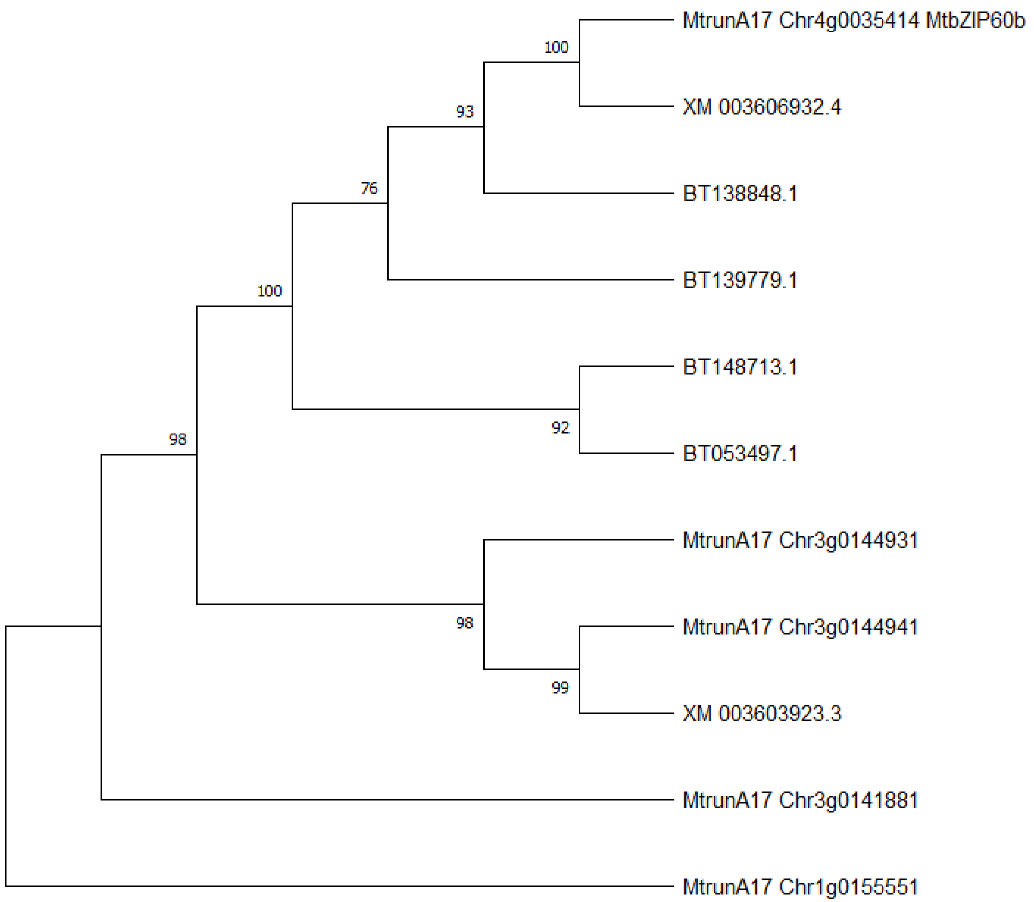


**Slide 26**. An unrooted UPGMA tree generated from a MUSCLE alignment of *MtbZIP60b* and its 10 putative homologs/sequence variants from *M. truncatula*. *MtbZIP60b* appears to be most related to XM_003606932.4. However, the remaining four sequences in the same clade also share very high percent nucleotide identity with *MtbZIP60b* (see the next slide). The tree was constructed using 1000 bootstrap replicates on cDNA or CDS sequences without trimming to preserve the information on the 5'-untranslated region, where the putative ncORF60 is located in *MtbZIP60b*.

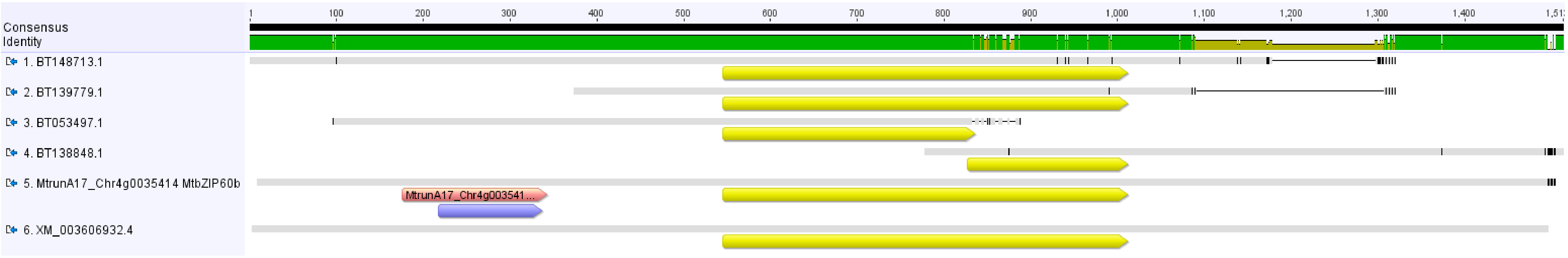


**Slide 27**. A MUSCLE alignment of the *MtbZIP60b* cDNA and five sequences from the same clade (Slide 26). Yellow, pink, and violet arrow bars indicate refORFs, the ncORF, and the strongly conserved region, respectively. The nearly complete absence of neutral polymorphic sites in the first half of the alignment indicates the origin of all six sequences from a single genetic locus. Only the sequence of *MtbZIP60b* has a corresponding annotated locus in the current version of the *M. truncatula* genome (v. 5.1.9).

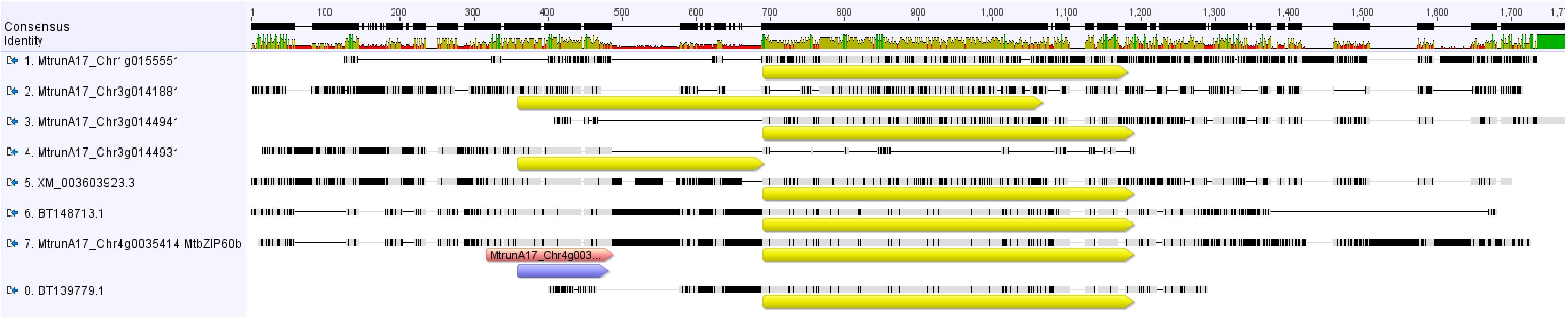


**Slide 28**. A MUSCLE alignment of the *MtbZIP60b* cDNA and seven selected putative homologs/sequence variants of *MtbZIP60b* from *M. truncatula*. NcORF60 (MtrunA17_Chr4g0035414_3F_168-335_168) is shown as a pink arrow bar. Despite the low percent identity among the sequences, the strongly conserved region of the ncProt (violet arrow bar) aligns with refProts of MtrunA17_Chr3g0141881 and MtrunA17_Chr3g0144931. Although sequences MtrunA17_Chr3g0144931 and MtrunA17_Chr3g0144941 are currently annotated as two different loci (v. 5.1.9), they probably correspond to two portions of a single genetic unit similar to MtrunA17_Chr3g0141881. This possibility is supported by their adjacent location in the genome (see the next slide).

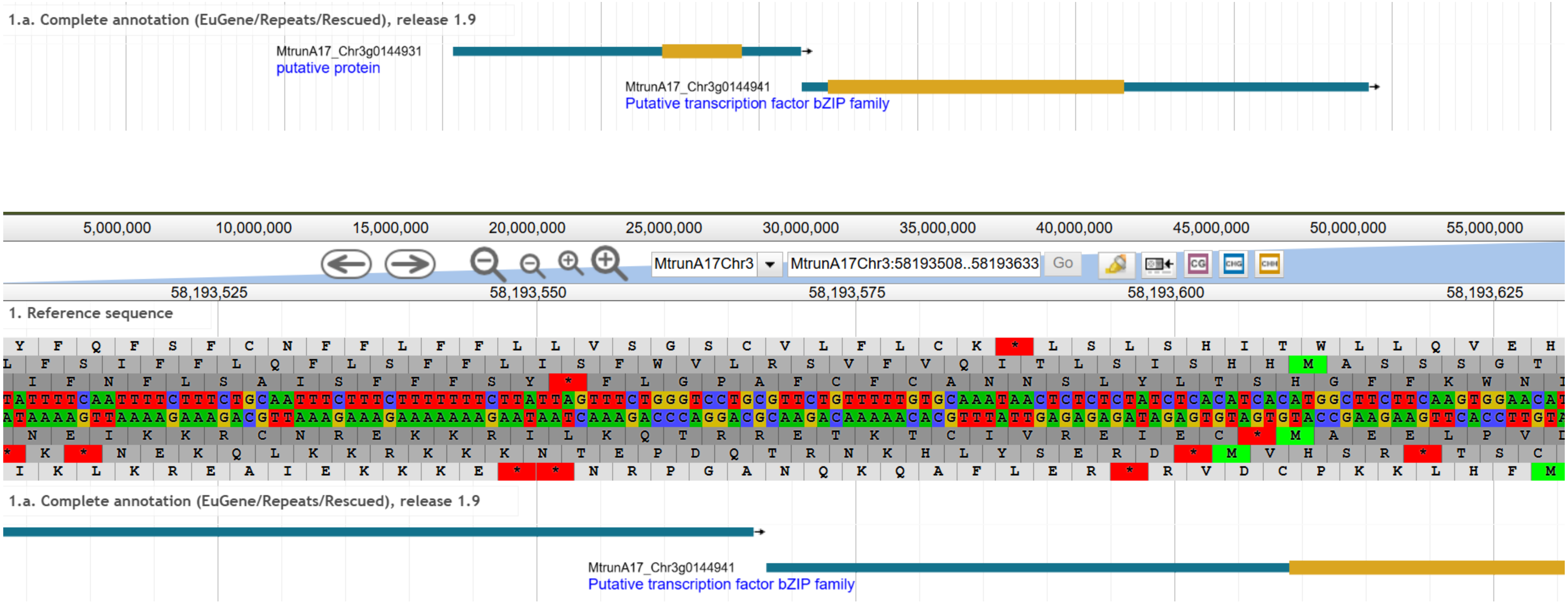


**Slide 29**. Two screenshots from the *M. truncatula* Genome Browser v. 5.1.9 illustrating the adjacent location of loci MtrunA17_Chr3g0144931 and MtrunA17_Chr3g0144941 in the genome. The loci are separated by a single base (C in the top strand). Thus, they are likely to correspond to two portions of a single genetic unit similar to MtrunA17_Chr3g0141881 (see the previous slide).

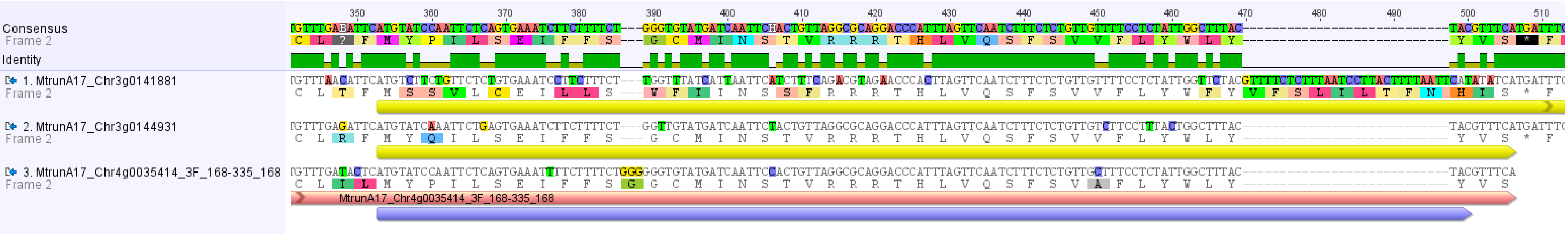


**Slide 30**. A MUSCLE nucleotide alignment of ncORF60 (MtrunA17_Chr4g0035414_3F_168-335_168) and two highly similar sequences from *M. truncatula*. The strongly conserved region of the ncProt (violet arrow bar) aligns with refProts (yellow arrow bars) of MtrunA17_Chr3g0141881 and MtrunA17_Chr3g0144931 and is highly similar to them at the amino acid level.

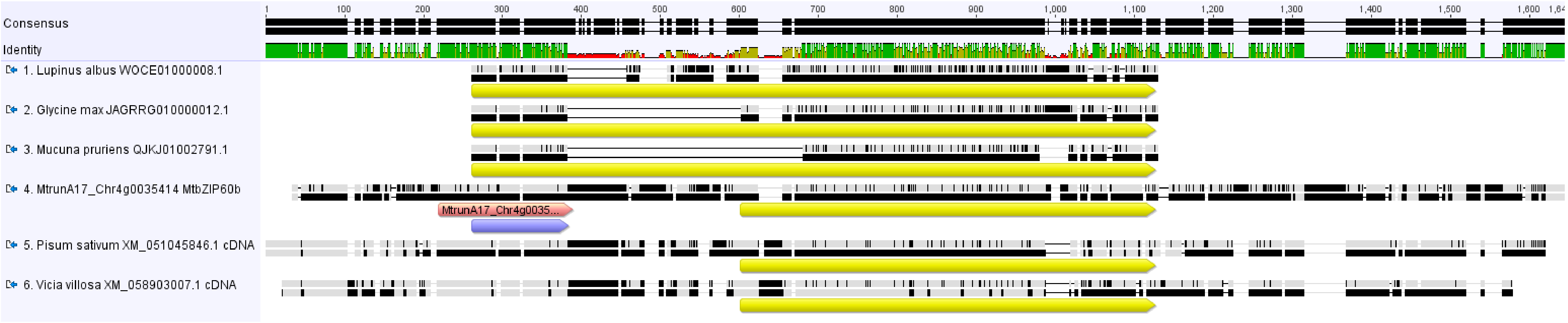


**Slide 31**. A MUSCLE alignment of the *MtbZIP60b* cDNA and CDS/cDNA sequences of homologs from five other legume species. The refORFs, the ncORF, and the strongly conserved region are shown as yellow, pink, and violet arrow bars, respectively. A sequence separating the strongly conserved region and the refORF of *MtbZIP60b* is not found in any annotated homolog in *M. truncatula* (Slide 28). However, it is present and partially conserved in two other legume species (sequences 5 and 6). This sequence is unlikely to correspond to a retained intron because it is missing from the genomic DNA of the long homolog MtrunA17_Chr3g0141881 (Slide 33). The separating sequence is shared by legumes from five genera, specifically *Medicago*, *Pisum*, *Vicia*, *Trifolium*, and *Lathyrus* (Slides 33 and 34).

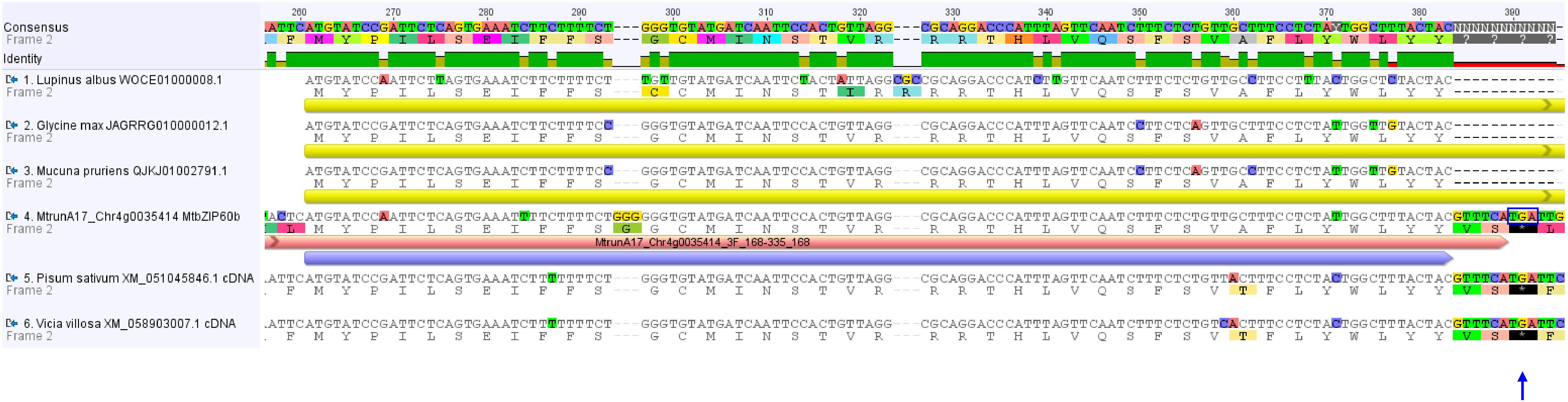


**Slide 32**. A fragment of the MUSCLE alignment shown in Slide 31. The very high conservation of the region depicted as a violet arrow bar in at least six different legume species suggests that it has been in use for translation for a long time. This conservation can be explained by a hypothetical scenario in which the ncORF acts as an exon under certain circumstances and is essential in sequences like *MtbZIP60b*. Its continuous translation without a special splicing event is prevented by an in-frame stop codon (blue box and vertical arrow) introduced by the separator sequence. Alternatively, the ncORF is translated and essential without being part of a longer sequence.

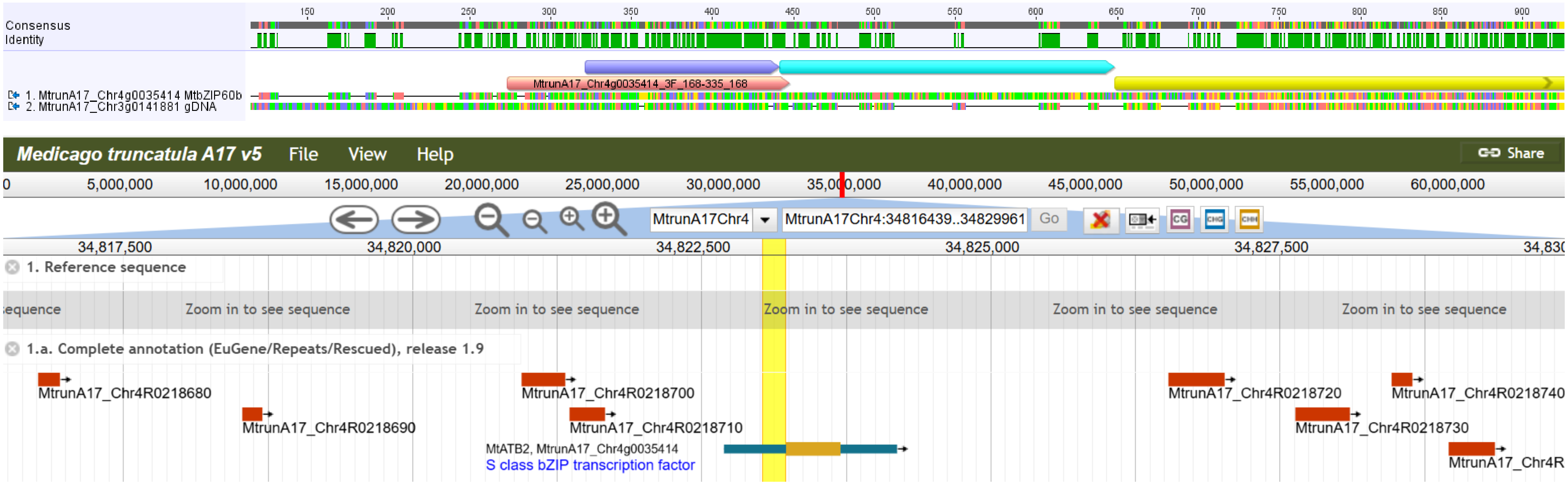


**Slide 33**. Top panel: A MUSCLE alignment of the *MtbZIP60b* cDNA and the genomic DNA of the long homolog, MtrunA17_Chr3g0141881. The sequence shown as a cyan arrow bar separates the strongly conserved region (violet arrow bar) from the refORF of *MtbZIP60b* (yellow arrow bar). It has no equivalent in the genomic DNA of MtrunA17_Chr3g0141881. This suggests that the ncORF evolved through an event other than the failure to splice out an intron. Bottom panel: The region highlighted in yellow corresponds to the separator sequence. It does not coincide with any transposon. We hypothesize that it corresponds to an insertion of an unknown origin (not a transposable element, see also the next slide). It is shared by legumes from five genera, specifically *Medicago, Pisum, Vicia, Trifolium, and Lathyrus* (see the next slide) and is lost or heavily modified in other *M. truncatula* homologs of *MtbZIP60b*. More likely, the common ancestor of these legume species already had two types of such sequences: at least one with the separator and at least one without. The separator effectively splits an ancient refORF into two parts, both of which retained the protein-coding potential. However, only the downstream one is annotated as a protein-coding sequence.

| | Description | Scientific Name | Max Score | Total Score | Query Cover | E value | Per. Ident | Acc. Len | Accession |
|---|---|---|---|---|---|---|---|---|---|
| ☑ | PREDICTED: Medicago truncatula bZIP transcription factor 11 (LOC11445368), mRNA | Medicago truncatula | 383 | 383 | 100% | 2e-101 | 100.00% | 1493 | XM_003606932.4 |
| ☑ | Medicago truncatula clone JCVI-FLMt-16A14 unknown mRNA | Medicago truncatula | 383 | 383 | 100% | 2e-101 | 100.00% | 1201 | BT148713.1 |
| ☑ | Medicago truncatula clone MTYFP_FQ_FR_FS1G-K-6 unknown mRNA | Medicago truncatula | 383 | 383 | 100% | 2e-101 | 100.00% | 776 | BT053497.1 |
| ☑ | Medicago truncatula clone mth2-31b9, complete sequence | Medicago truncatula | 383 | 383 | 100% | 2e-101 | 100.00% | 146189 | AC121244.13 |
| ☑ | Medicago truncatula clone JCVI-FLMt-8J24 unknown mRNA | Medicago truncatula | 316 | 316 | 83% | 2e-81 | 100.00% | 730 | BT139779.1 |
| ☑ | Medicago arabica genome assembly, chromosome: 1 | Medicago arabica | 248 | 248 | 79% | 9e-61 | 94.51% | 71875296 | OX326964.1 |
| ☑ | Medicago lupulina genome assembly, chromosome: 4 | Medicago lupulina | 243 | 243 | 77% | 4e-59 | 93.98% | 69616069 | OY283142.1 |
| ☑ | Trifolium repens isolate ACLI19 chromosome 04_Occ | Trifolium repens | 134 | 134 | 64% | 3e-26 | 86.03% | 64312545 | CP125842.1 |
| ☑ | Trifolium repens genome assembly, chromosome: 5 | Trifolium repens | 134 | 134 | 64% | 3e-26 | 86.03% | 61923754 | OZ377887.1 |
| ☑ | Trifolium occidentale genome assembly, chromosome: 4 | Trifolium occidentale | 128 | 128 | 64% | 1e-24 | 85.29% | 73078136 | OZ060771.1 |
| ☑ | Trifolium repens genome assembly, chromosome: 11 | Trifolium repens | 124 | 124 | 64% | 2e-23 | 84.56% | 59196037 | OZ377893.1 |
| ☑ | Trifolium bocconei genome assembly, chromosome: 6 | Trifolium bocconei | 119 | 119 | 73% | 7e-22 | 81.88% | 80304874 | OZ277739.1 |
| ☑ | Trifolium bocconei genome assembly, chromosome: 6 | Trifolium bocconei | 119 | 119 | 73% | 7e-22 | 81.88% | 76392940 | OZ277836.1 |
| ☑ | PREDICTED: Vicia villosa bZIP transcription factor 11-like (LOC131632227), mRNA | Vicia villosa | 119 | 119 | 47% | 7e-22 | 88.89% | 1305 | XM_058903007.1 |
| ☑ | Trifolium repens isolate ACLI19 chromosome 04_Pall | Trifolium repens | 119 | 119 | 64% | 7e-22 | 83.82% | 59895999 | CP125843.1 |
| ☑ | Trifolium fragiferum genome assembly, chromosome: 2 | Trifolium fragiferum | 119 | 119 | 64% | 7e-22 | 83.82% | 69388287 | OX940790.1 |
| ☑ | PREDICTED: Lathyrus oleraceus bZIP transcription factor 11 (LOC127108383), mRNA | Lathyrus oleraceus | 115 | 115 | 43% | 1e-20 | 90.11% | 1365 | XM_051045846.1 |

**Slide 34**. Results of a BLASTN search for subjects similar to the separator sequence. The search returns no hit beyond four genera: *Medicago, Trifolium*, *Vicia*, and *Lathyrus* (as of 25 February 2026). The sequence from *Pisum* (Slide 31) was missed by this search. The annotation of these sequences is limited to "unknow mRNA and "bZIP transcription factor". Thus, the separator sequence is unlikely to be a remnant of a transposon.

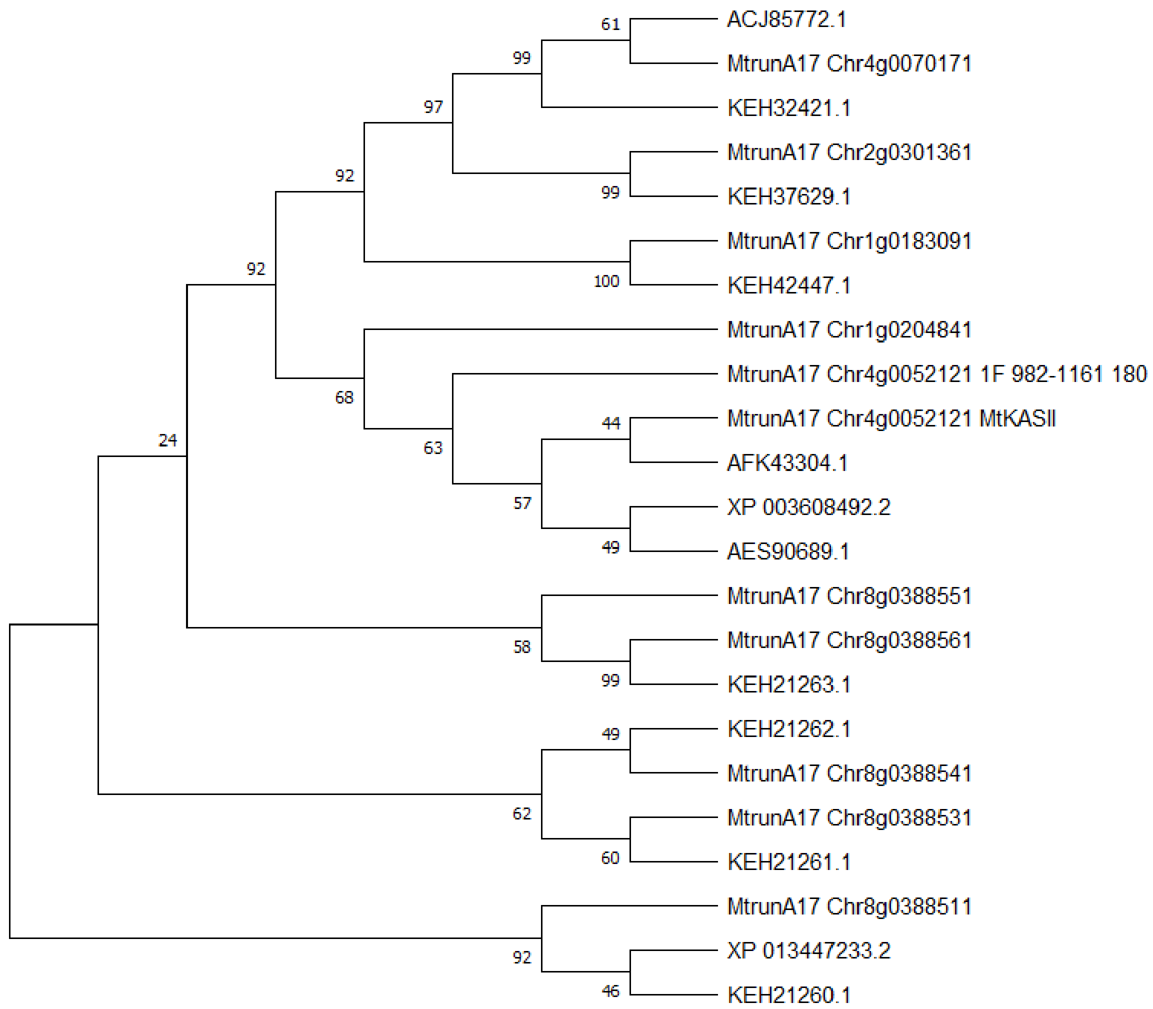


**Slide 35**. An unrooted Maximum Likelihood tree generated from a Clustal Omega alignment of MtKASII, its putative ncProt65 (MtrunA17_Chr4g0052121_1F_982-1161_180), and 21 MtKASII homologs/proteoforms from *M. truncatula*. Both MtKASII and its ncProt are related to MtrunA17_Chr1g0204841. This tree suggests that MtKASII and MtrunA17_Chr1g0204841 may have a recent common ancestor. The tree was constructed using 1000 bootstrap replicates on protein sequences without trimming. This methodological choice was conditioned by the fact that the region of interest is absent from three sequences (see the alignment details).

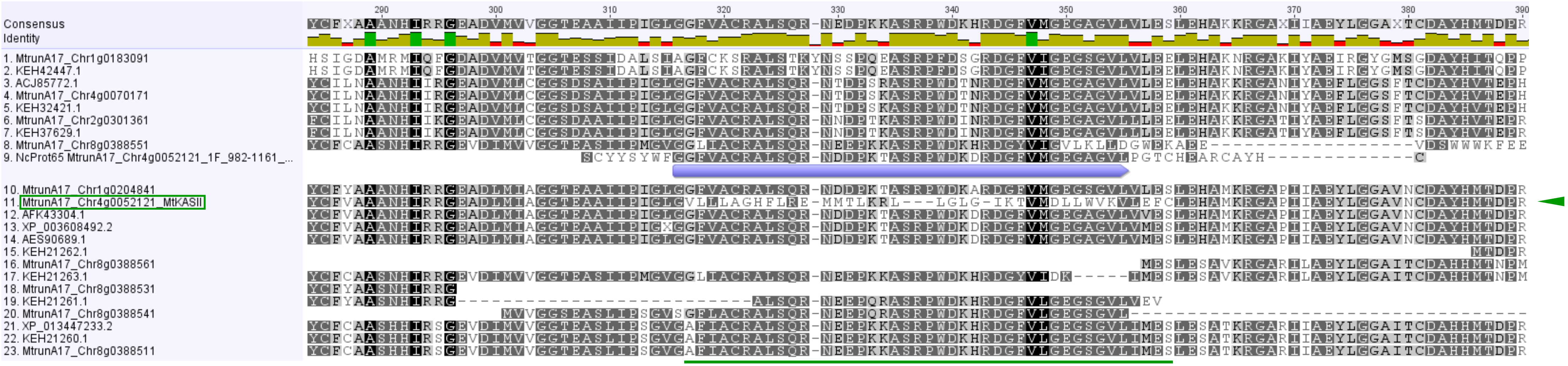


**Slide 36**. A fragment of a Clustal Omega alignment of MtKASII, its putative ncProt65 (MtrunA17_Chr4g0052121_1F_982-1161_180), and 21 putative homologs/proteoforms of MtKASII from *M. truncatula*. The same alignment was used for the tree shown in Slide 35. Among these proteins, only MtKASII contains a sequence translated from the reading frame that is alternative to the frame of all other sequences (green underlined selection). Both MtKASII and its ncProt are most similar to the refProt of MtrunA17_Chr1g0204841. This alignment suggests the very recent origin of the ncProt through two frameshifting mutations in the ancestor of *MtKASII*, which occurred in a conserved part of the sequence. The strongly conserved region of ncProt65 is shown as a violet arrow bar.

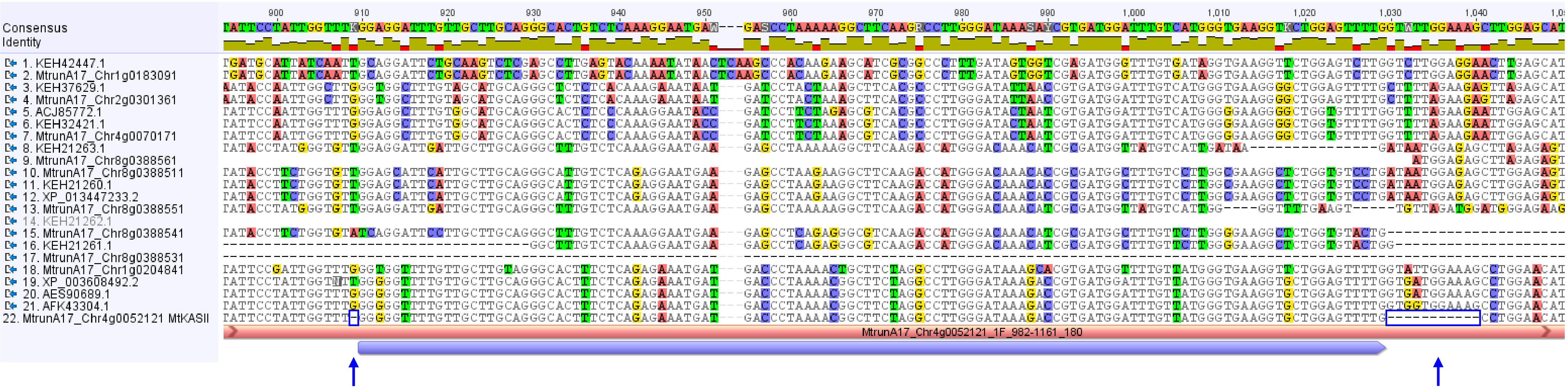


**Slide 37**. A fragment of a MUSCLE alignment of *MtKASII* CDS and 21 putative homologs/sequence variants from *M. truncatula*. The ncORF is shown as a pink bar. The strongly conserved region (violet arrow bar) was frameshifted by the deletion of one nucleotide immediately upstream of the region (blue box and vertical arrow). The reading frame was brought back to the refORF by the deletion of 11 nucleotides immediately downstream of the region (blue box and vertical arrow). This is an example of a "short round trip" mosaic protein (Çakır et al., 2023) created at the genomic DNA level without programmed ribosomal frameshifting. Namely, a short segment from the alternative frame was integrated into the strongly-conserved canonical protein, which was tolerated by natural selection.

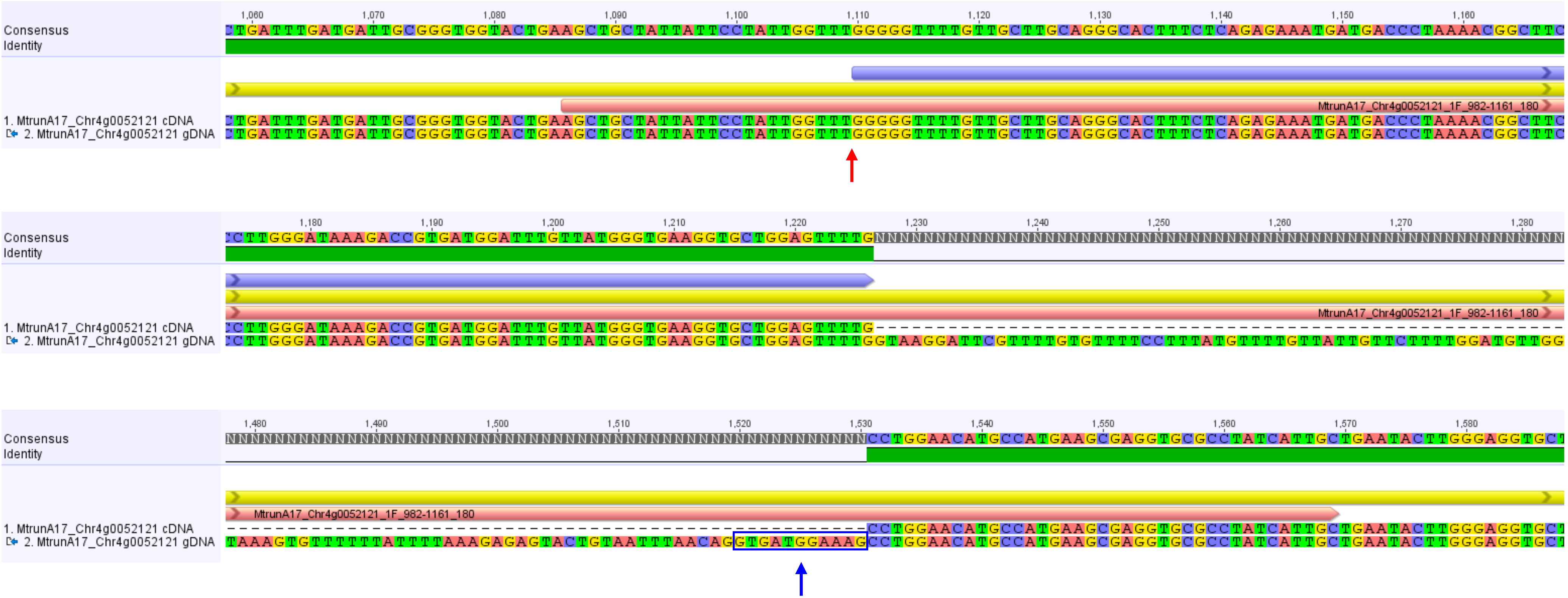


**Slide 38**. Three fragments of a MUSCLE alignment of the *MtKASII* cDNA and gDNA. The refORF, the ncORF, and the strongly conserved region are shown as yellow, pink, and violet bars/arrow bars, respectively. The top panel shows the beginning of the conserved region in Exon 2. The middle panel shows the end of that region, which coincides with the end of Exon 2. The bottom panel shows the beginning of Exon 3. While the deletion of a single G or T (red arrow) is an intra-exonic event that brought part the refORF into the alternative frame, the downstream frameshift (blue arrow) was caused by skipping the first 11 nucleotides of Exon 3 (GTGATGGAAAG). These nucleotides are still found as a part of CDS in XP_003608492.2 and AES90689.1 (see the previous slide).

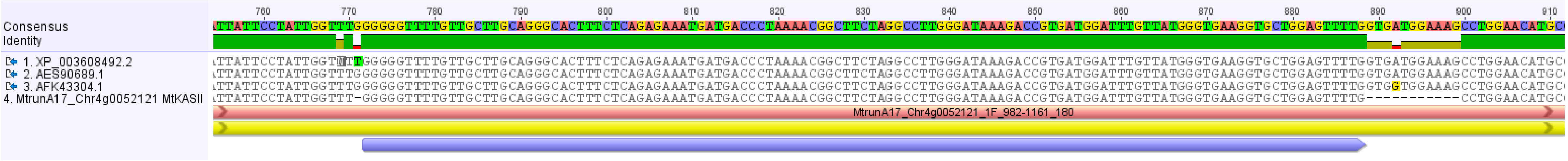


**Slide 39**. A fragment of a MUSCLE alignment of *MtKASII* CDS and three related sequences. Apart from the very few differences (visible in this fragment) upstream and downstream of the strongly conserved region (violet arrow bar), all the four sequences are identical. This suggests their origin from a single genetic locus. If the current gene model of *MtKASII* (MtrunA17_Chr4g0052121) is correct, it must have at least one non-frameshifted sequence variant similar to the gene model of MtrunA17_Chr1g0204841. Alternatively, the current frameshifted model of *MtKASII* is a sequencing artifact. Detection of genetically frameshifted MS peptides corresponding to this unusual model is necessary for its validation.

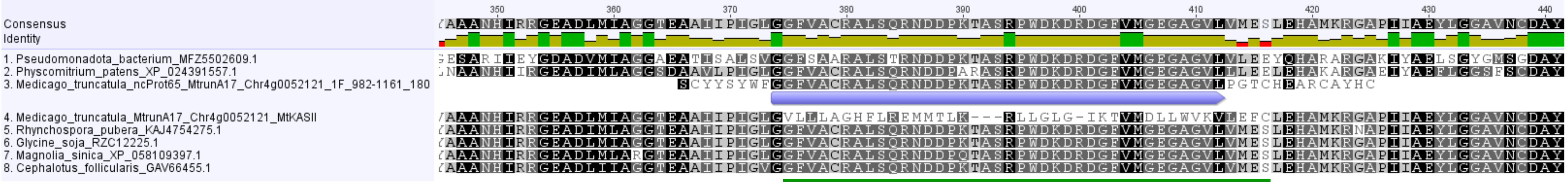


**Slide 40**. A fragment of a Clustal Omega alignment of MtKASII, its putative ncProt65 (MtrunA17_Chr4g0052121_1F_982-1161_180), and six MtKASII homologs from diverse taxa. Among these proteins, only MtKASII contains a sequence translated from the reading frame alternative to the frame of all other sequences (green underlined selection). The alignment shows that the ncProt evolved in a region conserved not only in embryophytes but also between embryophytes and bacteria (violet arrow bar). Represented taxonomic groups: *Glycine soja* – eudicots, legumes; *Cephalotus follicularis* – eudicots, non-legumes; *Rhynchospora pubera* – monocots; *Magnolia sinica* – magnoliids; *Physcomitrium patens* – bryophytes; *Pseudomonadota* bacterium – bacteria.

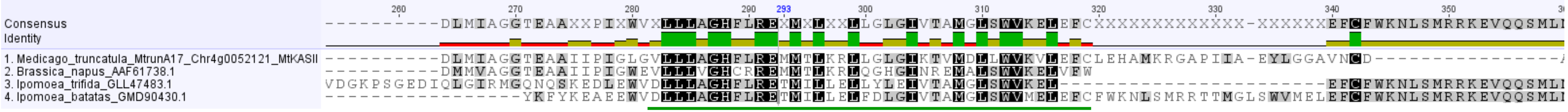


**Slide 41**. A fragment of a Clustal Omega alignment of MtKASII and three similar sequences from non-legume eudicots. The underlined region is translated from the reading frame that is alternative to the frame of all other homologs of MtKASII. There are no other sequences similar to this region in the NCBI non-redundant protein database (as of 25 February 2026). MtKASII and the *Brassica napus* sequence annotated as beta-ketoacyl-ACP synthetase 1 have 72% overall amino acid identity (66% in the underlined region). Two *Ipomoea* sp. sequences are annotated as unrelated proteins: bidirectional sugar transporter SWEET1-like (*Ipomoea trifida*) and ribonucleoside-diphosphate reductase large subunit-like (*Ipomoea batatas*). They have very low overall percent identity with MtKASII (14% and 11%, respectively). This suggests that the underlined sequence evolved independently in *B. napus* by a simpler frameshifting mutation, as follows from no return to the original frame and early termination of the sequence in this species.

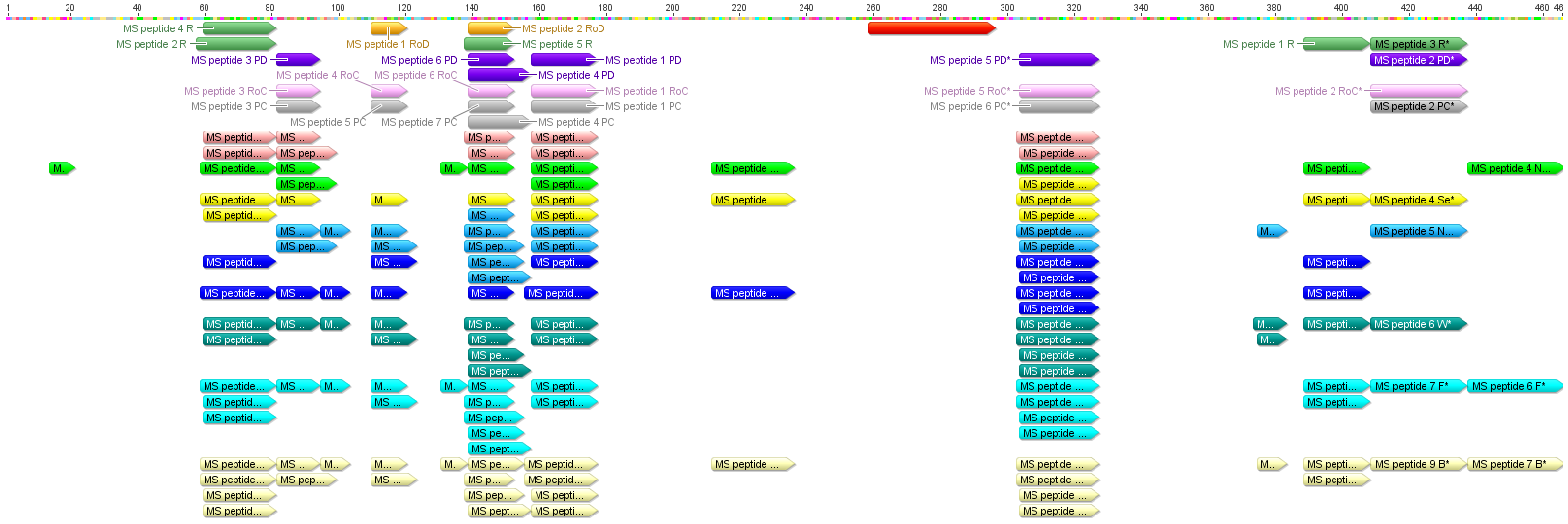


**Slide 42**. Mapping of MS peptides to the refProt of *MtKASII*. The region translated from the alternative frame is shown in red. It is not covered by any MS peptide from 16 different samples and three independent studies (PXD002692, Marx et al., 2016; PXD013606, Shin et al., 2021; PXD022278, Castañeda et al., 2021). Because our detection pipeline had no bias against MS peptides that match refProt sequences, this fact may indicate that the current gene model of *MtKASII* may be invalid. Different colors correspond to different proteomic samples. No peptide matching the refProt of *MtKASII* was identified in 28-day-old root nodules, phloem control, and phloem under drought conditions. Peptide counts correspond to the actual (redundant) occurrence of validated sequences in peptide reports generated during the search for chimeric peptides (Çakır et al., 2025b, Supplementary Dataset S36, folder Modelled_with_conserved_altProts). Peptide names ending with an asterisk are shared with the refProt of MtrunA17_Chr1g0204841. The map in Geneious® format is available in Data S17.

**Data S16**. Sequence analysis demonstrating the origin of highly conserved ncORFs in five selected genes. The dataset contains the information on ncORFs from transcripts of *MtTPST*, *MthGSHSb*, *MtPHO2-A*, *MtbZIP60b*, and *MtKASII*. In these genes (except for *MtTPST*) ncORFs are conserved stronger at the amino acid level compared to the nucleotide level. Graphical data were generated in Geneious® R7.1.9. Phylogenetic trees were constructed in MEGA-X v10.2.6. No phylogenetic tree was built for *MtTPST* and *MtPHO2-A* because *MtTPST* has no homolog in *Medicago truncatula*, and *MtPHO2-A* has only two homologs. All alignments in Geneious® and MEGA formats are available in Data S17.

**Contents**

Slides 1-5: *MtTPST*
Slides 6-15: *MthGSHSb*
Slides 16-25: *MtPHO2-A*
Slides 26-34: *MtbZIP60b*
Slides 35-42: *MtKASII*
