## Supplementary material for "Loss-of-function phenomics, ncORFs, and ambiguity of mutant phenotypes in *Medicago truncatula*": Data S17 Long alignments: Legend.docx

**Data S17**. Nucleotide alignments used in Data S16. MUSCLE alignments were constructed in Geneious® R7.1.9. Clustal Omega alignments were imported to Geneious® for visualization. All alignments were exported in two formats: Geneious® and MEGA. Geneious® files preserve annotation features shown in Data S16. The dataset also contains a map of MS-validated peptides on the refProt of *MtKASII*.
