## Supplementary material for "Loss-of-function phenomics, ncORFs, and ambiguity of mutant phenotypes in *Medicago truncatula*": Data S18 Gene and altORF models: Legend.docx

**Data S18**. Gene models and mapped ncORFs of 123 loci targeted by functional analyses in *Medicago truncatula* (Data S12). Geneious® files include CDS, cDNA (with mapped ncORFs), and gDNA sequences (v5.1.9) along with cDNA/gDNA alignments for each gene. Part 1 contains Tier 1 and Tier 2 sequences. Part 2 contains Tier 3 sequences. The files were generated in Geneious® R7.1.9 and can be viewed in any version of this software starting with version R7. NcORFs are referred to as altORFs (for alternative ORFs, Mouilleron et al., 2016) in these Geneious® files because the decision to adopt the naming convention of Mudge et al. (2022) was made after the generation of this dataset.
