## Supplementary figures and images for "Loss-of-function phenomics, ncORFs, and ambiguity of mutant phenotypes in *Medicago truncatula*"

### Graphical abstract

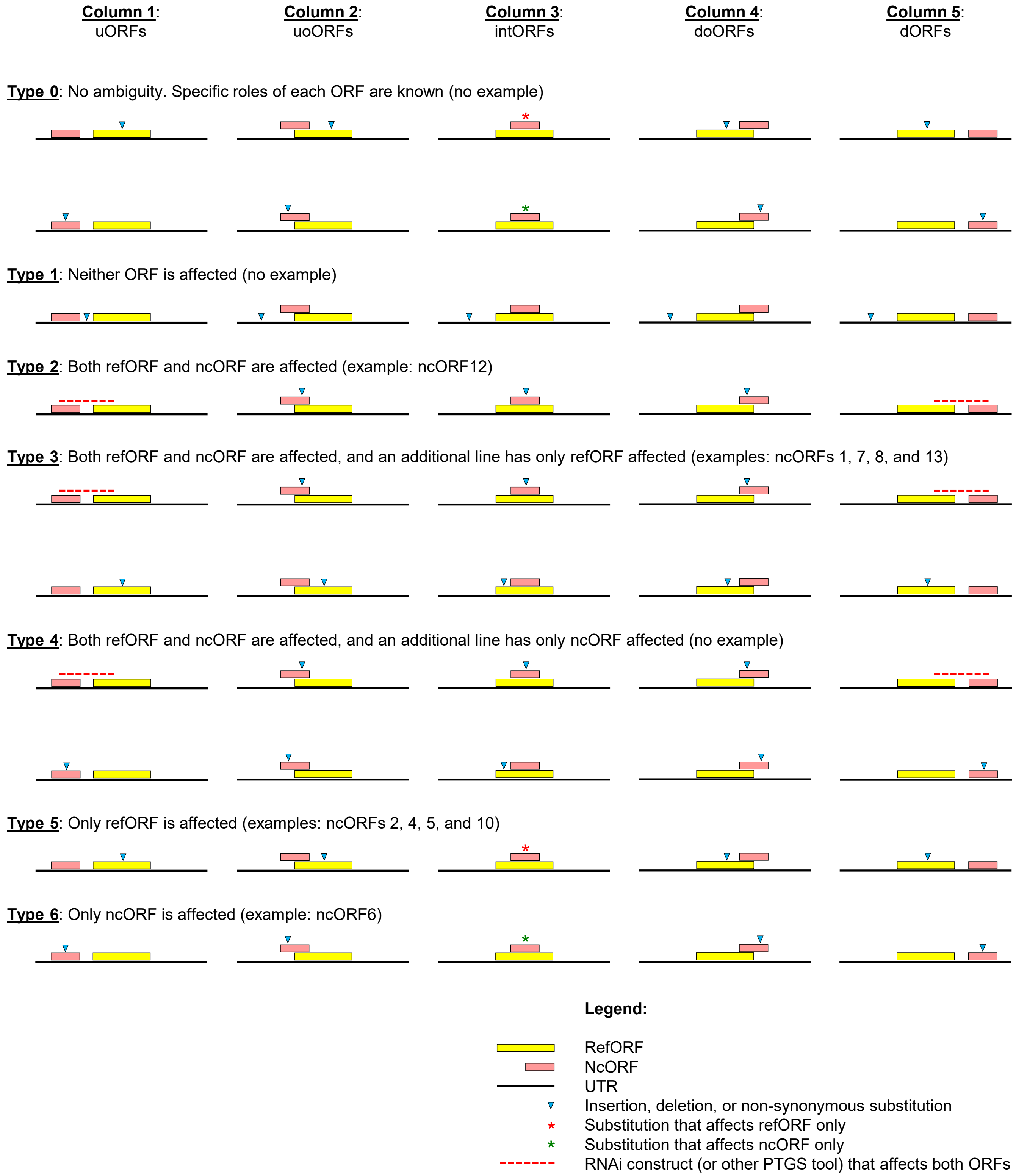
