## Supporting Discussion for "Loss-of-function phenomics, ncORFs, and ambiguity of mutant phenotypes in *Medicago truncatula*"

#### Unique proportions of altered, conditional, and neutral phenotypes in different groups of *M. truncatula* genes may reflect their relative biological importance

Our meta-analysis of loss-of-function phenotypes in *M. truncatula* revealed that different groups of genes have unique proportions of altered (A), conditional (C), and neutral phenotypes (N). These proportions can significantly deviate from the overall proportion in the dataset (62:27:11, respectively) and from the proportions of individual groups (Data S10 and S11). Not only specific protein families but also combined groups such as transcription factors, enzymes, transporters, channels, other membrane proteins, and peptides exhibit unique proportions of phenotypes. This information can be very useful for functional genomics specialists and breeders of legume crops because it can help focus on candidate genes that are likely to give scorable non-conditional phenotypes.

We attempted to compare this information with data from the major loss-of-function databases cited in the main text (Groth et al., 2007; Lloyd and Meinke, 2012; Courtier-Orgogozo et al., 2020; Baldarelli et al., 2024). None of these studies reported tolerance to mutations for protein classes comparable to those shown in Data S10. Using *A. thaliana*, Lloyd and Meinke (2012) addressed this aspect differently. Genes were organized in 11 classes based on protein function (Figure 5 in Lloyd and Meinke, 2012). However, those classes are heterogeneous with regard to gene families. The only meaningful comparison that can be made between this study and our data concerns transcription factors. Specifically, Lloyd and Meinke (2012) mentioned overrepresentation of transcription factors among genes with morphological phenotypes and their depletion among genes with embryo-lethal phenotypes. This underlines the role of transcription factors in fine-tuning of organismal functions. In our study, transcription factors have the second lowest proportion of A-phenotypes (53%) among combined groups (Data S11), which somewhat contrasts with the *A. thaliana*. At the same time, among 11 genes with the terms “lethal” or “lethality” in phenotype columns of Data S6, there are three transcription factors (embryo lethality, *MtBTS1*; seed lethality, *MtDASH*; seedling lethality, *MtFE*), which does not indicate the overall underrepresentation of this group among lethal phenotypes. The small number of genes with lethal phenotypes in our study, along with the difficulty of discriminating between embryo and gametophyte lethality when *Tnt1* mutants cannot be isolated, precludes accurate comparisons with the data of Lloyd and Meinke (2012).

Although the direct comparison with other studies cannot be made due to the difference in approaches (e.g., different grouping of genes and phenotypes; no tripartite stratification used, such as A:C:N in our study), similar observations were made in humans. Caldu-Primo et al. (2021) grouped human genes into tolerant and intolerant to mutations and considered two groups of phenotypes based on whether they concerned organismal fitness or cellular viability. At the same time, their grouping of protein classes was somewhat similar to ours (Figure 5 in Caldu-Primo et al., 2021). Significant deviations from random expectations were found in each protein class. For example, kinases and transcription factors were overrepresented in the category of intolerant phenotypes that affected organismal fitness. However, they were underrepresented in this category when the phenotypes concerned cellular viability. In our study, kinases have a relatively high proportion of A-phenotypes (71%), but transcription factors do not (Data S10 and S11). Their proportion of A-phenotypes (53%) is the second lowest among combined groups. In contrast, transporters have the second highest proportion of A-phenotypes among combined groups in our study (71%) but are not prominent with regard to their tolerance to mutations in the human study. The same concerns the combined group of enzymes that are relatively intolerant to mutations in our meta-analysis (66% of A-phenotypes) but are close to the random expectation in Caldu-Primo et al. (2021).

A recent study in humans used a different approach (Hasenahuer et al., 2023). Evolutionarily Constrained Coding Regions (CCRs), where protein-changing variants are rare in healthy individuals, were mapped to proteins they encode. This analysis revealed protein classes over- and underrepresented among CCRs (13 and 12 classes, respectively; Figure 6 in Hasenahuer et al., 2023). Only three of those classes were the same as in our analysis (transcription factors, defensins, and oxidoreductases). Transcription factors were moderately but significantly overrepresented among CCRs in Hasenahuer et al. (2023). In our study, they are among the lowest concerning the proportion of A-phenotypes (53%) (Data S11). Defensins and oxidoreductases are among the underrepresented classes in Hasenahuer et al. (2023) and in our dataset (Data S10). This brief comparison with other studies suggests that tolerance to mutations in different protein groups may be species-specific, although certain overlaps among different taxa are observed.

Beyond the three studies discussed here (Lloyd and Meinke, 2012, Caldu-Primo et al., 2021, and Hasenahuer et al., 2023), little data is available on differential tolerance to mutations among different protein classes and families. In contrast, many studies (including Lloyd and Meinke, 2012, Caldu-Primo et al., 2021, and Hasenahuer et al., 2023) report such differential tolerance among genes involved in different biological processes (for example, Tzafrir et al., 2004; Wang et al., 2015; Rubin et al., 2015; Hart et al., 2015; Tsherniak et al., 2017; Li et al., 2019; Rossiter et al., 2021). Quite expectedly, the common theme of those studies is essentiality of genes involved in transcription, translation, RNA processing, and ribonucleoprotein complex biogenesis. Because these studies address diverse aspects of essentiality and use different systems, no consensus exists about processes such as DNA replication, DNA repair, cell cycle, ATP binding, kinase activity, signaling passways, mitochondrial functions, metabolism, energy production, and transmembrane transport.

It should be noted that gene essentiality is not the same as the probability to exhibit a scorable phenotype, which is addressed in our study. Gene essentially has been a topic of intensive research. This field of studies has advanced so far as to predict which genes are likely to be essential (Lloyd et al., 2015). For example, genes with paralogs tend to be not essential for viability in animals (Chen et al., 2012; White et al., 2013). However, there are many essential duplicated genes involved in development (Chen et al., 2012). Developmental genes are clearly more essential than non-developmental ones in animal systems (Chen et al., 2012). In plants, the presence of multigene families makes the situation more complex. Among gene families with the largest proportions of A-phenotypes, we found families with hundreds of homologs in *M. truncatula* (Data S10 and S11). For example, transcription factors of the C2H2 family, cytochromes P450, and transcription factors of the MYB-HB and bZIP families have 643, 403, 146, and 87 homologs, respectively, in the latest genome annotation. Some studies emphasized the conditionality of essential genes (Rossiter et al., 2021 Caldu-Primo et al., 2021) and introduced a concept of bypassable gene essentiality (Li et al., 2019). Briggs et al. (2006) reviewed examples where the strength of phenotypes among homologous plant genes is highly unequal (the phenomenon of unequal redundancy). One member of a given multigene family can be essential, but a closely related member of the same family can be redundant. This observation suggests that no statistically supported family-specific proportions of phenotypes (A:C:N in our study) should be observed. This discrepancy indicates that the message of our meta-analysis deserves special attention. Very few other studies provided family-wise information on the relative tolerance to mutations (Lloyd and Meinke, 2012, Caldu-Primo et al., 2021, Hasenahuer et al., 2023). The rarity of such data for diverse evolutionarily related classes and families of proteins outlines our resource as quite unique. Moreover, Data S6 can be further mined for phenotype proportions specific to various biological processes. Such analysis can help plant researchers choose candidate genes that are likely to show a scorable phenotype in a specific process of interest, such as SNF, leaf development, or drought stress.

**Some currently annotated gene models in *M. truncatula* should be critically re-evaluated based on original loss-of-function studies and our ncORF data**

Many old gene models were split into two in the current genome annotation (v. 5.1.9). Table S2 lists five examples of such models mentioned in the Results section. Four of them were targeted by loss-of-function studies under the assumption of a single locus (*MtGSHS1*, *MtLICK1*, *MtROP7*, and *MtSYT1*). The fifth one, Medtr3g117120 (bZIP transcription factor), has not been studied yet. Evidence provided by the original functional studies and/or sequence alignments in Data S16 (see Supporting Results) suggests that the old models may be correct for those five examples. Many more instances of this type can be found in the current genome annotation, which may require critical re-evaluation.

**Table S2**. Five v. 4 gene models that may be correct without splitting implemented in genome annotation v. 5.1.9

| Locus ID v. 4 | Original locus name | Locus ID v. 5.1.9 | New locus name | Reference |
|---|---|---|---|---|
| Medtr7g113890 | *MtGSHS1* | MtrunA17_Chr7g0273151 | *MtGSHSa* | Frendo et al., 2001, 2005; Data S16 Slides 7-9 and 13 |
| | | MtrunA17_Chr7g0273161 | *MtGSHSb* | Frendo et al., 2001, 2005; Data S16 Slides 7-9 and 13 |
| Medtr1g102190 | *MtLICK1* | MtrunA17_Chr1g0204221 | *MtLICK1a* | Wang et al., 2025 |
| | | MtrunA17_Chr1g0204231 | *MtLICK1b* | Wang et al., 2025 |
| Medtr4g088055 | *MtROP7* | MtrunA17_Chr4g0046211 | *MtROP7a* | Wang et al., 2021b |
| | | MtrunA17_Chr4g0046221 | *MtROP7b* | Wang et al., 2021b |
| Medtr4g073400 | *MtSYT1* | MtrunA17_Chr4g0037111 | *MtSYT1a* | Gavrin et al., 2017 |
| | | MtrunA17_Chr4g0037121 | *MtSYT1b* | Gavrin et al., 2017 |
| Medtr3g117120 | NA (bZIP transcription factor) | MtrunA17_Chr3g0144931 | NA (bZIP transcription factor) | Data S16 Slides 28-30 |
| | | MtrunA17_Chr3g0144941 | NA (bZIP transcription factor) | Data S16 Slides 28-30 |

"NA" stands for "not available" (the locus has not been studied and named yet).

Some v. 4 gene models were duplicated in the latest annotation. This is exemplified by the v. 4 gene Medtr4g125520 (*MtMYB14*). Two entries in the genome assembly v. 5.1.9 correspond to this locus: MtrunA17_Chr4g0070611 and MtrunA17_Chr4g0070641. However, the original data presented by Liu et al. (2014) suggest that only one of the loci exists in the *M. truncatula* ecotype R108, specifically, the long version MtrunA17_Chr4g0070611 (see the Results section). This case is one of many that emphasize the challenge of functional annotation in the ecotype Jemalong A17 based on the data from the ecotype R108. Ideally, the genome browser should highlight ecotype-specific loci and specify the source of mutant data for each locus targeted by a loss-of-function approach.

**The specific role of ncORFs in correcting current gene models**

NcORF analysis is usually not regarded as a tool that can help rectify gene models. In Data S16, we provided four examples of how the MS validation and conservation analysis of ncORFs can point to flaws in the latest gene models. In genes *MtTPST*, *MthGSHSb*, *MtPHO2-A*, and *MtKASII*, the existence of conserved ncORFs at specific positions revealed the possibility that the current models are incorrect. In these models, regions that appear as conserved/translated ncORFs may be part of actual refORFs. These regions are disconnected from the annotated refORFs or moved to an alternative frame (*MtKASII*) by events such as alternative splicing and possibly sequencing artifacts. At the current stage of the genome assembly, sequencing errors are extremely improbable. But can they be entirely ruled out as the source of unusual gene models such as *MtKASII*? This question deserves an in-depth reexamination of sequencing data, preferably using alternative methods. Among the five genes shown in Data S16, only the model of *MtbZIP60b* is likely to be correct. However, even in this gene, the ncORF is a remnant of an ancient longer refORF (Data S16 Slides 28 and 31). This type of analysis helps understand the evolutionary origins of ncORFs.

It should be noted that the current models of some genes strongly differ from those described in the original functional studies. *MtPHO2-A* is the most remarkable example of this type. However, *MtTPST* and *MthGSHSb* also fall into this category (Data S12). In these cases, the old models are supported by our ncORF analysis.

These examples underline the importance of considering ncORFs in all genome annotation efforts. Even without evidence for translation, conservation signatures of ncORFs help rectify gene models and understand the evolution of coding regions.

**Conservation signatures of ncORFs cannot be used as reliable markers of their translation status**

Quite reasonably, conservation of ncORFs has traditionally been used as a proxy for translation evidence (Vanderperre et al., 2012, 2013; Kochetov et al., 2017; Samandi et al., 2017; Brunet et al., 2019, 2021; Pavesi, 2025), also in our earlier work (Çakır et al., 2025). However, considerations presented in Data S16 call for caution in using purifying selection signatures of ncORFs as evidence for their translation. *MtKASII* exemplifies the need to look closer into the origin of each conserved ncORF (Data S16 Slides 35-42). NcORF65 in this gene is super-conserved and under purifying selection. However, this conservation signature may simply reflect the misannotation of this region in the current gene model. Specifically, ncORF65 may be part of the refORF of *MtKASII*. Alternatively, if the current model is correct, relatively recent evolutionary events may have moved this highly conserved segment of an ancestral refORF into an alternative frame. In such a scenario, ncORF65 may have preserved the purifying selection signature because obliterating this signature requires longer time. Realizing this limitation of the conservation analysis does not make it less useful. On the contrary, such analysis of ncORFs can point out events overlooked by functional studies. Hence, large-scale profiling of ncORF conservation signatures like the one conducted in our study can improve the accuracy of interpretation of loss-of-function phenotypes.

**Large-scale conservation analysis of ncORFs can be facilitated by the comparison of BLASTP and BLASTN hit numbers**

As indicated above, conservation profiling of all potential ncORFs should be an integral part of a truly comprehensive transcriptome annotation. However, a transcriptome-wide search for ncORFs that are under purifying selection is not a trivial task. We propose that a simple comparison of hit numbers of ncORF sequences at the amino acid level and at the nucleotide level can help narrow down the candidate list of truly conserved ncORFs. In our analysis, 28 ncORFs in functionally characterized genes (out of 140) showed larger numbers of hits at the amino acid level (Data S12). Among them, 11 are under purifying selection (Data S13 and S14). This indicates that the relative abundance of DIAMOND-based BLASTP hits may be used as a proxy for strong conservation at the amino acid level. Highly conserved ncORFs shown in Data S16 were detected using this approach.

**Taxonomic ranges of BLASTP hits present a valuable framework for the detection of super-conserved ncORFs**

Not only the number of hits but also the range of taxa in which these hits are found can reflect the conservation of ncORFs at the amino acid level. In contrast to automated similarity searches such as BLASTP searches conducted with DIAMOND, deducing taxonomic ranges of hits from such data is a major computational challenge. It is conditioned by several factors. First, many entries in protein databases such as UniProt and GenBank are not supplied with proper taxonomic tags. Thus, in the first step, such information must be imported from alternative sources, for example, publicly accessible web resources. The next difficult step is to convert the wealth of information on individual taxonomic groups for all hits of a given ncProt into a single line describing the taxonomic range in a uniform way. In our study on 18,643 ncORFs with at least one significant hit (Supplementary Figure S1 in Çakır et al., 2025), we automated the first step. However, we were unable to automate the second step. Consequently, taxonomic ranges were curated manually. This manual effort paid off because it helped identify ncORFs with hits across all or nearly all major branches of the tree of life (e.g. ncORF53 in *MtPHO2-A*, ncORF65 in *MtKASII*, and ncORF12 in *MthGSHSb*). In this manuscript, we revealed the taxonomic ranges of only 140 ncORFs (Data S12). The information on taxonomic ranges of all 18,643 ncORFs will be disclosed later, after the publication of the entire set of ncORFs. NcORFs with signatures of purifying selection will be integrated into the genome browser of *M. truncatula*. In view of the technical difficulty of generating such information and its immense biological relevance, we suggest that computational efforts should be dedicated to automating this process. To address this need, we have developed a dedicated pipeline that will be published elsewhere. Furthermore, once the automation is possible, we suggest that genomic browsers of all organisms should integrate such information as part of the standard annotation of transcriptomes.

**Many ncORFs will consistently remain undetected with MS proteomics because of regions 100% identical to refProts**

During our conservation analysis of ncProts, we became aware of one serious bias universally applicable to ncORF validation using MS proteomics. Specifically, we noticed that many ncProts in our dataset contain regions longer than 6 aa that are 100% identical to GenBank entries from *M. truncatula* (Data S12 Columns W and X). While only a fraction of these subject sequences corresponds to annotated refProts in the current genome annotation of *M. truncatula* (Data S12 Columns Y and Z), this observation indicates that certain translated ncProts will never be validated with MS proteomics because the detection pipeline will misinterpret perfectly matching peptides as products of refORFs. This, of course, will depend on the type of digestive enzymes used for preparing the MS samples. However, certain ncProts will remain undetectable even in studies that use multiple enzymes or enzyme-free approaches because their sequences may not generate peptides that are detectable or uniquely identifiable by MS. This observation outlines a group of ncORFs that may deserve special attention and require special methods for validating their translation. As long as such ncORFs remain overlooked, functional annotation of a genome remains incomplete. To exemplify this challenge, we have listed 29 ncProts grouped with refProts in our proteomic analysis (Data S19). Earlier, we found 805 MS-validated non-chimeric ncProts in three proteomic datasets (Çakır et al., 2025). Those 29 ncProts would be added to the list of 805 MS-validated ncProts if they had no MS peptides 100% identical to refProts. This corresponds to only 3.6% of 805 MS-validated ncProts. However, this number is sufficiently large to make the research community aware of this intrinsic limitation of the standard MS proteomic analysis, when it comes to the detection of translons never anticipated before. Truly comprehensive functional analysis cannot ignore the existence of such undetectable and conditionally detectable ncORFs.

**NcProts with similarity restricted to *M. truncatula* sequences present an unexplored realm of potentially functional entities**

In the previous section, we described hard-to-detect ncProts that have regions of 100% identity to refProts of *M. truncatula*. Such ncProts can be further subdivided into two subgroups. The first group has hits among *M. truncatula* refProts along with refProts of other organisms. The second group, however, has 100% identity regions exclusively with *M. truncatula* refORFs. Traditionally, such intra-species conservation is not considered as biologically relevant. Thus, ncORFs conserved within one species are disregarded as non-conserved. This view, however, should be changed because when such ncORFs are translated, they may remain undetected by MS proteomics. A short region of an intact refORF may be duplicated through various mechanisms, including transposition. Such a region retains the protein-coding potential as a functional module that can be quickly incorporated into the genetic space of other, even unrelated, genes. However, during this incorporation, the reading frame of the duplicated segment may not match the refORF of the locus that received the duplicated sequence. Thus, while some of such duplications become part of refORFs, others will be incorporated into alternative reading frames. There is no known mechanism to prevent their translation, if such translation is beneficial for reproductive success. Artificially excluding such ncORFs from the pool of potential translons is an omission that hinders the progress of functional studies.

**NcORFs unify alternative splicing and mosaic translation**

One possible reason for so many conserved ncORFs found in a single transcriptome is their function as alternative exons. The same region of a pre-mRNA can be translated in different reading frames in the mature transcript, depending on how many nucleotides separate it from the translational start. Some sequences code for proteins in all reading frames. This property probably reflects the evolutionary "readiness" of such ncORFs to participate in translation in scenarios where the canonical reading frame is disrupted or does not provide the required product. Because of these considerations, it is evident that the role of ncORFs in alternative splicing must be the same as in the mosaic translation, which is a hypothetical mechanism proposed by our group based on earlier studies (Çakır et al., 2023, 2025). Specifically, ncORFs can be used as building blocks, or modules, that diversify the amino acid sequence of long proteins. This can be achieved via alternative splicing or via programmed ribosomal frameshifting. Thus, ncORFs unify these two processes as alternative ways to achieve the same goal: to produce proteins that are mosaic with regard to their domain composition. Conserved ncORFs, even with conservation limited to the host species, must be especially fit for this role of modules because they have been evolutionarily tested for protein coding. In Data S16, alternative splicing emerges as the major mechanism explaining the high degree of conservation found in some ncORFs. Thus, our data indirectly support the view on alternative splicing and programmed ribosomal frameshifting as two faces of the same phenomenon: biologically advantageous chimerism at the amino acid level, which promotes adaptation to the variety of internal and external conditions. For the broader discussion about the relevance of ncORFs to alternative splicing, and their joint contribution to the proteome, we refer the reader to the review by Manuel et al. (2023).

**MS-validated ncORFs introduce ambiguity into the interpretation of loss-of-function studies**

In contrast to conserved ncORFs, which may or may not be translated as discrete units, MS-validated ncORFs are likely to be translated either as independent translons or as part of refORFs in alternative splicing variants. When ignored, such sequences can compromise the correct interpretation of loss-of-function phenotypes, which are traditionally attributed to refORFs. We have recognized seven possible types of ambiguity associated with translated ncORFs in target genes (Figure 4). Four of them are present in our dataset of 10 MS-validated ncORFs under the assumption that the current gene models are correct (Table S3). The first type corresponds to zero-ambiguity (hence, Type 0), which indicates 100% clarity as to specific roles of a refORF and a ncORF. Types 1 and 2 represent complete ambiguity as to the relative contribution of the refORF and the ncORF into the loss-of-function phenotype. Types 3-6 represent incomplete ambiguity. We believe this classification is conceptually comprehensive. Thus, we hope it can be adopted by the research community.

Type 0 ambiguity

This type corresponds to no ambiguity: specific roles of the refORF and the ncORF are known. This requires affecting one ORF without changing the amino acid sequence or downregulating the other ORF. Achieving such level of clarity requires special tools and precautions (see our suggestions in a dedicated section below).

Type 1 ambiguity

Neither the refORF nor the ncORF are affected. This type represents complete ambiguity as to the relative contribution of each ORF into the loss-of-function phenotype. Specifically, the phenotype cannot be assigned to either the refORF or the ncORF without a dedicated study that mutagenizes them independently. This type can reasonably be expected to occur only rarely, for example, when an insertion or a small deletion is between the refORF and the ncORF. While such mutants are usually not considered informative for functional studies, this type of ambiguity should be included in a holistic classification, which is the aim of the present article. This type of ambiguity may be encountered when a researcher wrongfully assumes that the UTR was part of the refORF, for example, due to a mistake in the annotation of the coding region. Expectedly, Type 1 ambiguity has no corresponding MS-validated ncORF in our dataset.

Type 2 ambiguity

This type is opposite to Type 1. Both the refORF and the ncORF are affected, and no additional mutant alleles are available. This type represents complete ambiguity as to the relative contribution of each ORF into the loss-of-function phenotype. Specifically, the phenotype cannot be assigned to either the refORF or the ncORF without a dedicated study that mutagenizes them independently. This type is exemplified only by ncORF12.

Type 3 ambiguity

It is similar to Type 2. Both the refORF and the ncORF are affected in one line but only the refORF is affected in an additional line. This type represents incomplete ambiguity as to the relative contribution of each ORF into the loss-of-function phenotype. Specifically, the true loss-of-function phenotype of the refORF is known. However, the true loss-of-function phenotype of the ncORF should be assessed independently of the refORF. It may be different from the phenotype observed in the line affected in both ORFs because mutant proteins can interact. They can either cancel out the effect of independent mutations or can have a synergistic effect not observed in individual mutants. Type 3 is exemplified by ncORFs 1, 7, 8, and 13.

Type 4 ambiguity

It is similar to Types 2 and 3. Both the refORF and the ncORF are affected in one line but only the ncORF is affected in an additional line. This type represents incomplete ambiguity as to the relative contribution of each ORF into the loss-of-function phenotype. Specifically, the true loss-of-function phenotype of the ncORF is known. However, the true loss-of-function phenotype of the refORF should be assessed independently of the ncORF. It may be different from the phenotype observed in the line affected in both ORFs because mutant proteins can interact. They can either cancel out the effect of independent mutations or can have a synergistic effect not observed in individual mutants. Type 4 ambiguity has no corresponding MS-validated ncORF in our dataset.

Type 5 ambiguity

It is somewhat similar to Type 3. Only the refORF is affected. The ncORF is not affected. This type represents incomplete ambiguity as to the relative contribution of each ORF into the loss-of-function phenotype. Specifically, the true loss-of-function phenotype of the refORF is known. However, the loss-of-function phenotype of the ncORF is unknown. Thus, it should be assessed independently of the refORF. Type 5 is exemplified by ncORFs 2, 4, 5, and 10.

Type 6 ambiguity

It is opposite to Type 5 and somewhat similar to Type 4. Only the ncORF is affected. The refORF is not affected. This type represents incomplete ambiguity as to the relative contribution of each ORF into the loss-of-function phenotype. Specifically, the true loss-of-function phenotype of the ncORF is known. However, the loss-of-function phenotype of the refORF is unknown. Thus, it should be assessed independently of the ncORF. Type 6 is exemplified by ncORF6.

**Table S3**. MS-validated ncORFs that introduce ambiguity into the interpretation of loss-of-function studies

| NcORF | Locus name | Reference | Ambiguity type and description |
|---|---|---|---|
| NcORF1 | *MtVPT2* | Liu et al., 2023 | Type 3. Both the refORF and the ncORF are affected in one line but only the refORF is affected in an additional line. Regardless of the phenotypic difference between the two lines, the true LOF phenotype of the ncORF should be assessed independently of the refORF. |
| NcORF2 | *MtREV1* | Zhou et al., 2019 | Type 5. Only the refORF is affected. The LOF phenotype can be attributed exclusively to the refORF. The ncORF should be disrupted independently of the refORF in a dedicated study. |
| NcORF4 | *MtCKX6* | Wang et al., 2021a | Type 5. Only the refORF is affected. The LOF phenotype can be attributed exclusively to the refORF. The ncORF should be disrupted independently of the refORF in a dedicated study. |
| NcORF5 | *MtWD40-1* | Pang et al., 2009; Meng et al., 2019, 2023 | Type 5. Only the refORF is affected. The LOF phenotype can be attributed exclusively to the refORF. The ncORF should be disrupted independently of the refORF in a dedicated study. |
| NcORF6 | *MtTPST* | Zhang et al., 2025 | Type 6. Only the ncORF is affected. The LOF phenotype can be attributed exclusively to the ncORF. The refORF should be disrupted independently of the ncORF in a dedicated study. |
| NcORF7 | *MtAHL1* | Zhang et al., 2023 | Type 3. Both the refORF and the ncORF are affected in one line but only the refORF is affected in an additional line. Regardless of the phenotypic difference between the two lines, the true LOF phenotype of the ncORF should be assessed independently of the refORF. |
| NcORF8 | *MtCAS31* | Li et al., 2018 | Type 3. Both the refORF and the ncORF are affected in one line but only the refORF is affected in an additional |

| | | | |
|---|---|---|---|
| | | | line. Regardless of the phenotypic difference between the two lines, the true LOF phenotype of the ncORF should be assessed independently of the refORF. |
| NcORF10 | *MtSCR* | Dong et al., 2021 | Type 5. Only the refORF is affected. The LOF phenotype can be attributed exclusively to the refORF. The ncORF should be disrupted independently of the refORF in a dedicated study. |
| NcORF12 | *MthGSHSb* | Frendo et al., 2001, 2005 | Type 2. Both the refORF and the ncORF are affected (no additional mutant alleles are available). The LOF phenotype cannot be assigned to either of them without a dedicated study that mutagenizes the refORF and the ncORF independently. |
| NcORF13 | *MtMYC2* | Guo et al., 2024 | Type 3. Both the refORF and the ncORF are affected in one line but only the refORF is affected in an additional line. Regardless of the phenotypic difference between the two lines, the true LOF phenotype of the ncORF should be assessed independently of the refORF. |

"LOF" stands for "loss-of-function".

### How to discern specific roles of a refORF and a ncORF in a target gene?

Functional analysis of ncORFs is a challenge regardless of the location of a ncORF in a transcript. Let us begin with a situation illustrated in the third column of Figure 4, which exemplifies nested ncORFs. Insertions and deletions affect both a ncORF and a refORF in the region where the two ORFs overlap. Due to the degenerate nature of the genetic code, an ORF in one frame can be disabled independently from an overlapping ORF through a single-nucleotide change. Such a substitution can be non-synonymous for the target ORF but synonymous for the overlapping ORF. However, there is currently no tool capable of targeted mutagenesis of that type *in vivo*. CRISPR/Cas9-based approaches are still far from such high precision (Doudna & Charpentier, 2014; Guo et al., 2023). However, *in vitro* mutagenesis can help circumvent this limitation. We propose the following experimental setup for studying nested ncORFs. In the first step, both the refORF and the ncORF should be knocked out by a method other than PTGS (El-Sappah et al., 2021) in a stable transformant. This can be achieved by an insertion or deletion that disrupts both ORFs. In the second step, the stable knock-out line should be transformed independently with four constructs: (1) refORF+/ncORF-, which carries a missense substitution that affects the ncORF only; (2) refORF-/ncORF+, which carries a missense substitution that affects the refORF only; (3) refORF+/ncORF+, which carries the wild-type version of the gene; (4) refORF-/ncORF-, which is the genomic full-knockout version. Stable transformants complemented with the first two constructs should show the true loss-of-function phenotypes of the ncORF and the refORF, respectively. Here are examples of some realistic scenarios inferred from the differences between the phenotypes of the initial stable transformant (the double knock-out mutant) and the first two complementation lines. If all three phenotypes are altered but identical to each other, the ORFs may play redundant roles or be part of the same biochemical or regulatory pathway. If each of the three lines has a unique altered phenotype, each ORF has a unique role. Complementation with the wild-type construct (construct 3) should indicate if the mutant background is dominant negative. Failure to restore the wild-type phenotype using the third construct would indicate such a dominant negative effect. However, it cannot tell which of the mutant products acts as a dominant negative. Likewise, this experimental setup is not sufficient to reveal the interaction between the refProt and the ncProt or their mutant forms. Ideally, the knock-out status of each line should be validated with MS proteomics. However, this may be unnecessary if the phenotypes are fully consistent. Transformation with the fourth construct is optional. However, it can help with troubleshooting if the mutant phenotype of the transformant is reverted or heavily modified compared with the stable knock-out line. This can happen when the transformation system alone contributes to the phenotype of the first two complementation constructs.

Now, let us consider ORFs that are either separated on the same transcript (the first and the last columns in Figure 4) or overlap partially (the second and the fourth columns). It may look like demonstrating specific roles of the refORF and the ncORF is a straightforward task when the ncORF is not nested in the refORF. For example, insertional mutants can be established that differentially disable two ORFs in regions of no overlap. However, this simplified view may be very misleading for the following reasons. First, as we will show in the section on *Tnt1* mutagenesis, insertions of diverse types can affect splicing far away from the insertion site. This means that the effects of *Tnt1* and related insertional mutagenesis systems cannot always be limited to a specific position. For example, if an insertion is located in the first of two non-overlapping ORFs, the second ORF may still be affected. For this reason, it may be safer to base functional analysis on CRISPR/Cas9 to avoid off-target effects of large insertions. If *Tnt1* and related systems are used, the whole spectrum of mutant transcripts should be sequenced and carefully analyzed for off-target effects in such studies.

Regardless of the relative position of the ncORF, it should be remembered that off-target effects may not be limited to insertional mutagenesis. A mutation neutral to both the refORF and the ncORF can still affect the abundance of their respective translation products (refProts and ncProts) if it coincides with a regulatory element. Such elements are not limited to the promoter region. They can be found in UTRs, introns, and even coding exons (Mittanck et al., 1997; Neznanov et al., 1997; Scohy et al., 2000; Guo et al., 2006; Moabbi et al., 2012; Birnbaum et al., 2012; Ahituv, 2016; Laxa, 2017; Fiszbein et al., 2019). Thus, validating the effect of any mutation on transcription and translation of the refORF and the ncORF is important to avoid misinterpretation of loss-of-function studies. For example, a substitution located in the 5'-UTR between two non-overlapping ORFs may knock-down transcription of both units without affecting their amino acid sequences. Alternatively, it can selectively affect translation of the downstream ORF by changing the ribosome entry site. These considerations call for assumption-free experimental designs where all possibilities are tested for the maximal accuracy of conclusions. They also emphasize that the potential ambiguity types may be more diverse than those outlined in Figure 4.

#### *Tnt1* insertional mutagenesis: the double-faced god of functional genomics in *M. truncatula*

As we mentioned earlier, no other mutagenesis system in *M. truncatula* has been as popular as *Tnt1* (Figure S1). This unique status is due to the establishment of a large *Tnt1* mutant population at the dawn of *M. truncatula* genomics (Tadege et al., 2008; Lee et al., 2018; Kaur et al., 2021). More than half of the 673 genes listed in Data S6 have been studied with *Tnt1* so far. Despite the popularity and informativity of this crucial resource, surprisingly little attention was paid to its off-target effects other than the mutant background caused by multiple insertions in non-target genes. Because this topic has not been comprehensively discussed in any *M. truncatula* study so far, it is time to summarize the facts that should alert not only the community of *Tnt1* users in *M. truncatula*, but also the broader audience that uses insertional mutagenesis of other types in other systems.

Kryvoruchko et al. (2016) have been among the first to document and realize the importance of the *Tnt1* effects on splicing. However, such effects were soon reported also by other groups. Among the 673 genes studied by loss-of-function approaches in *M. truncatula*, 118 genes have at least one conserved or translated ncORF (Data S12). 65 of these genes were targeted by *Tnt1* alone or in combination with a different system. The effects of *Tnt1* on splicing were reported in five of these studies, which corresponds to approximately 7% (*MtSWEET11*, Kryvoruchko et al., 2016; *MtLAX2*, Roy et al., 2017; *MtSWEET1b*, An et al., 2019; *MtREV1*, Zhou et al., 2019; *MtYSL3*, Castro-Rodríguez et al., 2020). This means that out of 359 genes targeted by *Tnt1* so far, similar effects could be expected in approximately 25 cases. This is not a small number considering the steadily increasing popularity of the resource. Below, we provide brief details of each report and a broader discussion of results of Kryvoruchko et al. (2016) that have not been fully revealed in the original publication on *MtSWEET11*. Finally, we will give examples of similar effects observed in other types of insertional mutagenesis, which show that the interference with splicing is a generic feature of such systems.

According to Figures 3a and S4b in Roy et al. (2017), *MtLAX2* has eight exons. Both *Tnt1* lines, NF14494 (*mtlax2-1*) and NF16662 (*mtlax2-2*), carry the *Tnt1* insertion in Exon 4 (Figure 3a in that study). Instead of causing a truncation of the transcript or its degradation, *Tnt1* acts as an alternative exon so that the end of Exon 3 is spliced to the 3'-portion of *Tnt1*. In these mutant transcripts, 231 or 232 nt of *Tnt1* become integrated into the mRNA while Exon 4 upstream of the insertion site is spliced out. The splicing of the remaining exons is preserved (Figures S4b in Roy et al., 2017). However, the insertion coincides with the nested ncORF (ncORF138), which would result in Type 2 ambiguity (Column 3 in Figure 4), if the ncORF were translated. In this example, the effect on splicing is in cis because it deleted only the upstream portion of the affected Exon 4, and no other exons were affected. However, with regard to any ORF completely or partially located in the deleted segment of Exon 4, the effect is in trans (a sequence beyond the immediate insertion site is affected). The preserved segment of *Tnt1* prematurely truncated the long ORF but did not affect shorter ORFs upstream and downstream of the insertion site. If one of such ORFs were translated in a similar scenario, the refORF would be affected independently of upstream and downstream ncORFs (Type 5 ambiguity, Columns 1, 2, 4 and 5 in Figure 4).

The next example is *MtSWEET1b*, which has six exons (An et al., 2019). The first mutant line considered in the study, NF1309 (*sweet1b-1*), carries the *Tnt1* insertion in Exon 3 (Figure 5 in An et al., 2019), which caused almost complete skipping of the affected exon (Figure S8a in the same study). In the second line, NF3539 (*sweet1b-2*), the *Tnt1* insertion is in Exon 4 (Figure 5 in An et al., 2019). It caused complete skipping of Exon 4 downstream of the insertion site and partial skipping of Exon 4 upstream (Figure S8a in the same study). With regard to ncORF117, which is co-affected by the first insertion, this case would represent Type 3 ambiguity (Figure 4 Column 2), if the ncORF were found to be translated. Like in the case of *MtLAX2*, the effect of *Tnt1* on exons is in cis. However, the effect on potential ncORFs is in trans because regions upstream and downstream of the immediate insertion sites are affected.

The third example is *MtREV1*, which has 18 exons (Zhou et al., 2019). The *Tnt1* mutant line *ppf1-1* used in that study carries the insertion in Exon 6 (Figure 2a in Zhou et al., 2019). Two aberrant splicing variants were recovered from this mutant line (Figure S3a). The shorter form lacked the entire Exon 6. In the longer form, 11 nt of Exon 6 were replaced with 16 nt of *Tnt1*, which caused a frameshift and premature termination of the refORF translation. This example illustrates the same mode of *Tnt1* action on splicing as in *MtLAX2* and *MtSWEET1b* mutants but with one new feature: the incomplete removal of *Tnt1* without affecting the splicing in the longer transcript variant. Because neither ncORF2 (MS-validated) nor ncORF3 (conserved) were affected by these transcript rearrangements, this example falls into Type 5 ambiguity with regard to ncORF2 (Figure 4 Column 2).

The fourth example is *MtYSL3*, with seven coding exons and one non-coding exon the 5'-UTR (Castro-Rodríguez et al., 2020). One of the *Tnt1* lines used in that study, NF12068 (*ysl3-2*), carried the insertion in the promoter region. In the other line, NF17945 (*ysl3-1*), *Tnt1* was found at the end of the first coding exon (Exon 1 in Figure 3a, Castro-Rodríguez et al., 2020). In the mutant transcript of that line, *Tnt1* caused skipping the end of Exon 1 immediately downstream of the insertion site. This segment was substituted with 30 nt from the *Tnt1* sequence, which altered amino acids YSIAVG at that position but preserved the rest of the sequence (Figure S7a, Castro-Rodríguez et al., 2020). In this example, we see the same mode of *Tnt1* action on splicing as described above. NcORF118 is unaffected by this transcript rearrangement. If it were translated, it would cause Type 5 ambiguity (Figure 4 Column 4).

In studies on *MtLAX2*, *MtSWEET1b*, *MtREV1*, and *MtYSL3*, only cis effects on splicing were described. Specifically, the effects were limited to complete or partial skipping of the exon carrying the *Tnt1* insertion, with or without partial retention of the *Tnt1* sequence. In contrast, the study on *MtSWEET11* (Kryvoruchko et al., 2016) revealed a much broader range of effects on splicing, which included skipping exons and retention of introns far from the insertion site. Below, we describe the authors' motivation for the deeper look into the transcript fate of *MtSWEET11* and discuss the original data in full.

*MtSWEET11* was a gene with very high expectations as to its symbiotic phenotype. These expectations were based on the very strong and specific expression of this gene in rhizobia-populated root nodules and its transport specificity to sucrose (Kryvoruchko et al., 2016). Therefore, it was surprising to find that two independent *Tnt* mutant lines, NF12718 (*sweet11-1*) and NF17758 (*sweet11-2*), both affected in Exon 3 of *MtSWEET11*, showed absolutely no alteration in the nodulation phenotype, even under reduced illumination. The publication proposed an explanation based on functional redundancy. However, another possibility has not been discussed. Is it possible that the mutant transcripts retained partial or complete functionality? Naturally, the transcription of this gene in the mutants was checked with qRT-PCR. The cDNA-specific qRT-PCR amplicon was designed in Exons 5 and 6, far downstream of the insertion site. Nearly half of the wild-type expression signal was still detectable in the mutant lines. Then, regular PCR was conducted on cDNA prepared from mutant nodules, for each mutant line independently. In this PCR, primers were the same as in the cloning reaction amplifying the entire CDS. Products of this reaction were visualized on agarose gel. They appeared as a faint smear with a few accentuated bands ranging approximately between 600 and 6,000 bp (Supplemental Figure S10 in Kryvoruchko et al., 2016). The sizes of these bands corresponded to two predominant forms: one with the entire *Tnt1* retained in the CDS (6,147 bp) and the other with Exon 3 fully skipped (599 bp). The lengths of *Tnt1* and the CDS are 5,334 bp and 813 bp, respectively. Next, the gel was sliced and PCR products were purified from five different regions between and including the shortest and the longest band, separately for each mutant line. Then, these products were ligated into a cloning vector, and bacterial colonies were isolated for sequencing of the insert (20 colonies for line NF12718 and 12 for line NF17758). In addition to two major aberrant splicing forms mentioned above, more exotic splicing variants were recovered (Figure S6 of the present study). They can be classified into 11 categories. In two most frequently found forms, either Exon 3 alone or Exon 3 together with Exon 2 were skipped by the splicing machinery, which is very similar to the situation observed in other reported cases (see above). However, the authors also observed splicing forms in which distant exons were skipped (Types VIII, IX and X) or distant introns retained (Types III, IV, and VII), partially or completely. *Tnt1* was retained partially in some forms (Types VI, IX, X, and XI) and completely in Type VIII. Exon 3 nucleotides were copied from the opposite side of the insertion and reinserted next to the right border of *Tnt1* in Types XIII and XI. These findings indicate that the effects of *Tnt1* on splicing are by far more diverse than was believed before.

This data is relevant to the interpretation of the mutant phenotype of *MtSWEET11*. One aberrant splicing form (Type X) can potentially be translated into a relatively long protein of 223 aa, with most of the sequence unchanged compared to the 270-aa-long wild-type version (Figures S7 and S8 of the present study). It remains to be demonstrated if that form can rescue the null mutants of *MtSWEET11*. The absence of this form from line NF12718 suggests that such a possibility is unlikely. It should be noted that the experimental setup of Kryvoruchko et al. (2016) revealed only those aberrant transcript versions in which primer annealing sites were preserved. Thus, many other splicing forms could remain undetected. RNA-Seq is required to provide comprehensive data on all aberrant splicing forms in a *Tnt1* mutant line. By 2025, numerous *Tnt1* mutant lines were processed by RNA-Seq, which makes the large-scale analysis of aberrant splicing forms straightforward (for example, Wang et al., 2016; Zinsmeister et al., 2020; Cheng et al., 2021; Lalanne et al., 2021; Irving et al., 2022; Bai et al., 2022; Jaudal et al., 2022; Poulet et al., 2024). Unfortunately, none of these lines have corresponding MS proteomic data yet, which identifies the area where new data are wanted for the holistic view on this phenomenon. It would be very useful to learn which aberrant splicing forms detected in lines NF12718 (*sweet11-1*) and NF17758 (*sweet11-2*) are translated.

We should emphasize that these five examples (*MtLAX2*, *MtSWEET1b*, *MtREV1*, *MtYSL3*, and *MtSWEET11*) do not present a complete list of studies where the effect of *Tnt1* on splicing was documented. We did not check the structure of mutant transcripts in genes with no translated or conserved ncORFs. Conceivably, many more examples can be found upon closer examination of those studies. However, transcript structure is usually not analyzed by sequencing when *Tnt1* mutagenesis is used, which makes it difficult to assess the true frequency of transcript rearrangements. Thus, a dedicated large-scale study addressing this question is warranted.

#### Interference with splicing is a generic feature of insertional mutagenesis not limited to *Tnt1*

The effects of transposons on splicing have been known since 1990s (Ortiz and Strommer, 1990; Menssen et al., 1990; Varagona et al., 1992; Giroux et al., 1994). Since then, numerous studies have reported such effects in plants (e.g. Roquis et al., 2021; Berthelier et al., 2023) and animals (e.g. Lev-Maor et al., 2003; Chen et al., 2006; Rodríguez-Martín et al., 2016). This topic has recently been comprehensively reviewed (Pfaff et al., 2022; Emmerson and Catoni, 2025). It was also shown that T-DNA insertions cause similar transcript rearrangements in plants (Babiychuk et al., 1997; Smalle et al., 2002; Wang et al., 2003; Wang, 2008; Missihoun et al., 2012; Thulasi Devendrakumar et al., 2024). Inserted elements supply cryptic splice donors/acceptors or enhancers/silencers (Lev-Maor et al., 2003; Sorek et al., 2004; Lei and Vorechovsky, 2005; Pastor et al., 2009; Payer et al., 2019). They can also form dsRNA (e.g., inverted *Alu* pairs) that remodels the splice choice of nearby exons (Lee et al., 2024).

#### Relevance of insertion-induced aberrant splicing for functional genomics and biosafety

Natural transposition shapes the evolution of genomes through various processes. Besides numerous regulatory functions, transposons create novel proteins that are tested for fitness in specific biological contexts (Bourque et al., 2018; Nicolau et al., 2021; Betancourt et al., 2024; Almeida et al., 2025). It has been demonstrated that transposon effects similar to those we described in *M. truncatula* create aberrant protein isoforms subjected to natural selection due to the acquisition of novel functions (Arribas et al., 2024). In this section, we highlight the need to apply this knowledge in studies that involve induced insertional mutagenesis.

Let us begin with trans effects of insertions on ncORFs. As we discussed above, *Tnt1* and other types of insertional agents can affect segments of mature mRNA far away from an insertion site (trans effects on splicing). Unfortunately, not only such trans effects but also ncORFs are typically left without consideration in functional studies. When these two important phenomena are overlooked, the correct interpretation of loss-of-function phenotypes can be severely compromised. An insertion in a refORF can co-affect a distantly located ncORF. If such a ncORF is translated and crucial for a specific biological process, its loss-of-function phenotype is traditionally attributed to the refORF, even if the refORF has no crucial function. Conversely, if an insertion is in a UTR, and it causes a clear altered phenotype, it may have a triple effect: (1) downregulation of the entire transcript due to disruption of a regulatory element; (2) disruption of the refORF due to a trans effect on splicing; (3) co-disruption of a nested translated ncORF by the same trans effect. However, in such a scenario, the phenotype is conventionally attributed to the first effect alone (downregulation). Hence, the mutant is typically considered as a knockdown line, and no sequencing of the transcript is conducted, based on our in-depth analysis of the available literature. These two simple scenarios should be sufficient to show how much false information can be generated, and have likely been generated so far, with ncORFs and trans effects on splicing left behind the scenes.

Secondly, the potential effects of insertion-induced aberrant splicing on refORFs should be discussed, which is important even in the absence of ncORFs (Figure S9). Since insertions reorganize transcripts rather than knocking them out, newly created nucleotide sequences may be translated and functional (Arribas et al., 2024). In a natural environment, with millions of years available for evolutionary trials, sequences that have detrimental effects on the organism are eliminated. However, in an artificial population of mutants such as the *Tnt1* mutant resource, such harmful proteins may mimic loss-of-function phenotypes, while their true effects are due to the gain of function (for example, toxicity or suppression). If the wild-type version of the target protein has no crucial role, the toxicity effect of an aberrant proteoform may be misinterpreted as the loss of a crucial function of the refProt. Conversely, if the product of aberrant splicing acquires a positive function, it has the potential complement the true loss-of-function phenotype of a biologically important refProt (Figure S9).

It is conceivable that, in rare cases, a protein product of aberrant splicing may be highly toxic to humans. Given the large number of background insertions in each mutant line and the very large number of lines necessary for nearly saturating mutagenesis (ca. 20,000 for the *Tnt1* resource), the chance of such an unlikely event increases. Most importantly, the outcomes of induced missplicing are unpredictable because they are sequence- and position-dependent (thus, random). This consideration should raise awareness about biosafety concerns associated with any large insertional mutant population. After all, it may be wise to wear gloves when handling seemingly harmless mutant plants in a greenhouse. These unexpected effects of insertional mutagenesis on the phenotype can be compared with the underwater part of an iceberg: when overlooked, they may lead to unwanted outcomes (Figure S9).

It may be argued that biological relevance of the aberrant proteoforms discussed above should be minor because most misspliced transcripts reported in *M. truncatula* are expressed at very low levels. This, however, is not the case with *MtSWEET11*. Transcription of its mutant isoforms was very high (nearly half of the wild-type level), as detected with qRT-PCR primers specific to *MtSWEET11* (Kryvoruchko et al., 2016). This signal could not come from any contamination with genomic DNA because the forward qRT-PCR primer was designed at the border between Exon 5 and Exon 6, which exists as a continuous sequence only in the spliced mRNA of *MtSWEET11*. Even if the abundance of an aberrant splicing form is low, such a transcript can serve as a substrate for ribosomes and produce translation products.

Translation of low-abundance transcripts can be very efficient, as was shown and discussed in numerous studies, including the emerging topic of specialized ribosomes (Gygi et al., 1999; Griffin et al., 2002; Dinman, 2016; Emmott et al., 2019; Ferretti and Karbstein, 2019; Mair et al., 2020; López García de Lomana et al., 2020; Lipo et al., 2022; Reyna-Llorens et al., 2023). However, even if expressed and translated at a low rate, a protein can have a profound biological effect (Forde and McCutchen-Maloney, 2002; Ghaemmaghami et al., 2003; Gregor et al., 2007; Ding et al., 2013; Brumbaugh et al., 2014; Garza de Leon et al., 2017; Boschetti and Righetti, 2023), which further supports the need to consider aberrant proteoforms in functional studies.
