## Supporting Figures for "Loss-of-function phenomics, ncORFs, and ambiguity of mutant phenotypes in *Medicago truncatula*"

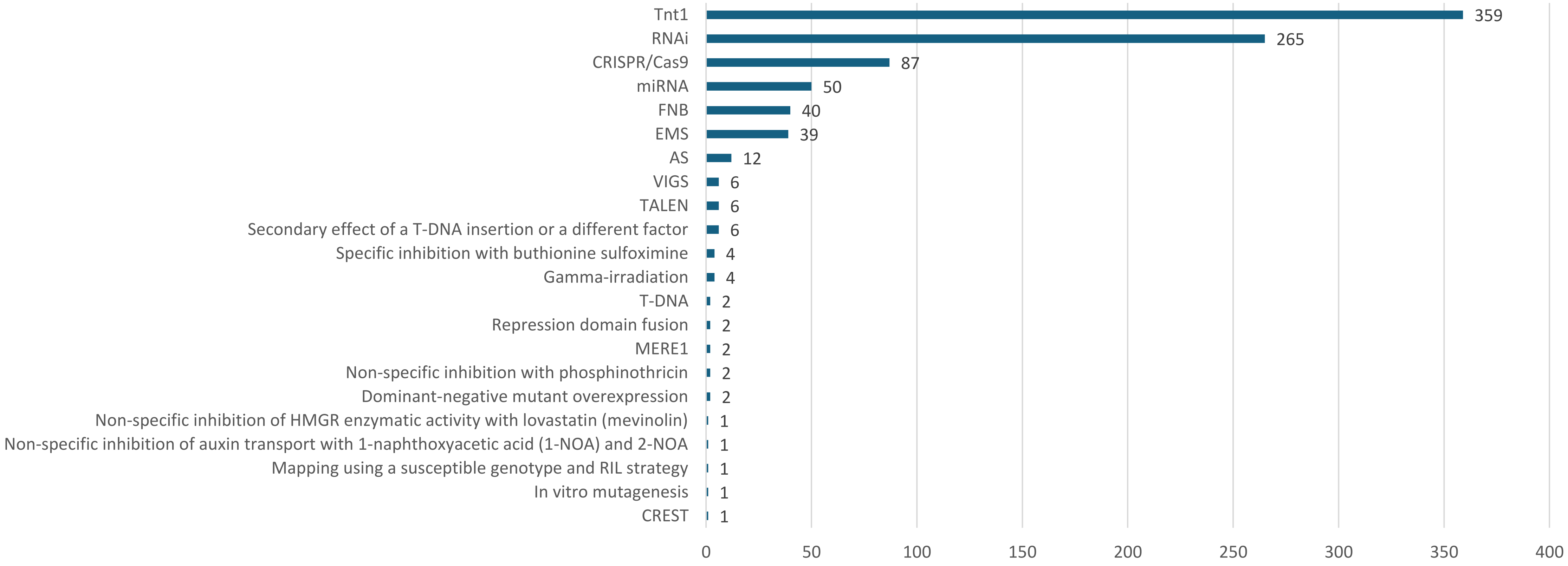


**Figure S1**. Loss-of-function approaches used for studying 673 genes in *Medicago truncatula*. The counts are not additive because many genes were studied by more than one approach. Acronyms are explained in Data S1-S4.

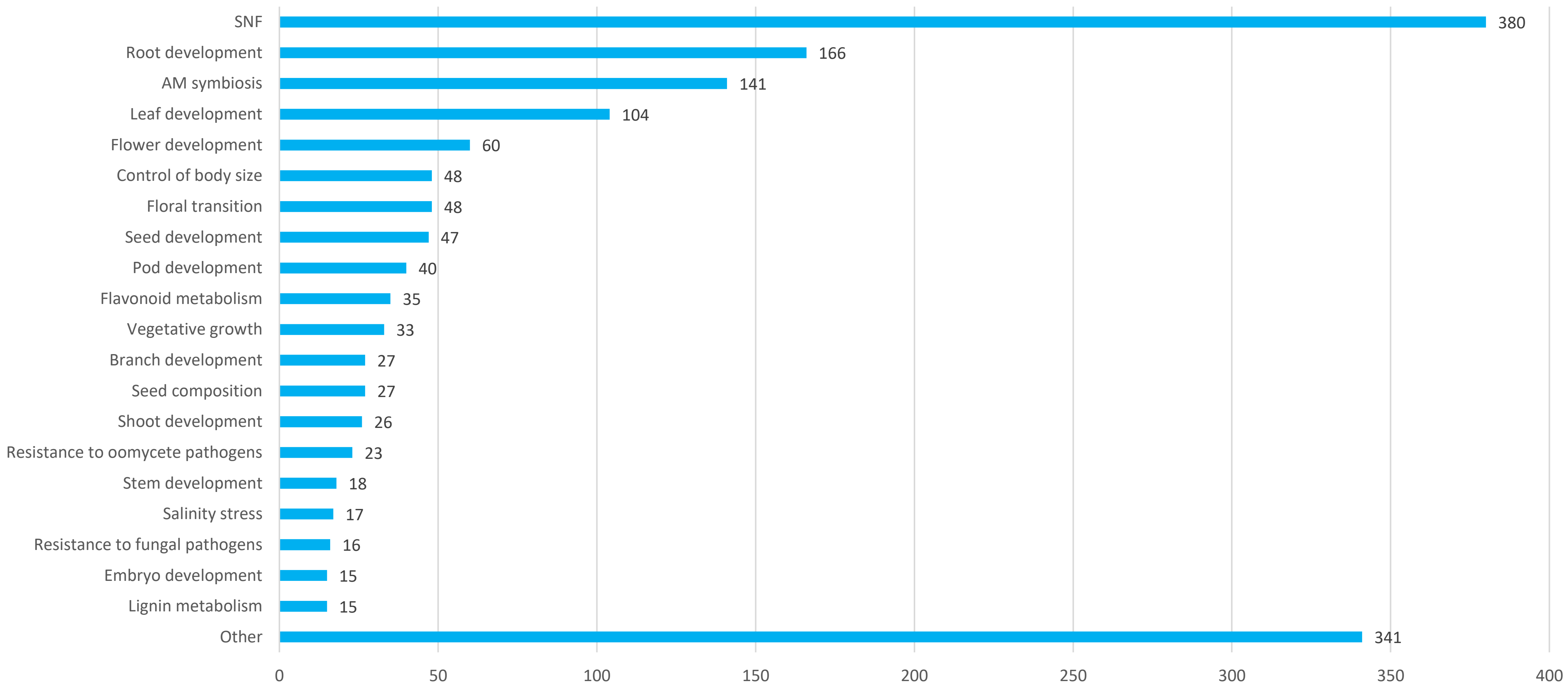


**Figure S2**. Biological processes examined in loss-of-function studies on 673 genes in *Medicago truncatula*. The counts are not additive because many genes were tested for more than one process. The complete list of biological processes is available in Data S9.

| Total count | A, count | C, count | N, count | A, proportion | C, proportion | N, proportion | Group |
|---|---|---|---|---|---|---|---|
| 224 | 119 | 77 | 28 | 53 | 34 | 13 | TF |
| 205 | 135 | 57 | 13 | 66 | 28 | 6 | Enzyme |
| 99 | 72 | 12 | 15 | 73 | 12 | 15 | Other |
| 73 | 52 | 13 | 8 | 71 | 18 | 11 | Transporter |
| 30 | 17 | 10 | 3 | 57 | 33 | 10 | Peptide |
| 22 | 11 | 9 | 2 | 50 | 41 | 9 | Membrane protein |
| 13 | 8 | | 5 | 62 | 0 | 38 | Channel |
| 7 | 5 | 1 | 1 | 71 | 14 | 14 | Unknown |
| 673 | 419 | 179 | 75 | 62 | 27 | 11 | All |

**Figure S3**. Different groups of protein classes have different proportions of phenotypes, which may reflect their relative functional importance. Letters A, C, and N stand for altered, conditional, and neutral phenotypes, respectively. Statistical analysis of the proportions is available in Data S11.

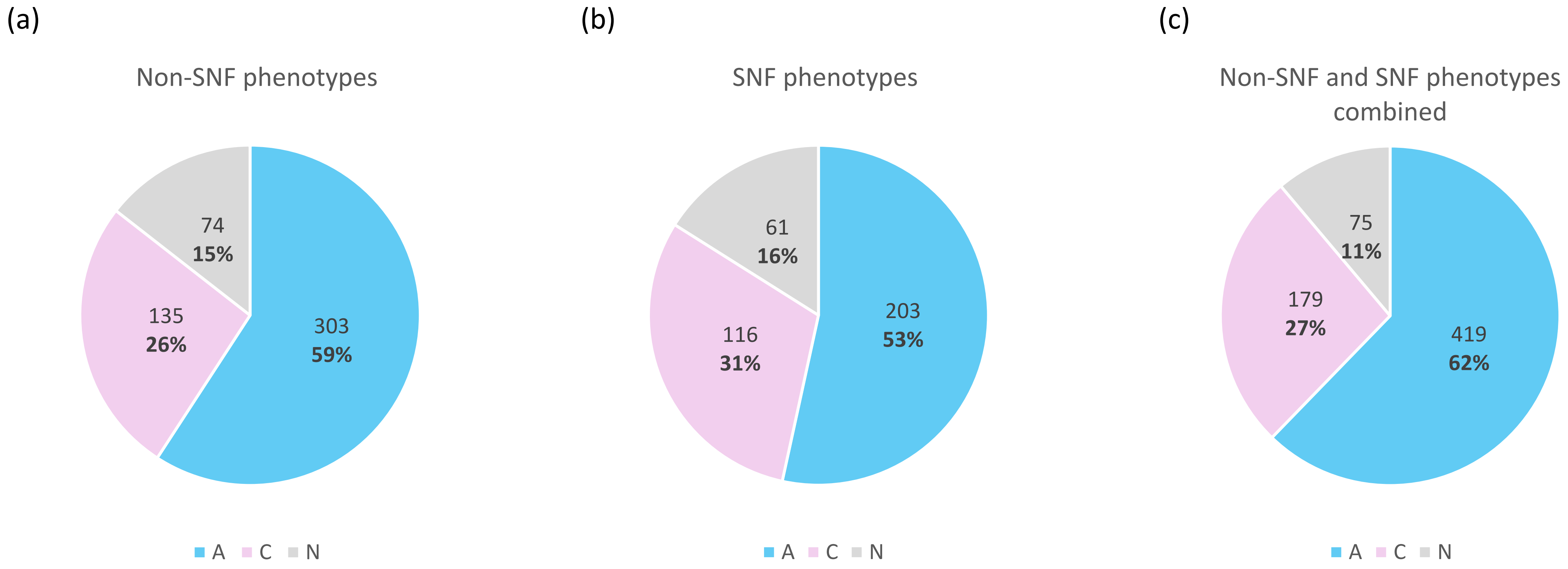


**Figure S4**. Partitioning of phenomics data into altered (A), conditional (C), and neutral (N) phenotypes. A phenotype is categorized as altered (A) if there is at least one phenotypic difference from the wild type that is not conditional. In this context, "conditional" refers only to genetic conditions, where there is ambiguity about the causative gene (e.g., a double mutant). It does not refer to non-genetic conditions such as specific growth conditions or mutant lines. A phenotype is categorized as conditional (C) if there is no "A"-phenotype and there is at least one phenotypic difference from the wild type that is genetically conditional. A phenotype is categorized as neutral (N) if there are no "A"- and "C"-phenotypes. SNF stands for symbiotic nitrogen fixation.

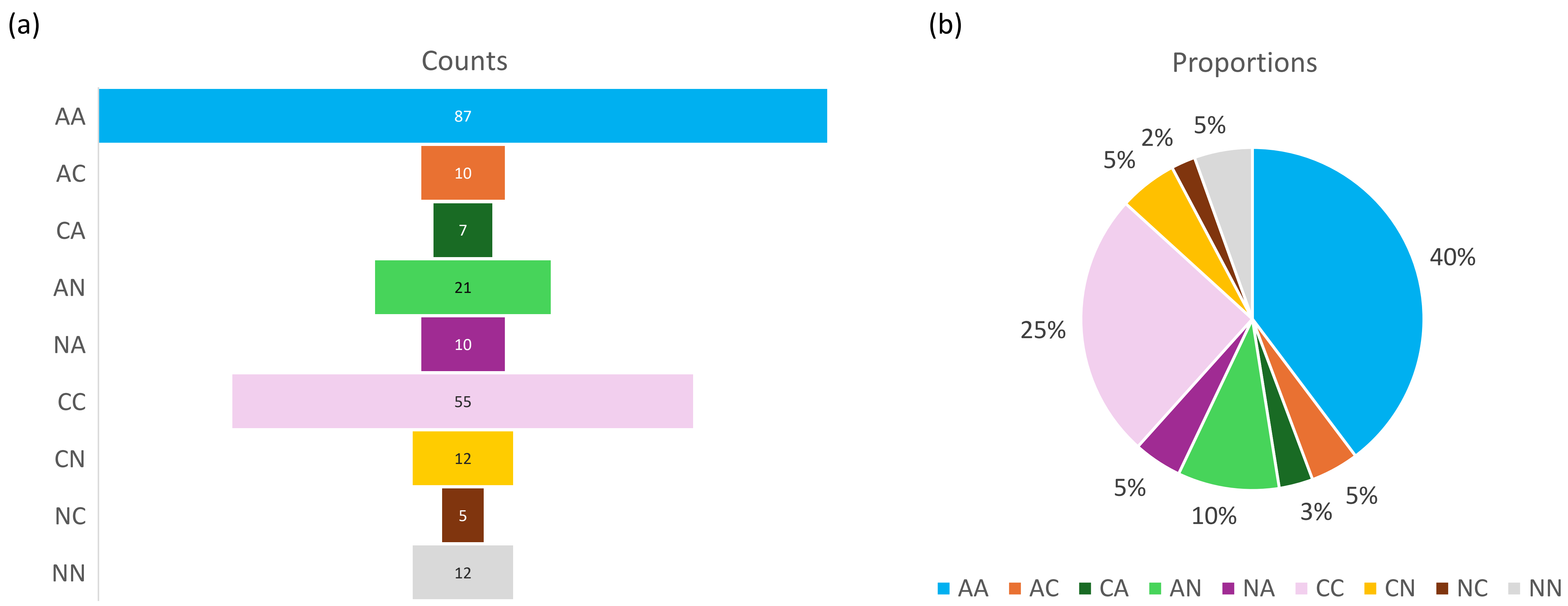


**Figure S5**. Phenotypes of 219 genes studied in the context of both symbiotic nitrogen fixation (SNF) and other processes. The first letter corresponds to a non-SNF phenotype, and the second letter corresponds to an SNF phenotype of the same gene. Letters A, C, and N stand for altered, conditional, and neutral phenotypes, respectively. (a) Counts of genes in corresponding phenotypic categories. (b) Proportions of genes in the same phenotypic categories.

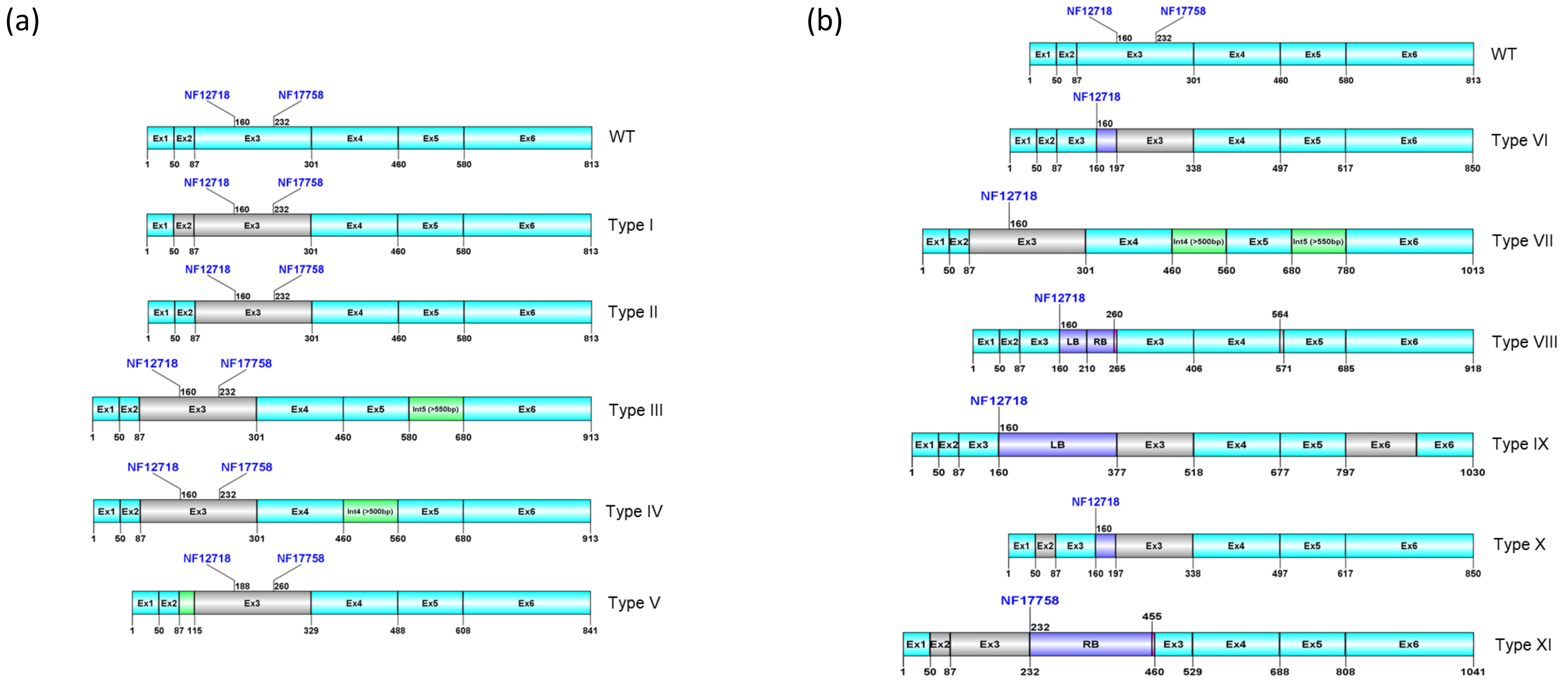


**Figure S6**. Aberrant transcript forms recovered from two *Tnt1* mutant lines of *MtSWEET11* (Kryvoruchko et al., 2016). The *Tnt1* insertions are located in Exon 3 in two different mutant lineages. Different types are shown in the descending order of their frequencies. (a) Five types of aberrant transcripts common to both mutant alleles, NF12718 and NF17758. (b) Six types of aberrant transcripts specific to either mutant allele. WT: exons in the wild-type version of the *MtSWEET11* coding sequence. Cyan boxes: exons. Gray boxes: skipped exons. Green boxes: retained introns. Violet boxes: *Tnt1* sequences. Magenta boxes: Exon 3 nucleotides copied from the opposite side of the insertion (Types XIII and XI). LB: left border of *Tnt1*. RB: right border of *Tnt1*. The entire *Tnt1* sequence is present between LB and RB in Type VIII but is not shown because of space limitations.

| Type | Frequency | | Mutant protein length*, aa |
|---|---|---|---|
| | NF12718, out of 20 clones | NF17758, out of 12 clones | |
| I | 6 | 5 | 44 |
| II | 2 | 2 | 37 |
| III | 1 | 2 | 37 |
| IV | 1 | 1 | 37 |
| V | 1 | 1 | 29 |
| VI | 2+2 | 0 | 56 |
| VII | 2 | 0 | 37 |
| VIII | 1 | 0 | 56 |
| IX | 1 | 0 | 56 |
| X | 1 | 0 | 223 (!) |
| XI | 0 | 1 | 25 |

**Figure S7**. Frequencies of aberrant transcript forms recovered from two *Tnt1* mutant lines shown in Figure S6. The table also indicates the lengths of mutant versions of MtSWEET11. Translation of most splicing forms is truncated early due to an in-frame stop codon. However, Type X splicing forms can potentially result in a relatively long protein shown in Figure S8. The length of the wild-type version of MtSWEET11 is 270 amino acids. Type VI has two variants. They differ by the length of the partially preserved sequence of *Tnt1*: the first 37 bp (Type VIa) or the first 74 bp (Type VIb) of *Tnt1* are preserved in these variants. They were equally represented among 20 clones from line NF12718 (thus, “2+2” in the corresponding row).

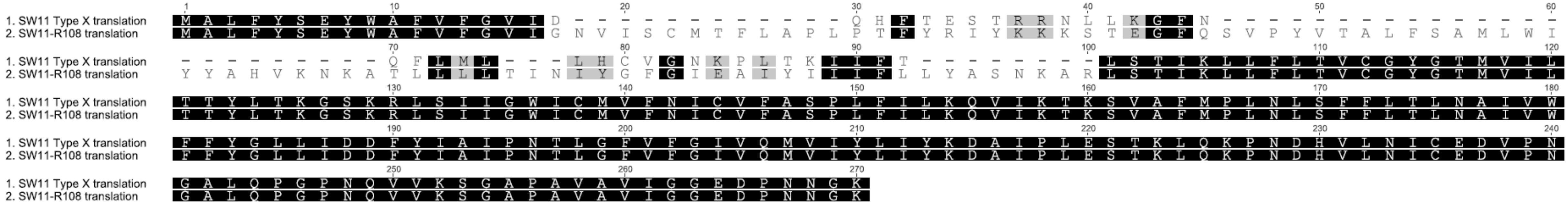

**Figure S8**. A ClustalW protein alignment of the mutant version of MtSWEET11 (Type X) with the wild-type version from ecotype R108 (the background of the *Tnt1* mutant population). Note that most amino acids are preserved in the mutant. Amino acids lost due to exon skipping are substituted with amino acids from the retained but frameshifted portion of Exon 3 and *Tnt1* acting like an alternative exon. The original reading frame is regained at position 101. Regardless of the effect on *MtSWEET11*, short reading frames (potential ncORFs) far from the insertion site are unaffected in this and some other splicing forms but are changed in some forms such as Type III, IV, VII, VIII, and IX, which makes this type of trans effect of *Tnt1* highly relevant for research on ncORFs.

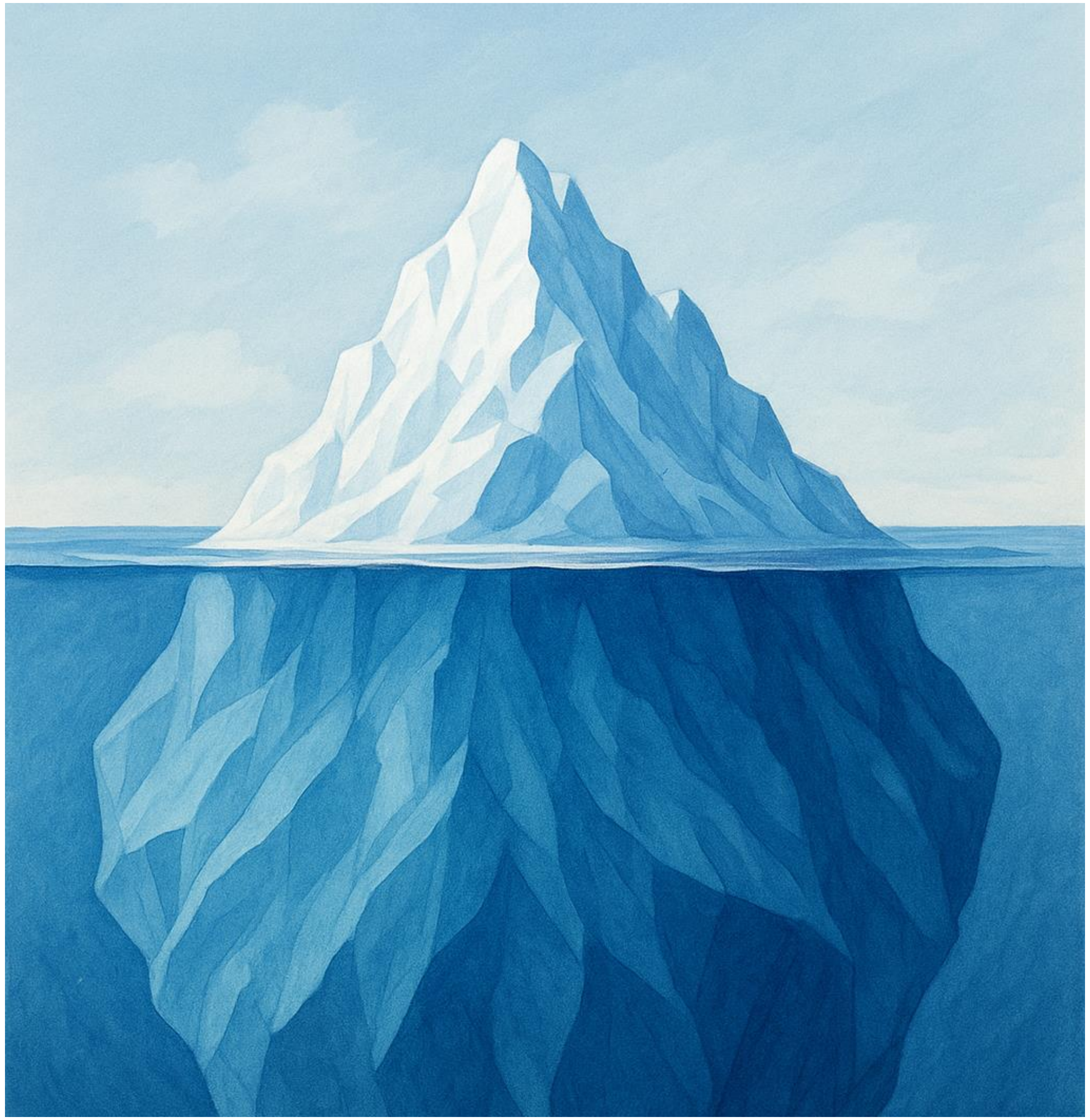

**Expected effects**:

- Affected or induced transcription
- Affected or induced translation
- Translation into a non-functional protein
- Partial functionality of the mutant gene product (same function as in the wild type, but at a lower rate)

**Unxpected effects**:

- Acquisition of a novel function, possibly a toxic effect (false-positive mutant phenotype)
- Acquisition of a novel function that reverts the true mutant phenotype (false-negative result, failure to demonstrate the true phenotype)
- Potential pleiotropic effect of the mutant product on the whole metabolome (biosafety)

**Figure S9**. Biological and biosafety implications of induced insertional mutagenesis. Induced insertional mutagenesis may have expected and unexpected effects on the phenotype of affected organisms. The unexpected effects can be compared with the underwater part of an iceberg. The figure lists some but not all possible scenarios.
