## Supporting Methods for "Loss-of-function phenomics, ncORFs, and ambiguity of mutant phenotypes in *Medicago truncatula*"

### Sequence similarity analyses of ncORFs and ncProts

Taxonomic ranges of ncProts were deduced from source species of all BLASTP hits using DIAMOND v. 0.9.14 as a search software (Buchfink et al., 2021) and UniProt v. 2020_02 as a reference database (UniProt Consortium, 2019). DIAMOND-based BLASTP hit numbers and taxonomic ranges listed in Data S12 were subsequently confirmed or edited based on the latest release of UniProt (Reference_Proteomes_2025_04, updated on 15 October 2025). BLASTN hit numbers listed in Data S12 were confirmed or edited using the latest release of the NCBI core_nt database (12 December 2025). Taxonomic information missing in UniProt was retrieved from the publicly accessible web resources. Taxonomic ranges were deduced manually from higher taxonomic ranks of source species. Each range was expressed as one of the following terms or a combination of these terms: "Species", "Genus", Legumes, Eudicots, Monocots, Flowering plants, Seed plants, Vascular plants, Bryophytes, Charophyta (Chara-like algae), Green algae, Red algae, Other protists, Fungi, Animals, Archaea, Eubacteria, and Viruses. "Species" and "Genus" terms represent corresponding species and genus names, if the taxonomic range is limited to only one species or members of a single genus, respectively. The taxa are listed in the ascending order of their evolutionary distances from *M. truncatula*. The term Eudicots refers to Core eudicots in our analysis. Basal eudicots and Magnoliids were ranked as Flowering plants. For example, if a ncProt has many significant hits among Core eudicots but also a single hit corresponding to a Basal eudicot, with no hits in other organisms, the taxonomic range is reported as Flowering plants. Charophyta (Chara-like algae), Green algae, and Red algae were separated from Other protists because they are the closest eukaryotic relatives of plants (One Thousand Plant Transcriptomes Initiative, 2019). Viruses were shown at the end of the list, although the latest evidence suggests that they may be related to Archaea (Harada et al. 2025).

To find ncORFs more conserved at the amino acid level compared to the nucleotide level, DIAMOND-based BLASTP hit numbers were divided by BLASTN hit numbers for each ncORF. The NCBI BLASTN tool (Altschul et al., 1990; Boratyn et al., 2013) was used with the non-redundant nucleotide collection as a search database. The focus was placed on mRNA hits (non-mRNA hits were filtered out). NcORFs with large ratios of BLASTP/BLASTN hits were subjected to manual searches for evidence of purifying selection. Specifically, synonymous substitutions were visualized in MUSCLE nucleotide alignments of ncORFs with subject sequences using Geneious® v. 7.1 (Dotmatics Ltd., MA, USA, https://www.geneious.com). Percentages of amino acid and nucleotide identities were calculated from corresponding alignments. NcORFs with larger percentages of amino acid identity compared to nucleotide identity were subjected to the analysis of dN/dS ratios (Hurst, 2002) using the MEGA-X software v. 10.2.6 (Kumar et al., 2008). A built-in Z-test option (with subsequent manually calculated Bonferroni correction for multiple comparisons) was used to estimate the statistical significance of dN/dS ratios. It should be noted that these tools were developed for the analysis of long sequences. The actual statistical significance of dN/dS ratios cannot be accurately computed for short sequences (<300 codons) such as many ncORFs (Zhang et al., 2006; Zhao et al., 2024). The detection of conservation signatures in characterized NCR peptides MtNCR086, MtNCR211, and MtNCR314 (Data S12) using our approach validates its efficiency in finding biologically relevant ncORFs.

### Phylogenetic analysis of genes harboring super-conserved ncORFs

To visualize the relationships between ncORF source sequences and their homologs, MUSCLE alignments were prepared in Geneious® v. 7.1 (Dotmatics Ltd., MA, USA, https://www.geneious.com). Because in a few cases MUSCLE alignments produced suboptimal results, some sequences were aligned using Clustal Omega (Sievers et al., 2011) and imported into Geneious® for visualization. The type of alignment is specified in Data S17, in which all the alignments are presented. Phylogenetic trees were built with MEGA-X v. 10.2.6 (Maximum Likelihood method) or Geneious® v. 7.1 (UPGMA method).

**Cloning and sequencing of aberrant splicing forms of *MtSWEET11***

The procedure was briefly described in the original article on *MtSWEET11* (Kryvoruchko et al., 2016). RNA from two *Tnt1* lines, NF12718 (*sweet11-1*) and NF17758 (*sweet11-2*), was converted to cDNA, followed by regular PCR with primers used for cloning the CDS of this gene. The PCR products were visualized on agarose gels. Five segments of the gel were sliced, separately for each *Tnt1* line, and PCR products were purified using Promega Wizard® SV Gel and PCR Clean-Up System (Promega, WI, USA). This step was necessary to minimize the ratio of agarose gel volume to the DNA content, which maximizes the DNA yield. The purified products were pooled for each *Tnt1* line and cloned into the Gateway® entry vector pDONR207 (Invitrogen, MA, USA) for sequencing. For lines NF12718 and NF17758, 20 and 12 bacterial colonies were picked up, respectively. Sanger sequencing was conducted in two directions using vector-specific and gene-specific primers indicated in Kryvoruchko et al. (2016). Most of the reads covered the entire length of the CDS (813 bp). The reads were aligned with the genomic DNA and cDNA of *MtSWEET11* from the *M. truncatula* ecotype R108.
