## Supporting Results for "Loss-of-function phenomics, ncORFs, and ambiguity of mutant phenotypes in *Medicago truncatula*"

### NcORFs in *M. truncatula* genes targeted by loss-of-function approaches

The overall purpose of this study was to improve the accuracy of functional genomics in *M. truncatula* in three ways: (1) comprehensive inventory of loss-of-function studies and their linkage to locus IDs of the current genome assembly; (2) correction of gene models, elimination of redundancy, and resolution of ambiguities associated with locus names; (3) revision of reported phenotypes, taking into account the potential contribution of ncORFs. This Supporting Results document is focused on ncORFs in *M. truncatula* transcripts annotated in the genome assembly v. 5.1.9. It considers only the loci targeted by functional analyses.

In a narrower sense, this study is a branch of our earlier work dedicated to the identification of translated and conserved ncORFs in the whole transcriptome of *M. truncatula* (Çakır et al., 2023; 2025b). We used a very broad definition of a ncORF as a stop-free region in any forward frame of any mature transcript (mRNA, ncRNA, rRNA, and tRNA) with a length of at least 60 nt, regardless of the presence of translational start codons, ribosomal entry sites, Ribo-Seq data, or any other evidence for translation. Importantly, our definition of a ncORF does not place any upper limit on its length. Thus, ncORFs in our study are not always short (compare with the term "sORF", which stands for "short ORF"). This is somewhat different from the classical definition of ncORFs (Mouilleron et al., 2016; Mudge et al., 2022; Álvarez-Urdiola and Riechmann, 2025) and is much more inclusive. Nevertheless, we adopted the naming convention established for different types of ncORFs by Mudge et al. (2022) (Figure 4, Data S12) as it reflects the distinctive functions, frequencies, and technical challenges with the identification of different ncORFs depending on their location relative to a refORF (Mudge et al., 2022). This classification recognizes five categories of mRNA-ncORFs: uORFs, uoORFs, intORFs, doORFs, and dORFs. It also proposes different names for non-mRNA-ncORFs depending on the transcript type: lncRNA-ORFs, circRNA-ORFs, and even miORFs (Mudge et al., 2022; Álvarez-Urdiola and Riechmann, 2025). Our study adds two more types to the non-mRNA-ncORF category: rRNA-ORFs and tRNA-ORFs (see also Çakır et al., 2025b). It also introduces one mRNA-ncORF type left beyond consideration in previous studies. NcORF97 in Data S12 starts in the 5'-UTR and ends in the 3'-UTR. We propose to term this type "bridge ORF" (bORF).

Using MS proteomic data from three publicly available studies (PXD002692, Marx et al., 2016; PXD013606, Shin et al., 2021; and PXD022278, Castañeda et al., 2021), we identified 805 MS-validated ncProts detected in at least one of 16 samples, 720 of which originated from mRNA transcripts and 85 from ncRNA (Çakır et al., 2025b, Supplementary Figure S6). These 805 sequences were mapped to the current genome and are available as a separate track in the genome browser (https://medicago.toulouse.inra.fr/MtrunA17r5.0-ANR/). At the same time, we investigated the similarity of *in silico* ncORF translations (ncProts) to annotated proteins using the curated global protein collection UniProt v. 2020_02 (UniProt Consortium, 2019). This analysis identified 18,643 *in silico* ncProts that have at least one significant hit in UniProt (e-value ≤ 0.001). Among those sequences, 13,078 had the top hit with percent identity of at least 70 (Çakır et al., 2025b, Supplementary Figure S1). Although many of such ncProts had similarity to *M. truncatula* proteins (Çakır et al., 2025b, Supplementary Figure S7), we conditionally refer to them as conserved. Later in the text, we will explain why this broader definition of evolutionary conservation is valuable for functional studies.

We compared locus identifiers of 673 genes studied by loss-of-function approaches with locus identifiers of transcripts that have translated and conserved ncORFs. This analysis revealed 137 mRNA-ncORFs and three lncRNA-ORFs in 118 loci targeted by loss-of-function approaches (140 ncORFs in Data S12). The number of ncORFs is higher than the number of loci because some genes have more than one ncORF (for example, *MtPHO2-A* has five ncORFs). Additionally, we identified eight ncORFs from Data S2 and S4 targeted by gain-of-function approaches only (five loci), which makes the total of 148 ncORF in Data S12. Among the 118 genes studied by loss-of-function approaches, there are eight loci with MS-validated ncORFs, three loci with ncORFs that were MS-validated and conserved at the same time, and 114 loci with conserved ncORFs. It should be noted that only one of the 11 MS-validated ncORFs in Data S12 has been known so far: ncORF9 in locus MtrunA17_Chr7g0229931. All three MS peptides of this ncORF match the characterized mature peptide MtNCR169 (Starker et al., 2006; Domonkos et al., 2013; Farkas et al., 2014; Horváth et al., 2015, 2023; Güngör et al., 2023; Li et al., 2025). It is different from the refProt of locus MtrunA17_Chr7g0229931. All the other ncORFs in the remaining loci were never considered in the original functional studies.

Next, we organized all 123 loci with 148 ncORFs in three tiers according to the potential biological relevance (Data S12 Column A). This list combines ncORFs studied by loss-of-function and gain-of-function approaches (140 and eight ncORFs, respectively). Tiers were based on loci, not on ncORFs, to enable visualization of ncORFs side-by-side when multiple ncORFs originate from the same locus. Tier 1 is composed of loci with at least one MS-validated ncORF. This tier includes a few conserved ncORFs that have no MS-validation. They were added because corresponding transcripts had at least one MS-validated ncORF. Tier 2 combines loci having at least one conserved ncORF with the top hit of at least 70% identity to any subject in the DIAMOND-based BLASTP search (NCBI/UniProt, e-value at most 0.001). This tier includes a few conserved ncORFs with the top hit below 70% identity (NCBI/UniProt, e-value at most 0.001), which correspond to transcripts with more than one ncORF. Tier 3 is reserved for loci having at least one conserved ncORF with the top hit below 70% identity (NCBI/UniProt, e-value at most 0.001). Tier 1 loci are likely the most biologically relevant, and Tier 3 loci are expected to be the least biologically relevant due to the lack of MS-validation and weak conservation signatures of their ncProts.

To build the master table of ncORFs (Data S12), we imported relevant data from Data S6 and obtained additional information for each of 123 loci from the original publications. The key points of this in-depth literature analysis are shown in columns N-R of Data S12. This includes the effect of loss-of-function tools on ncORFs and refORFs, notable phenotypic differences between mutant lines of the same genes (if multiple lines were available), and comments about the phenotypes, ncORFs, refORFs, and corresponding loci. Data S12 also contains many additional fields, which will be described later in the text. The format of some fields, e.g. columns AC-BM, was adopted from our publication on 156 chimeric peptides (Supplementary Dataset S1 in Çakır et al., 2025b).

### Detection of ncORFs can improve the interpretation of loss-of-function phenotypes

The overall purpose of this analysis is to point out scenarios that are usually overlooked in functional genomics studies in any organism, not only in *M. truncatula*. We used our dataset of 673 genes for an unbiased assessment of ncORFs as a factor affecting the interpretation of functional studies. This analysis shows the extent to which robust experimental designs are free of interpretation ambiguity, even when a translated ncORF was overlooked. It also exposes the weak points of standard experimental procedures that cause such ambiguity.

### MS-validated ncORFs

It is meaningful to consider MS-validated and conserved ncORFs separately because of the difference in the level of their biological relevance. MS-supported ncORFs can reasonably be expected to have more relevance compared to ncORFs that have only conservation signatures. MS peptides detected as products of Tier 1-ncORFs are unique to those ncORFs. As follows from the analysis of their potential alternative sources using ChiMSource (Çakır et al., 2025a), these MS peptides cannot originate from any other locus in the *M. truncatula* genome, transcriptome, and repeatome in the course of conventional translation (Data S12, columns BN, BR, and BU). In the following few sections, ncORF9 was omitted from the narrative because it corresponds to the characterized mature peptide of *MtNCR169* (Starker et al., 2006; Domonkos et al., 2013; Farkas et al., 2014; Horváth et al., 2015, 2023; Güngör et al., 2023; Li et al., 2025). The detection of this peptide in our study can serve as a positive control for the validity of our approach. NcORFs 4, 6, 12, and 13 are described in the first place because of certain unique features. The remaining MS-validated ncORFs are presented in the ascending order of their chromosomal locations (Data S12).

**NcORF12 in *MthGSHSb***

Conceptually, methods based on post-transcriptional gene silencing, or PTGS (RNAi, miRNA, AS, and VIGS [Virus-Induced Gene Silencing]), cannot discriminate between the phenotypes of ncORFs and refORFs because they affect the entire transcripts hosting both ORFs (El-Sappah et al., 2021). Out of 11 genes with MS-validated ncORFs, only one was targeted by a PTGS tool, with no alternative methods used, *MthGSHSb* (Frendo et al., 2001, 2005). Downregulation of this gene using an antisense construct co-affected a related gene *MthGSHSa* (Frendo et al., 2001, 2005) and resulted in a conditional SNF phenotype (Data S6 and Data S12). Evidently, translated and conserved ncORF12 located in the 3'-UTR of *MthGSHSb* was co-affected, too. Thus, the reported nodulation phenotype can be attributed to the disruption of at least three translational units, or translons, according to the new terminology proposed recently by Świrski et al. (2025): refORFs of *MthGSHSa* and *MthGSHSb* along with ncORF12. NcORF12 is one of the prominent characters in our later discussion on the role of conserved ncORFs in functional genomics (Data S16).

**NcORF13 in *MtMYC2***

MS-validated ncORF13 is an example of the opposite type because its contribution to the reported phenotype is unlikely. It is nested in the coding sequence (CDS) of *MtMYC2*, which is a locus targeted by CRISPR/Cas9 in Guo et al. (2024). The study compared two mutant lines in which the ncORF was unaffected (*mtmyc2-1* and *mtmyc2-20*) with one line in which the ncORF was deleted (*mtmyc2-10*). There is no phenotypic difference among these three mutant lines, which suggests that the ncORF is dispensable for the processes studied by this group (Data S6 and Data S12). This example may seem rather fortuitous because the study was not specifically designed to target the refORF and the ncORF differentially. The ORFs were disrupted differentially by chance. However, it shows how important it is to consider multiple mutant lines and to refrain from PTGS as the only tool in a study. The case of ncORF13 is special because it is also free of ambiguity associated with the known effect of *Tnt1* on splicing (see further discussion on the limitations of *Tnt1* as a tool). At the same time, even this example lacks a valid control for the specific role of the overlapping translated ncORF because the only line in which the ncORF was deleted (*mtmyc2-10*) was simultaneously disrupted in the refORF, too. Thus, a dedicated study is warranted, in which the ncORF can be affected without changing the amino acid sequence of the refORF (see our recommendations on how this can be achieved in Supporting Discussion). The phenotype of ncORF13 may be different from that recorded in line *mtmyc2-10*, when considered independently from the refORF.

**NcORF6 in *MtTPST***

NcORF6 in locus *MtTPST* is a special case somewhat similar to the situation with ncORF12 (*MthGSHSb*). The *Tnt1* insertion NF18993_high_103 (FST retrieved from the *Tnt1* mutant database of *M. truncatula*; Tadege et al., 2008; Lee et al., 2018; Kaur et al., 2021) is located close to the 3'-terminus of this ncORF. This region corresponds to the 3'-UTR of locus MtrunA17_Chr4g0028941 (Data S16 Slide 1), according to the gene model of the latest genome assembly (v. 5.1.9). However, according to the gene model presented by the authors of the original study (Zhang et al., 2025), this ncORF is the end of the refORF. The exon-intron structure shown in Figure 3a of Zhang et al. (2025) is presumably based on the alignment of the cDNA sequence directly cloned from multiple samples rather than any predicted sequence. Thus, the original model is likely to be correct, which means that the ncORF is a misannotated part of the true refORF of this locus (Sequence 3 in Data S16 Slide 1). This model is supported by MS data (two MS peptides found specifically in 10-day-old nodules, Data S12, Data S16 Slide 1). Moreover, the ncProt and the refProt have high similarity to the same class of enzymes (protein-tyrosine sulfotransferase, Data S12, columns AU and AV). In addition, the taxonomic range of DIAMOND-based BLASTP hits of ncProt6 is broad (flowering plants). Together, these lines of evidence indicate that the refORF and ncORF constitute one continuous sequence. Alternatively, if the latest model is correct, the loss-of-function phenotype attributed to the refORF of *MtTPST* in Zhang et al. (2025) is caused by the disruption of the ncORF alone, without affecting the refORF of this gene. This ambiguity highlights the need for careful critical comparison of the new models with models reported in the original studies, like in the case of many other genes described earlier in the text. NcORF6 is another prominent example that will be discussed in the context of conservation analysis (Data S16).

**NcORF4 in *MtCKX6***

NcORF4, which is nested in the CDS of *MtCKX6*, is the only MS-validated ncORF in a gene that gave no altered phenotype when targeted by *Tnt1* (Wang et al., 2021). However, this does not mean that the ncORF is dispensable for the process under study (root development) because the *Tnt1* insertion in line NF14616 is located outside the ncORF. Thus, although the refORF of this gene is not involved in root development, the loss-of-function phenotype of the ncORF is yet to be determined.

**NcORF1 in *MtVPT2***

NcORF1 is located at the very beginning of the CDS in *MtVPT2*. The only *Tnt1* line studied by Liu et al. (2023) is inserted outside the ncORF. The same locus was targeted by RNAi in that study. Because the nodulation phenotypes of the *Tnt1* line NF0817 and the RNAi line are very similar, it may seem that the results rule out the involvement of the ncORF in the process under study (SNF). However, because the ncORF spans the border between the first two exons, the interpretation of this data is still ambiguous. In the section dedicated to the off-target effects of *Tnt1* on splicing (see Supporting Discussion), we will explain why the reported phenotypes of both *Tnt1* and RNAi lines of genes like *MtVPT2* may be equivocally attributed to the refORF and the ncORF. This ambiguity calls for a dedicated study in which the refORF and the ncORF can be disrupted differentially.

**NcORF2 in *MtREV1***

NcORF2, located in the first exon of *MtREV1*, is not affected by the *Tnt1* insertion in the study of Zhou et al. (2019). In contrast to *MtVPT2*, this gene was not targeted by RNAi. Thus, the reported phenotype can be attributed exclusively to the refProt of this locus. This is further supported by the fact that the off-target effect of *Tnt1* on splicing shown in Figure S3a of that study did not disrupt ncORF2. It only affected Exon 6. Elucidation of the potential role of ncORF2 requires a dedicated study that can alter the amino acid sequence of ncORF2 without affecting the refORF of *MtREV1*.

**NcORF5 in *MtWD40-1***

NcORF5 in the 3'-UTR of *MtWD40-1* is not affected by *Tnt1* insertions studies by Pang et al. (2009) and Meng et al. (2019b, 2023). Because ncORF5 does not coincide with an exon-exon border, it is unlikely that the *Tnt1* insertions co-affected this ncORF in trans. Thus, the reported phenotypes can be attributed exclusively to the refORF of this gene. This warrants a study in which the loss-of-function phenotype of ncORF5 can be assessed independently of the refORF.

**NcORF7 in *MtAHL1***

NcORF7 is nested in the CDS of *MtAHL1*. The gene was studied by CRISPR/Cas9 and RNAi (Zhang et al., 2023). While the ncORF was not affected in the CRISPR/Cas9 lines, the RNAi construct downregulated both the refORF and the ncORF because RNAi triggers systemic degradation of the entire transcript. The mutant phenotypes of the knockout and the knockdown lines shown in Figure 3 of Zhang et al. (2023) are very similar to each other, which suggests that the downregulation of ncORF7 may have added little if anything to the observed phenotype. Nevertheless, even in this situation, an unequivocal conclusion about the biological role of ncORF7 (or absence of such a role) can be made only after the ncORF is mutated independently from its refORF, that is, by an approach that leaves the refORF intact.

**NcORF8 in *MtCAS31***

NcORF8 spans the border between the CDS and the 3'-UTR of *MtCAS31*. This ncORF is not affected in TALEN lines analyzed in the study of Li et al. (2018). In contrast, the *Tnt1* mutant line NF5714 in the same study carries an insertion in ncORF8, which disrupts both the refORF and the ncORF. There is no strong difference between the phenotypes of the TALEN lines and the *Tnt1* mutant, which may suggest that ncORF8 is not essential for the biological processes studied by Li et al. (2018), namely SNF and drought stress. However, there are subtle differences between the microscopic images of TALEN lines and the *Tnt1* mutant discernible in Figures 6a and 6b of that study. These differences can potentially be attributed to ncORF8, which is unaffected in the TALEN lines. While the differences are minor and may have no biological relevance, the true loss-of-function phenotype of ncORF8 is unknown. To clarify the potential role of ncORF8 (or absence of such a role), the ncORF must be disrupted independently of its refORF.

**NcORF10 in *MtSCR***

NcORF10 is nested in the CDS of *MtSCR*. The *Tnt1* insertions studied by Dong et al. (2021) are located outside the ncORF. Thus, the true loss-of-function phenotype of ncORF10 is unknown. At the same time, because ncORF10 spans an exon-exon border, it is possible that the *Tnt1* lines are affected in both the refORF and the ncORF due to the effect of *Tnt1* on splicing, which can be exerted in trans, as will be demonstrated later (see Supporting Discussion). This ambiguity calls for a dedicated study, in which ncORF10 can be mutagenized independently from its refORF.

**Conservation analysis of ncORFs helps understand their evolutionary origins and refine annotated gene models**

The second group of ncORFs that deserves the attention of functional genomics specialists can be conditionally called conserved ncORFs. The term "conservation" that we use in this context is very broad as it refers to ncProts with at least one significant hit in the DIAMOND-based BLASTP similarity search using UniProt v. 2020_02 (UniProt Consortium, 2019), with an e-value at most 0.001. Naturally, this broad definition raises a valid concern about the biological relevance of conserved ncORFs. Specifically, sequence similarity analysis of ncORFs may seem to provide very little, if any, useful information for functional genomics studies in any organism, unless ncORFs are super-conserved. In this section and in the corresponding section of Supporting Discussion, we will show how important it is to consider conservation of ncORFs at any level, even when the taxonomic range of conservation signatures is very limited.

Before we introduce individual conserved ncORFs, we need to explain our methodology. One of the goals of our conservation analysis was to use the taxonomic range of subject sequences in the following three scenarios. In Scenario 1, a ncProt has significant hits in dozens or hundreds of eukaryotic species, mainly in the plant kingdom. Such a ncProt is either conserved and functional at the amino acid level or conserved and non-functional, without clear evidence of purifying selection. Thus, we searched specifically for ncProts conserved at the amino acid level stronger than at the nucleotide level. In Scenario 2, a ncProt has hits limited to a specific taxonomic group that does not include our model organism (*M. truncatula*, the family Fabaceae or Leguminosae, commonly known as the legume family). For example, a ncProt has hits exclusively in the fungal kingdom. Such a ncORF may look like a candidate for horizontal gene transfer (HGT). We have identified dozens of ncORFs potentially reflecting inter- and intra-kingdom HGT events in *M. truncatula*, some with truly interesting taxonomic ranges. However, closer analysis revealed that none of them originated via HGT. This observation indicates that the BLASTP similarity search alone is not sufficient for the detection of true HGT events. In Scenario 3, a ncProt has significant hits exclusively in *M. truncatula*. This scenario may seem to be of the least interest to functional studies because intra-species conservation is not thought to reflect the evolutionary pressure to preserve sequence function. In a dedicated section, we will explain why this scenario is important for a comprehensive functional annotation of a genome (see Supporting Discussion).

As a part of our effort to annotate ncORFs in *M. truncatula*, we deduced taxonomic ranges for all 18,643 ncProts reported in Supplementary Figure S1 of our study on chimeric peptides (Çakır et al., 2025b). 13,078 of them had top hits with at least 70% identity to annotated proteins from UniProt. The taxonomic range of each ncProt was based on the taxonomic status of source organisms of all hits (subject sequences). Many ncProts had more than 100 hits and some had thousands of hits (up to 153,028 hits). For the descriptive statistics on hits, we refer the readers to Supplementary Figures S2 and S3 in Çakır et al. (2025b). The complete list of conserved ncORFs and their corresponding taxonomic ranges will be published in a study dedicated to non-chimeric ncProts in *M. truncatula*. Among 673 loci studied by loss-of-function approaches in *M. truncatula*, 111 loci had at least one conserved ncORF, 132 ncORFs in total. Five more loci, with eight conserved ncORFs in total, were studied by gain-of-function approaches only. Taxonomic ranges of each ncORF in that list are shown in Column I of Data S12.

Next, we searched for ncORFs conserved stronger at the amino acid level compared to the nucleotide level. As a starting point for this search, we used a very simplified ("naïve") strategy, which permitted the transcriptome-wide coverage. Specifically, we compared the number of hits in the DIAMOND-based BLASTP searches (UniProt) with the number of hits in the BLASTN searches (NCBI) for each ncORF. The assumption was that a sequence with stronger conservation at the amino acid level should have more hits in the BLASTP search and the broader taxonomic range. Such an assumption is valid for some but not all sequences. Another obvious limitation of this simplified approach is the fact that the number of BLASTN hits depends on the type of database and on its scope/size. To partially alleviate this problem, we removed hits of the genomic DNA type, focusing exclusively on cDNA molecules. This comparison revealed thousands of ncORFs with more hits at the protein level compared to the nucleotide level, which will be published in a dedicated study. Among 140 conserved ncORFs shown in Data S12, only 28 fell into this category (ncORFs highlighted with cyan in Column F of Data S12).

The large-scale automated analysis described above (BLASTP/BLASTN hit ratio) was followed by a small-scale manual search for ncORFs conserved stronger at the amino acid level compared to the nucleotide level (percent identity difference and dN/dS ratio). For this analytical domain, we used the NCBI BLASTN tool, which guided us in the search for abundant synonymous differences in nucleotide sequences of ncORFs from *M. truncatula* and their best hits from other organisms. This analysis showed that only 11 out of 28 ncORFs shortlisted above were conserved stronger at the amino acid level (highlighted with cyan or green in Column E of Data S12). Among these 11 ncORFs, three corresponded to characterized NCR peptides MtNCR086 and MtNCR314 (Saifi et al., 2024), and MtNCR211 (Starker et al., 2006; Kim et al., 2015; Horváth et al., 2023), which proves that our strategy was efficient in finding biologically relevant ncORFs. It should be noted that ORFs of these peptides are different from the refORFs of corresponding loci annotated in the latest genome assembly (v. 5.1.9). Furthermore, refProts of these loci have no similarity to know proteins (MtrunA17_Chr4g0018034 for MtNCR211) or have similarity exclusively to unknown proteins in *M. truncatula* (MtrunA17_Chr3g0083311 for MtNCR086; MtrunA17_Chr3g0083301 for MtNCR314). This raises a concern about how refORFs are defined by the automated annotation pipeline of the *M. truncatula* genome, especially considering that all three loci are annotated as units producing the corresponding NCR peptides. Can these loci translate refProts other than NCR peptides? MtNCR211 is a special case; it corresponds to ncORF56 of locus MtrunA17_Chr4g0018031 (ncRNA) but is also a ncORF of locus MtrunA17_Chr4g0018034 (mRNA), which occupies the same genomic space. Besides these NCR-related ncORFs, there are eight ncORFs conserved stronger at the amino acid level. They are highlighted with cyan in Column E of Data S12. To the best of our knowledge, these ncORFs were not considered in the original studies on corresponding loci. We graphically highlighted the details of their conservation in Data S13. We also assessed non-synonymous and synonymous rates in nucleotide sequences of these ncORFs and calculated statistical significance of their ratios (dN/dS) under the hypothesis of purifying selection (Hurst, 2002). This information, generated using the distance and selection tools of the MEGA-X software v. 10.2.6 (Kumar et al., 2018), is also available in Data S13. The complete details of the dN/dS analysis with MEGA are shown in Data S14. Corresponding short alignments in MEGA format are available in Data S15. Data S16 visualizes alignments that help understand the evolutionary origins of conserved ncORFs. These alignments in MEGA and Geneious® formats are available in Data S17. Data S18 contains the complete set of gene models for all 123 genes listed in Data S12. The models are in Geneious® format, with annotated features that include refORFs, ncORFs, and exon-intron maps.

**NcORF6 in *MtTPST* calls for revision of the annotated gene model**

NcORF6 has already been discussed in the section about MS-validated ncORFs, where we have explained why this ncORF is likely to be the continuation of the *MtTPST* refORF. Here, we will focus on its conservation and putative origin. NcORF6 was detected in the 3'-UTR of a protein-tyrosine sulfotransferase-encoding gene involved in SNF, root development, and many other processes (Zhang et al., 2025), which are summarized in Data S6. *MtTPST* has no homologs in *M. truncatula* (Data S16, Slide 1). NcORF6 is not listed in Data S13 because it is not under purifying selection. Nevertheless, it is homologous to the 3'-portions of protein-tyrosine sulfotransferase refORFs of at least five other legume species (Data S16, Slide 2). In the current gene model of MtrunA17_Chr4g0028941, it is disconnected from the refORF through an early in-frame stop codon introduced by a putative insertion of 915 bp (Data S16, Slide 3), which is present in a slightly modified form in *M. arabica* and *M. lupulina* (93 and 94% identity, respectively) but has limited query coverage (9-70%) and similarity (77-92%) to other sequences in GenBank (as of 25 February 2026). This insertion is flanked by a sequence AGGC (Data S16, Slides 3 and 4), which is very similar to the classical exon-intron and intron-exon border motifs (Mount, 1982; Shapiro & Senapathy, 1987). After the removal of this insertion precisely at the putative splicing sites, the ncORF becomes a smooth continuation of the refORF and is seamlessly aligned with refORFs of the other five legumes (Data S16, Slide 5). This indicates that the insertion is likely to act as a retained intron in the actual gene model. Consequently, there must be an alternative splicing form in which this intron is removed. Such a form is supported by MS data (two peptides). The absence of evidence for purifying selection in this region is surprising, given the evidence for translation and homology to conserved refProts. It suggests that the amino acid sequence of ncProt6 is repurposed compared to the C-terminal refProt portions of *MtTPST* homologs from other species.

**NcORF11 and ncORF12 in *MthGSHSb* call for revision of the annotated gene model**

NcORF12 has already been discussed above because it is one of the two ncORFs that are MS-validated and conserved at the same time. Here, we will focus on its conservation and origin, together with the second conserved ncORF in *MthGSHSb*, ncORF11. This gene was downregulated by an antisense construct along with its homolog *MtGSHSb* in the study by Frendo et al. (2005), which resulted in a conditional nodulation phenotype (Data S6). *MthGSHSb* and *MtGSHSb* encode homo-glutathione (hGSH) and glutathione (GSH) synthases, respectively (“h” stands for “homo”). In their earlier work (Frendo et al., 2001), the authors deduced the origin of the homo-glutathione synthase gene from the duplication of the glutathione synthase gene. However, at the time of the publication, the genome annotation of *M. truncatula* was in its infancy. Thus, the group was unaware of four homologs of this type existing in *M. truncatula*: MtrunA17_Chr7g0273151 (*MtGSHSa*), MtrunA17_Chr7g0273131 (*MthGSHSa*), MtrunA17_Chr7g0273161 (*MtGSHSb*), and MtrunA17_Chr7g0273141 (*MthGSHSb*). In the previous version of the *M. truncatula* genome (v. 4), *MtGSHSa* and *MtGSHSb* were combined in one locus, Medtr7g113890, which corresponds to the long form called *MtGSHS1* (glutathione synthase) in Frendo et al. (2001). In contrast, locus MtrunA17_Chr7g0273141 (*MthGSHSb*) was unknown in the *M. truncatula* genome (v. 4). Together with locus MtrunA17_Chr7g0273131 (*MthGSHSa*), it corresponds to the short form referred to as homo-glutathione synthase *MtGSHS2* in Frendo et al. (2001). Several putative homologs/sequence variants of these genes can be found in GenBank (Data S16 Slide 6). Some of them probably originate from the same locus with *MthGSHSb* (Data S16 Slide 7). Others originate from the glutathione synthase locus *MtGSHSb* or may represent products of unknown loci (Data S16 Slide 8). Thus, there is much uncertainty about the actual number of homo-glutathione (hGSH) and glutathione (GSH) synthase homologs in *M. truncatula*. Moreover, it is not easy to discriminate between sequences that encode homo-glutathione synthases and those encoding glutathione synthases, none of which are annotated as such in the current genome assembly. We made the first step toward better understanding of these sequences by aligning them with each other and with all similar sequences available in GenBank (Data S16 Slide 6). Based on this analysis, we named MtrunA17_Chr7g0273151 and MtrunA17_Chr7g0273131 *MtGSHSa* and *MthGSHSa*, respectively, for the following reasons. They correspond to the left portion of the long form called *MtGSHS1* (glutathione synthase), as shown in Data S16 Slide 9. Thus, we added the “a” at the end of each name to discriminate these sequences from those that correspond to the right portion of *MtGSHS1* (“a” for left, “b” for right). However, only MtrunA17_Chr7g0273151 (*MtGSHSa*) groups with *MtGSHS1* (glutathione synthase, see Data S16 Slide 6). Thus, the prefix “h” is missing from its name ("h" stands for "homo"). By contrast, MtrunA17_Chr7g0273131 (*MthGSHSa*) groups with the short form referred to as homo-glutathione synthase *MtGSHS2*. Thus, we added the prefix "h" to its name. Likewise, the phylogenetic tree and the alignment shown in Data S16 Slides 6 and 9, respectively, were the basis for naming loci MtrunA17_Chr7g0273161 (*MtGSHSb*) and MtrunA17_Chr7g0273141 (*MthGSHSb*). Specifically, they correspond to the right portion of the long form called *MtGSHS1* (glutathione synthase); hence the "b" added to their names ("a" for left, "b" for right). While MtrunA17_Chr7g0273161 (*MtGSHSb*) groups with *MtGSHS1* (glutathione synthase), MtrunA17_Chr7g0273141 (*MthGSHSb*) is found in the same clade with *MtGSHS2* (homo-glutathione synthase); hence the prefix "h" in *MthGSHSb* ("h" stands for "homo").

NcORF11 is located at the border between the 5'-UTR and the CDS of *MthGSHSb* (Data S16 Slide 7). It has a relatively narrow taxonomic range of DIAMOND-based BLASTP hits (flowering plants). However, it possesses a highly conserved region of 66 nt that is under purifying selection (Data S13 Slides 1-3). The synonymous rate (dS) is significantly higher than the non-synonymous rate (dN) for this region in the alignments with the homoglutathione synthetase cDNA from *Phaseolus vulgaris* (eudicot, legume), the glutathione synthetase cDNA from *Salix viminalis* (eudicot, non-legume), and the chloroplastic glutathione synthetase cDNA from *Populus trichocarpa* (eudicot, non-legume) (Data S13 Slides 1-3). NcProt11 aligns with the refProts of *MtGSHS1* and *MtGSHS2* (Data S16 Slide 10). As follows from Data S16 Slides 11 and 12, ncORF11 may have evolved from a recent intron retention event, which preserved the first four nucleotides of Intron 1 in the cDNA (TATG). Alternatively, ncORF11 may be the start of the refORF in a splicing form missed by the current genome annotation.

NcORF12 resides in the 3'-UTR of *MthGSHSb* (Data S16 Slide 7). In contrast to ncORF11, it has an unusually broad taxonomic range of BLASTP hits: vascular plants, bryophytes, Charophyta (Chara-like algae), green algae, other protists, fungi, and animals (Data S12). However, the purifying selection signature of ncORF12 is weaker (Data S13 Slides 4 and 5). The synonymous rate (dS) is significantly higher than the non-synonymous rate (dN) in the alignment with the chloroplastic glutathione synthetase cDNA from *Cicer arietinum* (eudicot, legume) (Data S13 Slide 4). In the alignment with the glutathione synthetase cDNA from *Cucumis sativus* (eudicot, non-legume), the dN/dS ratio and its significance are not computable with the MEGA software even though the identity value at the amino acid level (71%) is higher than at the nucleotide level (67%) (Data S13 Slide 5). The most probable cause of the noncomputability is the combination of short alignment length and high divergence between a legume and non-legume. NcORF12 corresponds to the end of refORFs of several other sequences, including *MtGSHS1* and *MtGSHS2* from Frendo et al. (2001), and *MtGSHSb* (Data S16 Slides 7, 13, and 14). It was disconnected from the refORF by a premature in-frame stop codon introduced via an insertion of 85 bp, which has only a few hits in the BLASTN search using the NCBI non-redundant nucleotide database (as of 25 February 2026, Data S16 Slide 14). It is possible that this insertion acts as a retained intron in the current gene model of *MthGSHSb* because of two sites very similar to the classical exon-intron and intron-exon borders (AGGT and AGGA, respectively, in Data S16 Slide 15) (Mount, 1982; Shapiro & Senapathy, 1987). When these 85 nucleotides are removed precisely at the putative splicing sites (results not shown), the cDNA of *MthGSHSb* aligns perfectly with many other sequences listed in Data S16 Slide 15. As indicated earlier, ncORF12 is translated. This follows from our detection of two MS peptides, VIVNNESGYMVR and DKVIVNNESGYMVR (Data S12), in six proteomic samples of three independent studies (PXD002692, Marx et al., 2016; PXD013606, Shin et al., 2021; PXD022278, Castañeda et al., 2021). These lines of evidence strongly suggest that the latest genome annotation missed a splicing variant lacking this putative retained intron. Moreover, it is possible that *M. truncatula* has a long form that combines the upstream short "a" sequence (like *MtGSHSa* and *MthGSHSa*) with the downstream short “b” sequence (like *MtGSHSb* and *MthGSHSb*) in one unit similar to *MtGSHS1*, AC172101.1, and AF075700.2 (Data S16 Slides 9 and 13). This possibility is supported by the fact that such long forms are ubiquitous among eudicots (results not shown).

**NcORFs 51, 53, 54, and 55 correspond to modified portions of an ancestral, longer ORF, which suggests that *MtPHO2-A* is a functional unitary pseudogene**

*MtPHO2-A* is involved in the regulation of phosphate homeostasis, nodulation, and vegetative growth (Curtin et al., 2017; Čermák et al., 2017; Huertas et al., 2023), as follows from the reduced biomass and overaccumulation of phosphate in roots and old leaves of *mtpho2-a* mutants inoculated with rhizobia (Data S6). *MtPHO2-A* has five ncORFs with the ratio of BLASTP/BLASTN hits above one (Data S12). Four of them are under purifying selection as indicated by the statistical significance of dN/dS ratios (Data S13 Slides 6-18). The taxonomic ranges of BLASTP hits are relatively narrow for ncORFs 51, 54, and 55, but are extremely broad for ncORF52 and especially ncORF53. The latter one has significant hits in vascular plants, bryophytes, Charophyta (Chara-like algae), green algae, red algae, other protists, fungi, animals, archaea, eubacteria, and viruses. NcORFs 51 and 52 reside in the 5’-UTR. NcORF52 partially overlaps with the refORF of this gene. NcORFs 53, 54, and 55 are in the 3’-UTR (Data S16 Slide 16). NcORF52 is out of frame relative to the refORFs of two homologs of *MtPHO2-A* (Data S16 Slides 17 and 18). The remaining four ncORFs largely preserved the original amino acid sequence of the ancestral, long refProts of *MtPHO2-B* and *MtPHO2-C* (Data S16 Slides 17, 21, 22, 24, and 25). The current refORF length of the *MtPHO2-A* cDNA (945 bp) is about one third of the refORFs of its two *M. truncatula* homologs: 2,772 bp in *MtPHO2-B* and 2,742 bp in *MtPHO2*-C. This analysis indicates that *MtPHO2-A* can be categorized as a functional unitary pseudogene according to the definition of Cheetham et al. (2020). The considerable shortening of the refORF did not deprive this gene of its important function and endowed four ncORFs with protein-coding potential, as detected in our study.

Details of the alignment shown in Data S16 Slide 16 help elucidate the putative evolutionary events that led to the current gene model of *MtPHO2-A* (Data S16 Slides 17-25). The original reading frame of the hypothetical ancestor was shifted immediately downstream of the conserved region of ncORF51 via the loss of a single nucleotide (presumably A). This event brought ncORF52 into a frame that is alternative to refORFs of *MtPHO2-B* and *MtPHO2-C* (Data S16 Slide 17). Nevertheless, ncProt52 has a very broad taxonomic range of DIAMOND-based BLASTP hits, as far from *M. truncatula* as fungi and animals. Many of these hits correspond to proteins of the same functional group as MtPHO2-A (ubiquitin-conjugating enzyme, Data S12). This similarity suggests that diverse taxa explored this alternative reading frame and made it part of the refORFs of their *MtPHO2-A* homologs (Data S16 Slide 16). The ancestral reading frame was regained by the modern version of *MtPHO2-A* via the deletion of multiple nucleotides at the end of ncORF52 (Data S16 Slide 18). However, the refORF was prematurely terminated after only 945 bp by a single-nucleotide deletion (presumably T), which caused a frameshift and introduced an early in-frame stop codon (Data S16 Slide 19). The downstream cDNA sequence until the beginning of ncORF53 is not present in the cDNA of *MtPHO2-B* and *MtPHO2-C*. Most likely, it originates from an intron retention event because it is found in the genomic DNA of all three genes (Data S16 Slide 20). This possibility is further supported by the presence of two AGGT sites precisely at the beginning and at the end of this putative retained intron (Data S16 Slides 19 and 21). The sequence AGGT is very similar to the classical exon-intron and intron-exon borders (Mount, 1982; Shapiro & Senapathy, 1987). This intron retention event terminated the refORF of *MtPHO2-A* but brought the downstream ncORF53 back to the same frame with refORFs of the other genes. Thus, ncProt53 is highly similar to refProts of ubiquitin-conjugating enzyme genes in many diverse lineages (Data S12).

Likewise, the cDNA segment between the strongly conserved regions of ncORFs 53 and 54 is likely to originate from an incomplete intron retention event (Data S16 Slide 22). This sequence is partially present in the genomic DNA of *MtPHO2-B* but is lost or heavily modified in the genomic DNA of *MtPHO2-C* (Data S16 Slide 23). This putative retained intron is flanked by sequences AGGT and AGCA (Data S16 Slides 22 and 24), which resemble classical motifs of exon-intron and intron-exon borders (Mount, 1982; Shapiro & Senapathy, 1987). Because of this incomplete intron retention, the conserved region of ncORF54 is out of frame with the refORFs of *MtPHO2-B* and *MtPHO2-C*. However, it is nearly identical to the refProts at the amino acid level (Data S16 Slide 24).

NcORF55 terminates exactly at the same position as the refORFs of *MtPHO2-B* and *MtPHO2-C*. NcORF55 regained the same reading frame as the refORFs because of a two-nucleotide insertion at the beginning (Data S16 Slide 25). This explains why ncProt55 is highly similar to ubiquitin-conjugating enzymes from seed plants (Data S12).

The purifying selection signatures of ncORFs 51, 53, 54, and 55 (Data S13 Slides 6-18) may reflect their very recent origin. It is conceivable that parts of a conserved refORF can retain their sequences for some time after pseudogenization. If this is true for *MtPHO2-A*, it represents a remarkable case of a rapid pseudogenization, where only a portion of the original refORF is sufficient for an important biological role. Alternatively, these ncORFs may be translated as discrete units or as part of a longer refORF missed in the current genome annotation. The current gene model of *MtPHO2-A* may simply correspond to an incompletely spliced form. If these ncORFs are translated, it is important to know which of them could have contributed to the mutant phenotype attributed to the refProt of *MtPHO2-A*.

The current refORF definition and the annotated exon-intron structure of *MtPHO2-A* are different from those reported in the functional studies. Only one or two out of six highly conserved *PHO2*-like ORFs in locus MtrunA17_Chr4g0009054 have been targeted by Curtin et al. (2017), Čermák et al. (2017), and Huertas et al. (2023). According to the current model (v. 5.1.9), CRISPR/Cas9 and TALEN lines affected only ncORF52 located at the end of the 5'-UTR. It was assumed to be part of the refORF in all three studies. The *Tnt1* line NF12360 contains an insertion that cannot be accurately located in MtrunA17_Chr4g0009054 because FSTs NF12360_high_9 and NF12360_low_9 contain no *Tnt1* end signature that is normally used for deducing the exact position. The *Tnt1* insertion may have affected either ncORF52 alone or together with the upstream ncORF51 (both reside in the 5'-UTR; ncORF52 partially overlaps with the refORF). This is one of a few remarkable cases in which the mutant phenotype cannot formally be linked to the currently annotated refORF unless it is affected by the *Tnt1* insertion via its interference with normal splicing (see Supporting Discussion). Sequence features of *MtPHO2-A* serve as an excellent example of ambiguity introduced by ncORFs as a hidden dimension of the protein-coding capacity. This gene helps illustrate how conservation analysis of ncORFs can improve the accuracy of gene models and the interpretation of loss-of-function studies.

**NcORF60 in *MtbZIP60b* is a conserved remnant of a longer refORF**

Transcription factor MtbZIP60b is involved in the control of leaf senescence (Xing et al., 2024). It has at least four related loci in the current genome annotation of *M. truncatula* (Data S16 Slide 26) and additional unannotated sequences in GenBank (Data S16 Slide 27), which may be sequence variants of *MtbZIP60b*. NcORF60 resides in the 5'-UTR of *MtbZIP60b* (Data S16 Slide 27). Although it has a modest taxonomic range of DIAMOND-based BLASTP hits (eudicots and monocots, Data S12), it is under purifying selection (Data S13 Slides 19-21). The conserved region of ncORF60 corresponds to the beginning of refORFs in the alignment with two *M. truncatula* homologs, MtrunA17_Chr3g0141881 and MtrunA17_Chr3g0144931

(Data S16 Slide 28). Although short sequences MtrunA17_Chr3g0144931 and MtrunA17_Chr3g0144941 are currently annotated as two different loci (v. 5.1.9), they probably correspond to two portions of a single genetic unit similar to MtrunA17_Chr3g0141881. This possibility is supported by their adjacent location in the genome, where they are separated by a single nucleotide C (Data S16 Slide 29). NcProt60 is highly similar to refProts of MtrunA17_Chr3g0141881 and MtrunA17_Chr3g0144931 (Data S16 Slide 30).

In the alignment of the *MtbZIP60b* cDNA with homologs from five other legume species, the conserved segment of ncORF60 corresponds to the 5'-portion of refORFs that are similar in structure to MtrunA17_Chr3g0141881 (Data S16 Slide 31, sequences 1-3). Specifically, these refORFs combine regions corresponding to the ncORF and the refORF of *MtbZIP60b* in one continuous sequence in *Lupinus albus*, *Glycine max*, and *Mucuna pruriens* (Data S16 Slide 31). However, in *Pisum sativum* and *Vicia villosa*, the refORFs begin at the same place as in the annotated version of *MtbZIP60b*, which supports the current model of this gene. A sequence separating the strongly conserved ncORF region and the refORF of *MtbZIP60b* introduces an early in-frame stop codon, which terminates ncORF60 (Data S16 Slide 32). This separating sequence is not found in any annotated homolog in *M. truncatula* (Data S16 Slide 28, sequences 1-4). This segment is unlikely to correspond to a retained intron because it is missing from the genomic DNA of the long homolog MtrunA17_Chr3g0141881 (Data S16 Slide 33). The sequence is shared by legumes from five genera, specifically *Medicago*, *Pisum*, *Vicia*, *Trifolium*, and *Lathyrus*, and is not a remnant of a transposon (Data S16 Slides 33 and 34).

In the *Tnt1* mutant line studied by Xing et al. (2024), the insertion is outside ncORF60. The ncORF is unlikely to be affected by *Tnt1* in trans because ncORF60 does not span an exon-exon border. The separation of ncORF60 from the refORF is not recent because this region is conserved at the amino acid level in five diverse legume species. Thus, the purifying selection signature of ncORF60 may point to an important role, which remains to be demonstrated. NcORF60 may be translated either as a discrete unit or as an exon of the longer refORF similar to that of MtrunA17_Chr3g0141881.

**Super-conserved ncORF65 in *MtKASII* is an artifact or an evolutionary paradox**

MtKASII is a ketoacyl-[acyl-carrier-protein] synthase involved in AM symbiosis. It was studied by RNAi (Jiang et al., 2017). MtKASII has many annotated homologs in *M. truncatula* and some related sequences in GenBank (Data S16 Slide 35). NcORF65 is nested in the CDS of this gene (Data S12). Both MtKASII and ncProt65 are the most similar to the refProt of MtrunA17_Chr1g0204841. However, ncProt65 also shares much similarity with refORFs of other *M. truncatula* homologs (Data S16 Slide 36). NcORF65 has several unique features. The taxonomic range of DIAMOND-based BLASTP hits of ncProt65 is unusually broad: vascular plants, bryophytes, Charophyta (Chara-like algae), green algae, red algae, other protists, fungi, animals, archaea, and eubacteria (Data S12). It is under purifying selection, as follows from dN rates significantly lower than dS rates in alignments with homologs from *Glycine soja* (eudicot, legume), *Cephalotus follicularis* (eudicot, non-legume), *Rhynchospora pubera* (monocot), and *Magnolia sinica* (magnoliid) (Data S13 Slides 22-25). In the alignments of ncORF65 with homologs from *Physcomitrium patens* (bryophyte) and a bacterial species from *Pseudomonadota*, the dN/dS ratios and their statistical significance are not computable with the MEGA software, probably because of the combination of short alignment length and high divergence between taxa. However, in these alignments, the conservation at the amino acid level is much stronger than at the nucleotide level. For the bryophyte, the difference is 22% (95%-73%), and for the bacterium, it is 24% (95%-68%), which is the largest difference observed in the entire dataset (Data S13 Slides 26 and 27, Data S14).

The putative evolutionary origin of ncORF65 is remarkable. The strongly conserved part of the ancestral refORF was frameshifted by the deletion of one nucleotide immediately upstream of the highly conserved region (Data S16 Slide 37). After only 117 bp, the reading frame was brought back to the refORF by the deletion of 11 bp immediately downstream of the highly conserved region. This is an example of a "short round trip" mosaic protein (Çakır et al., 2023) created at the genomic DNA level without programmed ribosomal frameshifting. Specifically, a short segment from the alternative frame was integrated into the strongly conserved canonical protein, which was tolerated by natural selection. While the deletion of a single G or T is an intra-exonic event that brought part of the refORF into the alternative frame, the downstream frameshift was caused by skipping the first 11 nucleotides of Exon 3 (GTGATGGAAAG) (Data S16 Slide 38). These nucleotides are still found as a part of CDS in *M. truncatula* sequences XP_003608492.2 and AES90689.1 (Data S16 Slide 37).

Before accepting the current frameshifted gene model of *MtKASII* as correct, it is necessary to consider evidence against its existence. There are three GenBank entries that group together with *MtKASII* in the phylogenetic tree shown in Data S16 Slide 35: XP_003608492.2, AES90689.1, and AFK43304.1. Apart from the very few differences visible in the alignment upstream and downstream of the strongly conserved region of the ncORF, all four sequences are identical over the entire length (Data S16 Slide 39). This suggests their origin from a single genetic locus. If the current gene model of *MtKASII* is correct, it must have at least one non-frameshifted sequence variant similar to the gene model of MtrunA17_Chr1g0204841 (Data S16 Slide 37). Alternatively, the current frameshifted model of *MtKASII* is a sequencing artifact. Validation of this unusual model requires detection of genetically frameshifted MS peptides and verification of the corresponding transcript sequence using RNA-Seq.

NcProt65 is highly similar to refProts from diverse plant lineages and even a bacterium (Data S16 Slide 40). Among these proteins, only MtKASII contains a segment translated from the reading frame that is alternative to refORFs of all other sequences. However, there are three exceptions in GenBank: one MtKASII homolog from *Brassica napus* and two unrelated proteins from the genus *Ipomoea* (Data S16 Slide 41) use the region homologous to ncORF65 as a part of their refORFs. The ncProt65-like sequence evolved independently in *B. napus* via a simpler frameshifting mutation, as follows from no return to the original frame and early termination of the sequence in this species. Two *Ipomoea* sp. sequences are annotated as unrelated proteins: bidirectional sugar transporter SWEET1-like (*I. trifida*) and ribonucleoside-diphosphate reductase large subunit-like (*I. batatas*). They have very low overall percent identity with MtKASII (14% and 11%, respectively). The extreme rarity of using ncORF65-like sequences as refORF parts in the tree of life highlights this ncORF either as an evolutionary paradox or an artifact. If a short refORF region homologous to ncORF65 is conserved across so many taxa, how can these two frameshifting mutations be tolerated by natural selection? On one hand, there is no MS proteomic evidence for translation of this ncORF in 16 different samples of three independent studies (PXD002692, Marx et al., 2016; PXD013606, Shin et al., 2021; PXD022278, Castañeda et al., 2021). On the other hand, the currently annotated mosaic refProt of *MtKASII* is not MS-validated either. While there are numerous MS peptides that match the refProt of *MtKASII*, none of them match the region translated from the alternative frame (Data S16 Slide 42). Without the conservation analysis of ncORF65, the current model would remain unchallenged. With this analysis, the model stands out and calls for closer attention.
