## Supplementary material for "Loss-of-function phenomics, ncORFs, and ambiguity of mutant phenotypes in *Medicago truncatula*": Table S1

**Table S1**. Loci incorrectly named in the genome browser v. 5.1.9 (an extended version of Table 3)

| Name† | Actual locus ID (publication) | Publication | Misnamed locus ID (v. 5.1.9) | Correct name and publication for a misnamed locus |
|---|---|---|---|---|
| MtCbf4 | MtrunA17_Chr1g0203861 | Zhang et al., 2016b | MtrunA17_Chr1g0195851 | MtNF-YB6 (Baudin et al., 2015) |
| MtChOMT1‡ | MtrunA17_Chr1g0200491 | Wu et al., 2024 | MtrunA17_Chr3g0085211,<br>MtrunA17_Chr7g0218131 | MtOMT2 (Wu et al., 2024),<br>MtChOMT3 (Wu et al., 2024) |
| MtGA20ox1 | MtrunA17_Chr1g0204181 | Li et al., 2021b | MtrunA17_Chr6g0474391 | MtGA20ox7 (Li et al., 2021b) |
| MtHDT3 | MtrunA17_Chr7g0265761 | Li et al., 2021a | MtrunA17_Chr4g0027181 | MtHDT1 (Li et al., 2021a) |
| MtMYC2 | MtrunA17_Chr8g0366751 | Guo et al., 2024 | MtrunA17_Chr5g0411341 | MtBHLH2 (Deng et al., 2019) |
| MtPRP1‡ | MtrunA17_Chr4g0013391 | Erickson et al., 2020 | MtrunA17_Chr4g0013371,<br>MtrunA17_Chr4g0013401 | MtPRP2 (Erickson et al., 2020),<br>MtPRP3 (Erickson et al., 2020) |
| MtPRR7 | MtrunA17_Chr1g0181811 | Wang et al., 2023g | MtrunA17_Chr8g0345901 | MtPRR9b (Wang et al., 2023g) |
| MtRac1 | MtrunA17_Chr6g0486041 | Wang et al., 2021d | MtrunA17_Chr4g0046211 | MtROP7a (Wang et al., 2021d) |
| MtROP9 | MtrunA17_Chr5g0405791 | Kiirika et al., 2012 | MtrunA17_Chr6g0486041 | MtRac1 (Wang et al., 2021d) |
| MtSCR | MtrunA17_Chr7g0245601 | Dong et al., 2021 | MtrunA17_Chr5g0396491 | MtSCL3 (Seemann et al., 2022) |
| MtSLB1 | MtrunA17_Chr5g0447381 | Yin et al., 2020; Zhou et al., 2021 | MtrunA17_Chr5g0447371 | NA |

†The table lists correct names and corresponding locus identifiers. These names were incorrectly assigned to other loci in the genome browser v. 5.1.9, which are shown in column 4.
‡Names MtChOMT1 and MtPRP1 were misassigned to two genes each in the genome browser v. 5.1.9.
References in this table have letter codes (a, b, c, etc.) according to Data S7, which may be different from the letter codes in the main text.
